# The first assembled chromosome-level genome of an individual from Karnataka, India

**DOI:** 10.1101/2024.08.18.608448

**Authors:** Anisha Mhatre, Yash Chindarkar, Apoorva Ganesh, Aditi Padmakar Thakare, Prakruti Mishra, Aditya Sharma, Febina Ravindran, Subhashini Srinivasan, Bibha Choudhary

## Abstract

We present the first chromosome-level assembled genome of an individual from Karnataka, India (KIn1), using de novo assembly of 12x coverage of PacBio HiFi reads generated in-house and 30x Hi-C data from the most ethnically related individual to KIn1 from Andhra Pradesh (HG04712), publicly available under International Genome Sample Resource (IGSR). We achieved an N50 of 137 Mb and an L50 of 9—very close to the maximum achievable N50 of 147 Mb and minimum achievable L50 of 8 respectively for human genomes.

A comparison of the KIn1 assembly with assemblies of four other individual genomes from a Puerto Rican (PR1), Ashkenazi (Ash1), Hans Chinese (Hans1), and Northern European (T2T) reported to date from resources like IGSR and/or Genome-in-a-Bottle (GIBR) collections shows that the longest scaffolds from KIn1 cover 80-90% of the reference chromosomes and exhibit near collinearity. We find a unique inversion in the chromosome 7 euchromatic region and other major translocations in chromosomes 17 and 20 in KIn1. We also produced a chromosome-level assembly for another individual from Punjab, Lahore (HG03492), for use in our comparison. The analysis of variants among select IGSR individuals using the KIn1 assembly as a reference reveals that the Indian Telugu from the UK (ITU) population in IGSR is ethnically closest to KIn1, followed by the Sri Lankan Tamil from the UK (STU) as representatives of South Asian ancestry.

## INTRODUCTION

The order of the 3 billion base pairs in the human genome was first deciphered from a pool of 5 individuals and was reported in 2001^1^. This momentous achievement led to envisioning a day when the majority of individuals’ genomes could be deciphered for as little as $1000. With substantial funding to fulfill this vision, sequencing technologies now exist that allow sequencing of individual human genomes for tens of thousands of US dollars.

It took two decades to close numerous gaps in the genome published in 2001^2^ known as GRCh38. The improved version of the first human genome is a mosaic of several individuals and still retains hundreds of gaps due to heterogeneity among the individuals pooled to create the reference. These incomplete regions include large gaps at all centromeres and the short arms of acrocentric chromosomes.

Advances in second-generation long-read sequencing technologies have generated a gapless assembly, known as Telomere-to-Telomere (T2T), from an individual of Northern European descent, establishing it as a new reference against which other individual genomes can now be compared^3^.

As of today, assemblies of genomes from a Puerto Rican individual(PR1)^4^, a Han Chinese individual (Han1)^5^, and an Ashkenazi Jew individual (Ash1)^6^ have been reported, compared, and annotated against the Telomere-to-Telomere (T2T) reference. The PR1 genome assembly of an individual with six out of eight great-grandparents born in Puerto Rico (HG01243 from the NHGRI collection) was generated utilizing PacBio HiFi (34X), Illumina (59X), and ONT long reads (35X). The assembly using ONT reads with Flye provided the best N50 and served as the foundation for merging with other assemblies. This was further integrated with a MaSuRCA assembly generated from Illumina, ONT, and PacBio-CLR reads, as MaSuRCA showed the best consensus with Illumina variants. Scaffold orientation was performed using the T2T-CHM13 assembly. A total of 147 gaps in the Flye assembly were filled by merging with MaSuRCA contigs, and the remaining gaps were resolved using the T2T-CHM13 assembly. The PR1 assembly utilized 280 Mbp from the T2T-CHM13 assembly to fill the gaps. Validation of the PR1 assembly employed Mercury, PacBio HiFi reads, and Hi-C data from the same individual.

The assembled genome of Ash1 utilized HG002, an Ashkenazi individual who is part of the Personal Genome Project (PGP)^7^. Publicly available raw data was employed for this assembly^8^, including 71x Illumina, 23x ONT, and 29x PacBio HiFi sequencing coverage. Two assemblies were generated: one using Illumina and ONT data, and another integrating all three datasets. Integration of PacBio HiFi data resulted in a larger total assembly size and a higher N50 (18.2 Mb vs. 4.9 Mb), which was selected for further refinement. The scaffolds from this assembly were aligned to GRCh38 to fill gaps, incorporating 58.3 million bases from GRCh38. The resulting gaps in Ash1 totaled 82.9 Mbp, compared to 84.7 Mbp in GRCh39.p13. Validation of the assembly included comparison with variants from the same sample in the database, providing a significant advantage. Additionally, the report indicated approximately 1 million fewer homozygous SNPs compared to an unrelated Ashkenazi individual when aligned against GRCh38.

The genome data reported for a Southern Hans Chinese male (Han1) (HG00621) were generated and publicly released by the Human Pangenome Reference Consortium (https://github.com/human-pangenomics/hpgp-data). The initial assembly of Han1 was constructed using 39x PacBio high-fidelity (HiFi) reads and 35x Oxford Nanopore Technology (ONT) reads. Assembly of all HiFi reads with Hifiasm and all ONT reads with Flye resulted in a HiFi draft assembly with improved contiguity, boasting an N50 contig size of 95.8 Mb and comprising only 182 contigs, compared to an N50 of 40.9 Mb and 1,658 contigs from Flye. Contig orientation was performed using MaSuRCA with T2T-CHM13 as a reference, like PR1 and Ash1 assemblies described previously. This step identified 12 misassembled contigs that required manual splitting. Additionally, there were 77 unplaced contigs totaling approximately 39 million bases. The objective was to close all remaining gaps by integrating other assemblies with corresponding sequences from the T2T assembly.

In 2023, the first draft of the human pangenome was published using data from emerging sequencing technologies generated as part of IGSR samples^9^. This work capitalized on the ethnic diversity within the IGSR sample collection to achieve phased assembly of 47 ethnically diverse individuals. PacBio HiFi data was generated at a coverage of approximately 40x, complemented by Illumina reads from both parents. Although many samples included ONT, Illumina, and Bionano reads publicly available through IGSR, these were not utilized in the report. The authors employed the trio-HiFiasm tool, which utilizes short-read Illumina data from both parents to produce phased contigs from assemblies generated by PacBio HiFi data. These assemblies remain at the contig level and have not yet been scaffolded to achieve chromosome-level representations, which are essential for genome-wide comparisons. The authors acknowledge 217 inter-chromosomal joins, with only one occurring in the euchromatic region. These mis-assemblies were manually corrected.

In 2021, an effort to optimize human genome assembly demonstrated that PacBio HiFi reads of 30x coverage along with HiC reads are sufficient to achieve a chromosome-level phased assembly^10^ without the need for pedigree data and raw reads from other technologies. Despite large representation of individuals from South Asian ancestry in IGSR, a chromosome-level reference assembly from this region is missing, challenging the study of diversity within Indian population. Our in-house efforts to study diversity within this population using population specific variation against hg38, a reference distant to South Asian population, was unable to segregate individuals by ethnicity (data not shown).

Here, we report the first genome of an individual of Indian descent from the state of Karnataka (KGP) not part of any public consortia.

## RESULTS

### Optimization of coverage requirement

The individual genomes reported so far has used at least 30-40x coverage of PacBio HiFi data along with Illumina and ONT technologies. Our goal here is to find out the minimal required coverage to obtain chromosome-level assemblies of individuals to keep the cist low and yet can be used as a reference. A publicly available dataset with 1800x coverage of PacBio HiFi data (SRR11434954) generated for a E. coli strain from SRA database offered an opportunity to systematically test the coverage requirement. Randomly extracted reads representing 1x, 2x, 3x, 4x, 8x, 12x, 16x and 20x coverage were assembled using hifiasm. The experiment was repeated 5 times. The N50, L50, and the assembled genome-size of all five experiments across increasing coverage is shown Figure 1. According to this, near 12x coverage the entire genome is represented in contigs as shown by the genome size. Also, at 16x coverage of PacBioHiFi reads the longest contig represents the chromosome.

**Figure 1:**
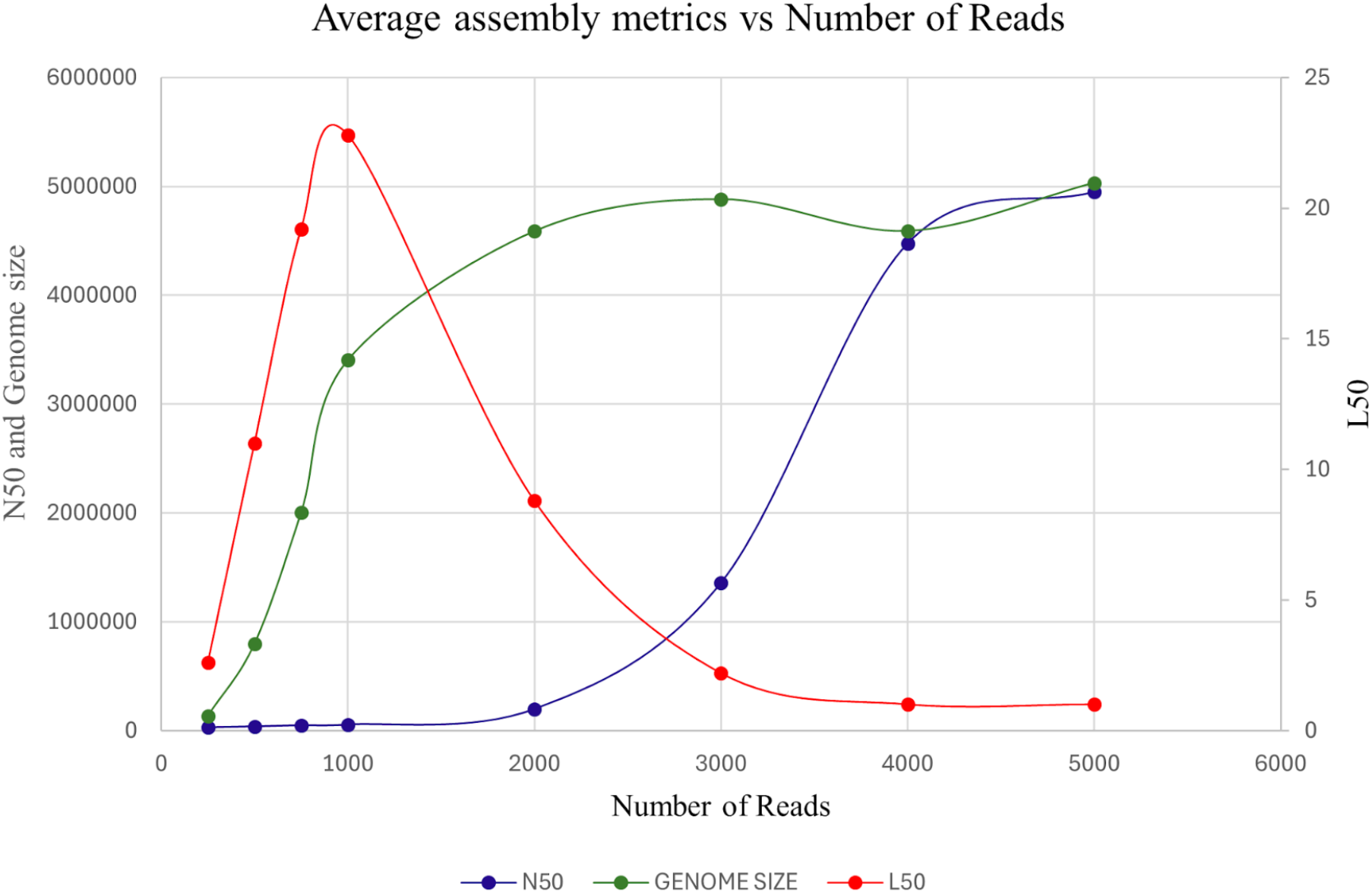
Graph depicting change in L50, N50 and genome size with increasing coverage

### Assembly of KIn1

DNA was extracted from blood of an individual from Karnataka (KIn1) after ethics clearance (NHM/SPM/4-PART FILE/2020-21) and after ethnicity screening consistent with individuals screened for the 1000 Genomes Project. PacBio sequencing produced 12x coverage of HiFi reads, which were assembled using hifiasm after filtering for bell adapters, resulting in contigs L50 of 3,417. The contigs were scaffolded using publicly available HiC data from an individual (HG04217, IGSR), most closely related by ethnicity to the subject using YaHS (Yet Another HiC Scaffolder). The resulting scaffold N50 was 137 Mb and L50 was 9, suggesting a chromosome-level assembly. The longest scaffolds representing chromosomes, except for acrocentric chromosomes, covered over 90% (Table 1, Column 5) relative to T2T assembly.

**Table 1:**
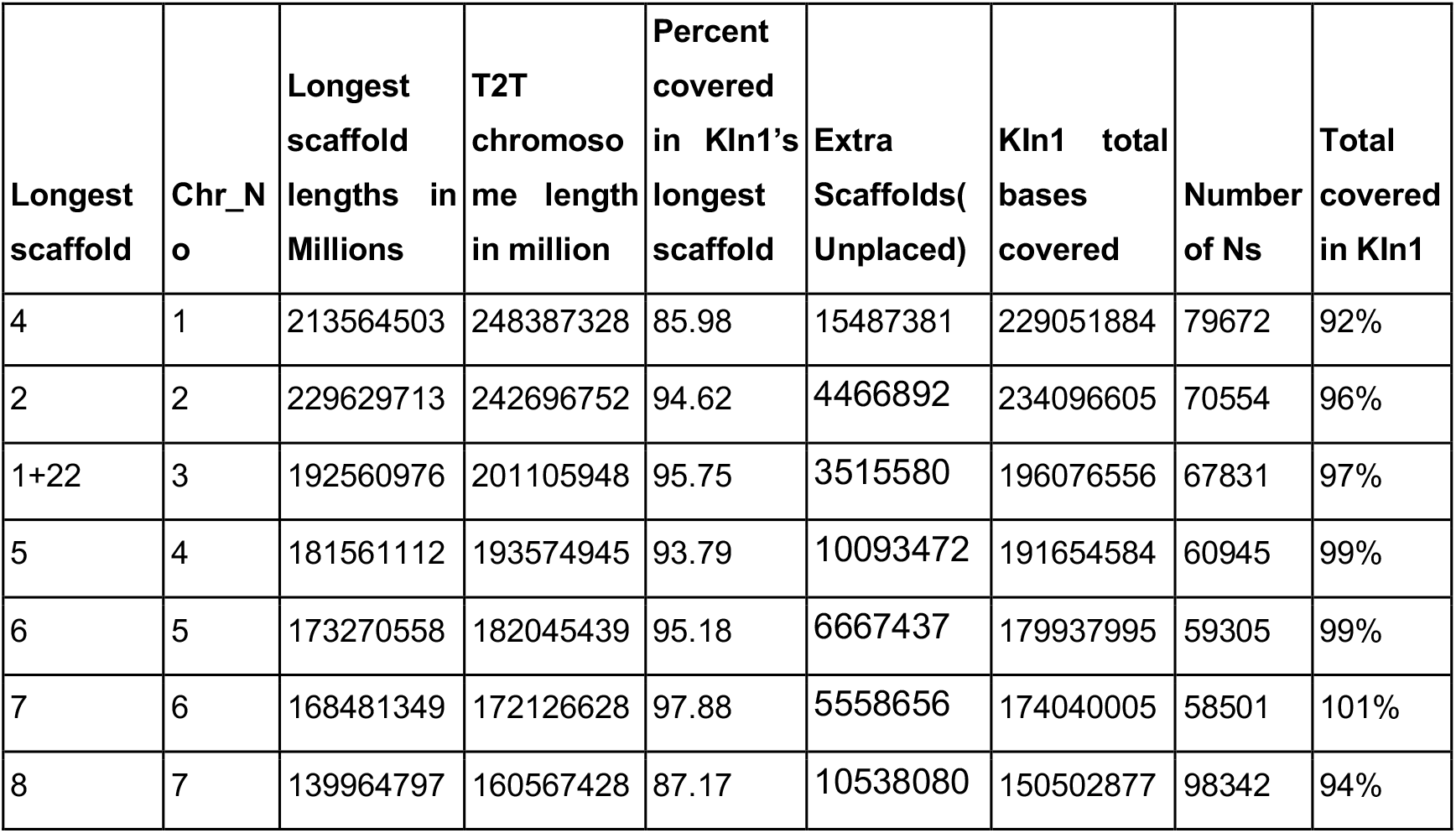
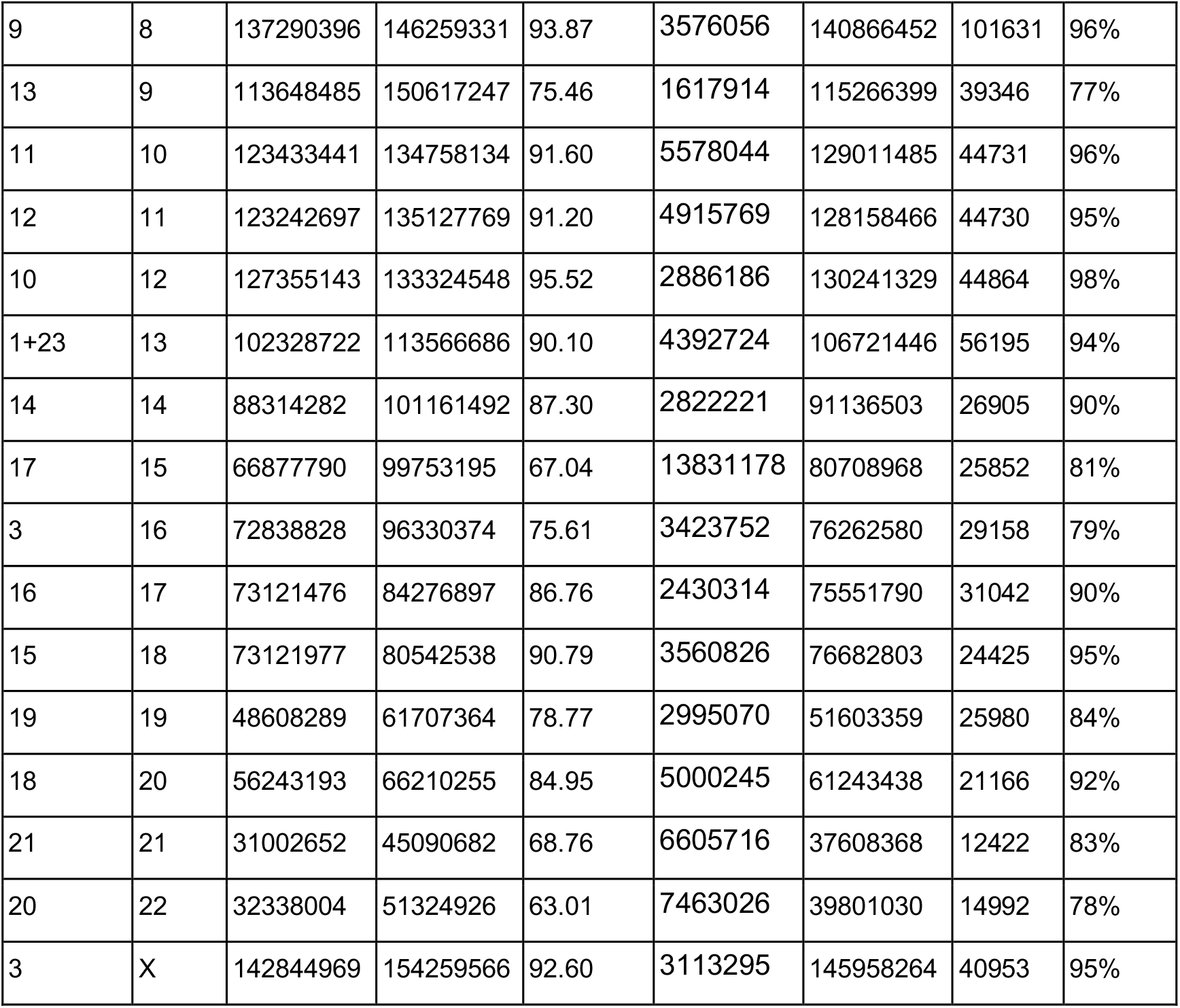
Statistics of Kin1 assembly.

| **Longest scaffold** | **Chr_N**<br>**o** | **Longest scaffold lengths in Millions** | **T2T chromoso**<br>**me length in million** | **Percent covered in KIn1's longest scaffold** | **Extra Scaffolds( Unplaced)** | **KIn1 total bases covered** | **Number of Ns** | **Total covered in KIn1** |
| --- | --- | --- | --- | --- | --- | --- | --- | --- |
| 4 | 1 | 213564503 | 248387328 | 85.98 | 15487381 | 229051884 | 79672 | 92% |
| 2 | 2 | 229629713 | 242696752 | 94.62 | 4466892 | 234096605 | 70554 | 96% |
| 1+22 | 3 | 192560976 | 201105948 | 95.75 | 3515580 | 196076556 | 67831 | 97% |
| 5 | 4 | 181561112 | 193574945 | 93.79 | 10093472 | 191654584 | 60945 | 99% |
| 6 | 5 | 173270558 | 182045439 | 95.18 | 6667437 | 179937995 | 59305 | 99% |
| 7 | 6 | 168481349 | 172126628 | 97.88 | 5558656 | 174040005 | 58501 | 101% |
| 8 | 7 | 139964797 | 160567428 | 87.17 | 10538080 | 150502877 | 98342 | 94% |
| 9 | 8 | 137290396 | 146259331 | 93.87 | 3576056 | 140866452 | 101631 | 96% |
| 13 | 9 | 113648485 | 150617247 | 75.46 | 1617914 | 115266399 | 39346 | 77% |
| 11 | 10 | 123433441 | 134758134 | 91.60 | 5578044 | 129011485 | 44731 | 96% |
| 12 | 11 | 123242697 | 135127769 | 91.20 | 4915769 | 128158466 | 44730 | 95% |
| 10 | 12 | 127355143 | 133324548 | 95.52 | 2886186 | 130241329 | 44864 | 98% |
| 1+23 | 13 | 102328722 | 113566686 | 90.10 | 4392724 | 106721446 | 56195 | 94% |
| 14 | 14 | 88314282 | 101161492 | 87.30 | 2822221 | 91136503 | 26905 | 90% |
| 17 | 15 | 66877790 | 99753195 | 67.04 | 13831178 | 80708968 | 25852 | 81% |
| 3 | 16 | 72838828 | 96330374 | 75.61 | 3423752 | 76262580 | 29158 | 79% |
| 16 | 17 | 73121476 | 84276897 | 86.76 | 2430314 | 75551790 | 31042 | 90% |
| 15 | 18 | 73121977 | 80542538 | 90.79 | 3560826 | 76682803 | 24425 | 95% |
| 19 | 19 | 48608289 | 61707364 | 78.77 | 2995070 | 51603359 | 25980 | 84% |
| 18 | 20 | 56243193 | 66210255 | 84.95 | 5000245 | 61243438 | 21166 | 92% |
| 21 | 21 | 31002652 | 45090682 | 68.76 | 6605716 | 37608368 | 12422 | 83% |
| 20 | 22 | 32338004 | 51324926 | 63.01 | 7463026 | 39801030 | 14992 | 78% |
| 3 | X | 142844969 | 154259566 | 92.60 | 3113295 | 145958264 | 40953 | 95% |

The linearity of the longest scaffolds representing each chromosome of KIn1 was validated using virtual markers of 2000 base pairs sampled every 1 million bases from the T2T assembly. These markers were mapped against the scaffolds to identify those associated with chromosomes and authenticate the linearity of the assembly. Figure 2 shows dot plots for representative chromosomes and all chromosomes with enlarged plots shown in Supplementary Figure 1A-X. Table 1 lists the scaffolds associated with chromosomes (Column 1), their lengths (Column 3), and the percent coverage (Column 5) of these scaffolds compared to T2T-CHM13-v2.0.

**Figure 2:**
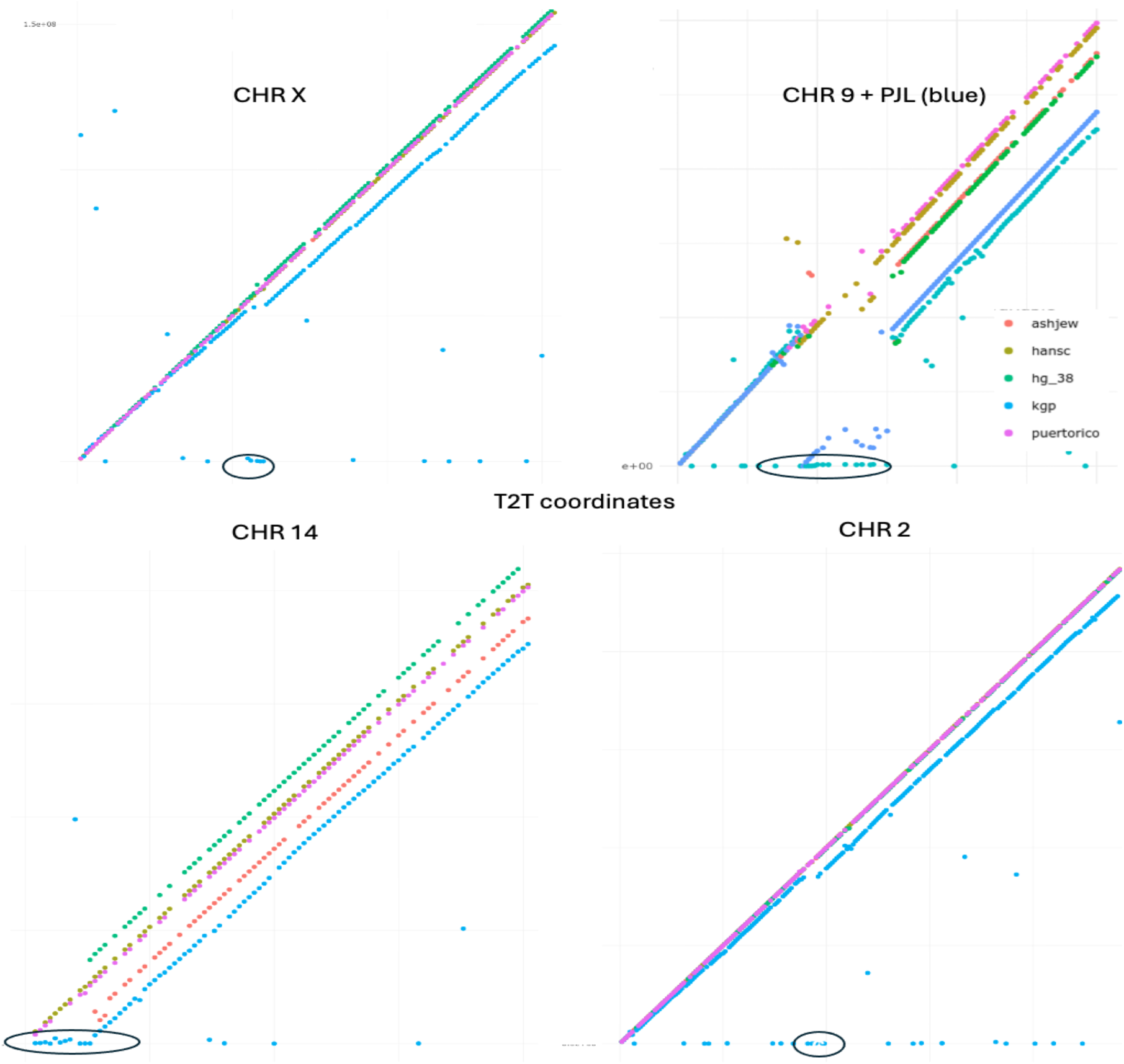
A comparative dot plot of a few select chromosomes from Ash1, PR1, Han1, hg38 and KIn1 compared to T2T. Orange is Ash1, yellow is Hans1, green is hg38, purple is PR1, cyan is KIn1 and blue is PJL. Unplaced centromere scaffold from KIn1 is circled on x axis.

A majority of the chromosomes were represented intact in single long scaffolds with the exception of 3, 13, X and 16. The q arms of chromosomes 3 and 13 was found assembled together in scaffold 1 head-to-head near centromeres, which were split using mapping to chromosome 3 and 13 of T2T. The p-arm of chromosome 3 was found intact in scaffold 22 and was stitched to provide full length chr3 for KIn1 genome. Similarly, chromosome 16 and X assembled together in scaffold 3 and were split using high resolution markers from T2T near the potential joining region. The percent lengths of the longest scaffolds corresponding to each chromosome is given in column 5 relative to T2T assembly (Column 4). Column 9 provides the percentage of bases represented in each chromosome from all scaffolds/contigs including the longest with respect to T2T. Accordingly, among the meta-centric chromosomes, except for chromosomes 9, 16, and 19, more than 90% of the T2T assembly is represented in KIn1. Out of the exceptions chromosomes 9 and 16 are known to have large centromere, notorious for gaps in major assemblies. The unplaced scaffolds/contigs mapping to T2T are included in the final genome file with their respective T2T coordinates (PRJNA1138089).

We have compared individual genomes from different ethnicities such as Puerto Rican (PR1), Ashkenazi (Ash1), Hans Chinese (Hans1), and Northern European (T2T) and KIn1 by mapping 2 Kb markers spaced every one million bases along the chromosomes of T2T-CHM13-v2.0. To include the genome of another individual from a South Asian descent in our comparison, we have used the HiC data and the publicly available contigs from the pangenome project to assemble the genome of an individual (HG03492) from Punjab. Figure 1 displays a dot plot of T2T markers aligned to the respective genomes: Ash1 in orange, Han1 in yellow, hg38 in green, PR1 in pink, KIn1 in cyan and Pun1 in blue.

Unlike Ash1, Han1 and PR1, KIn1 assembly has not undergone gap filling or manual correction for mis-assemblies within the euchromatic regions. However, the blue dots along the diagonal from the longest scaffolds of the KIn1 assembly, aligning with T2T markers, cover approximately 90% of the chromosomes, suggesting the achieved linearity of scaffolding using HiC data from HG04217, an individual most ethnically related to KIn1. The cyan dots scattered across the X-axis are unplaced scaffolds hitting with the T2T markers, which cover ∼ 5% of the genome on the average. As shown by black circles in Figure 2, there are clusters of unplaced scaffolds near the centromeres where HiC data may be lacking paired links. The other unplaced scaffolds fill the gaps in the longest scaffolds left out during HiC scaffolding. Particularly, in chromosome 15 there are 3 scaffolds millions in length are unplaced (Supplementary Figure 1).

The linearity of the genome assemblies compared in this report is evident from dots lining up along the diagonal. Since gaps in Ash1 and PR1 are filled with sequences from T2T and assemblies and corrected manually, it is not surprising that they are arranged along the diagonal. The shift in KIn1 scaffolds away from the diagonal near the centromere is consistent with the unplaced scaffolds from centromere. This effect is most dramatic in chromosome 9 (Figure 1) and 16 (Supplementary Figure 1), which are known to have the longest centromeres. In fact, even in PJL with contigs generated from much higher coverage also missed placing centromeres in these chromosomes. The number of unplaced scaffolds outside of the centromere may reflect variation in the regions of the genome/chromosome between the HG04217 with respect to KIn1. In all genomes compared here, considerable non-linearity within the centromeric regions is found, suggesting lack of conservation within centromeres across ethnicity. The gaps along the diagonal in all genomes compare here may represent population specific variation.

### Interrogation of inversions and translocations greater than 1 Mb

As shown in Figure 3 and Supplementary Figure 1, we find two major inversions compared to T2T that are not reported for Ash1, PR1 and Han1 assemblies. For example, we find a 9 Mb inversion in the euchromatic regions of chr7. Figure 3 left shows chromosome 7 with inversions within the p-arm. Upon interrogation of the breakpoints for the distal inversion, it is found that there are no contigs spanning the breakpoints suggesting that the inversion is a results of scaffolding using HiC reads from HG04217. In the right we provide the exact coordinates of the 100 Ns introduced near the two breakpoints by the tool YaHS during the scaffolding representing gaps. These Ns represent 10,000 bps and 15000 bps in the distal and proximal gaps respectively in chr 7 of KIn1 assembly as shown in Figure 3 right. Our dot plot in Supplementary Figure 1 also shows the only unique inversion in Han1 reported in chr 8.

**Figure 3:**
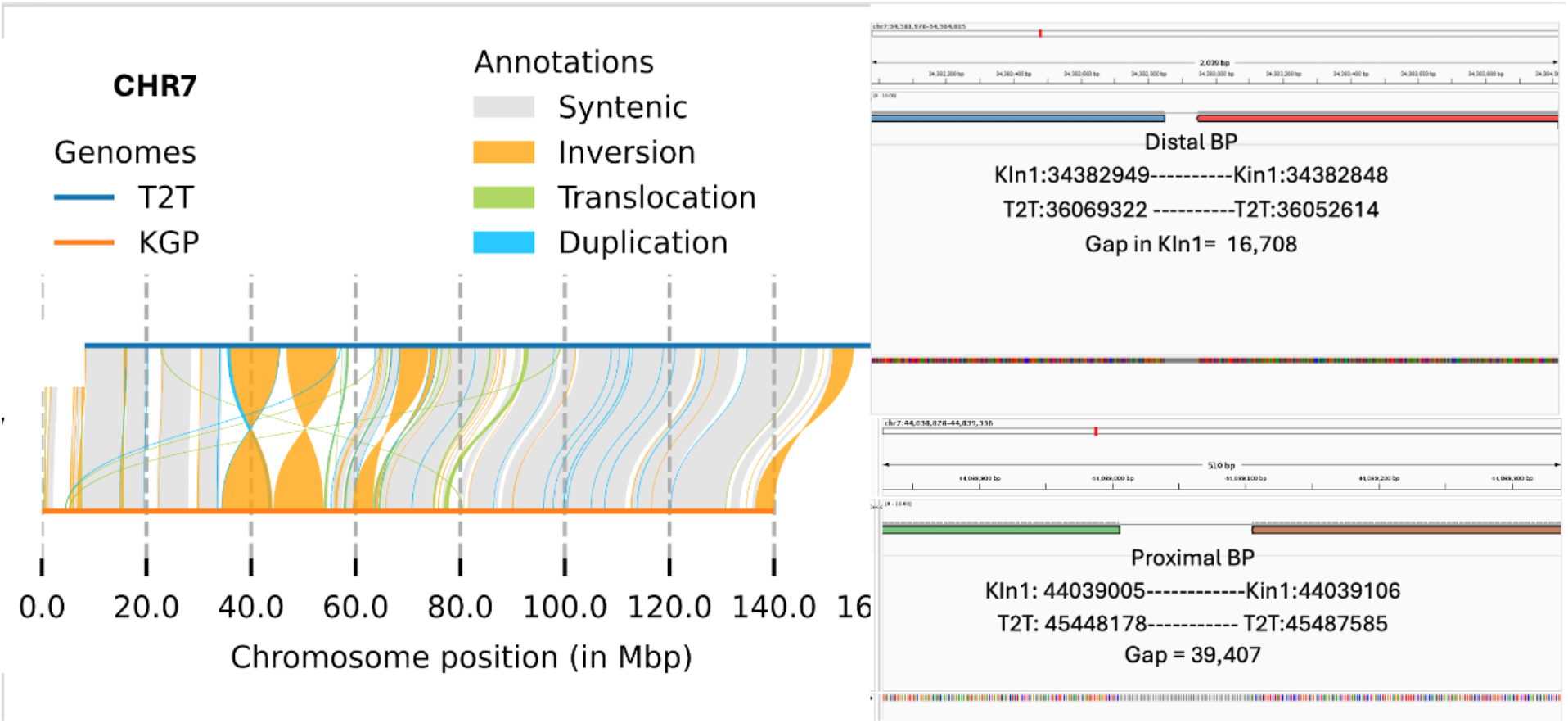
Showing large inversions(orange) in chromosome 7 of KIn1 against T2T-CNM13-v2.0. On the right we show exact coordinates of breakpoints.

In chromosomes 17 and 20 we find large translocations. In chromosome 17 of KIn1 a telomere-to-telomere fusion event and in chromosome 20 the translocation brings the tail of the q-arm to the beginning of q-arm next to the centromere. For lack of karyotype on this individual, we restrained from manual fixing. Furthermore, such large movements, does not affect annotation and/or usefulness of the assembly. We also noticed such a translocation in chromosome 18 of Pun1 assembly (see Supplementary Figure 1).

### Variant calls using KIn1 as reference

We have performed variant calls of individuals including three ethnic groups represented in IGSR and a few distal ones such as Great Britain. Table 3 shows the number of variants predicted from the samples using WGS data from ITU, STU, and British from UK. As expected, the number of variants between ITU, individuals from the neighboring state to Karnataka, the closest ethnicity to the individual representing KIn1, is much lower than and those from STU, a Tamil speaking ethnicity from Sri Lanka. Accordingly, ITUs representing the ethnicity from a state neighboring Karnataka, have the least variants ranging from 1.8-1.9 million, STU in the range of 2.3-2.6 million, and Great Britain ranging from 2.5-2.9 million. The number of variants called from the same two individuals from Great Britain using T2T as reference range from 2.7-3.3 million, which is more by 13% with respect to KIn1 assembly as reference, perhaps reflecting gaps in KIn1 compared to T2T.

**Table 3:**
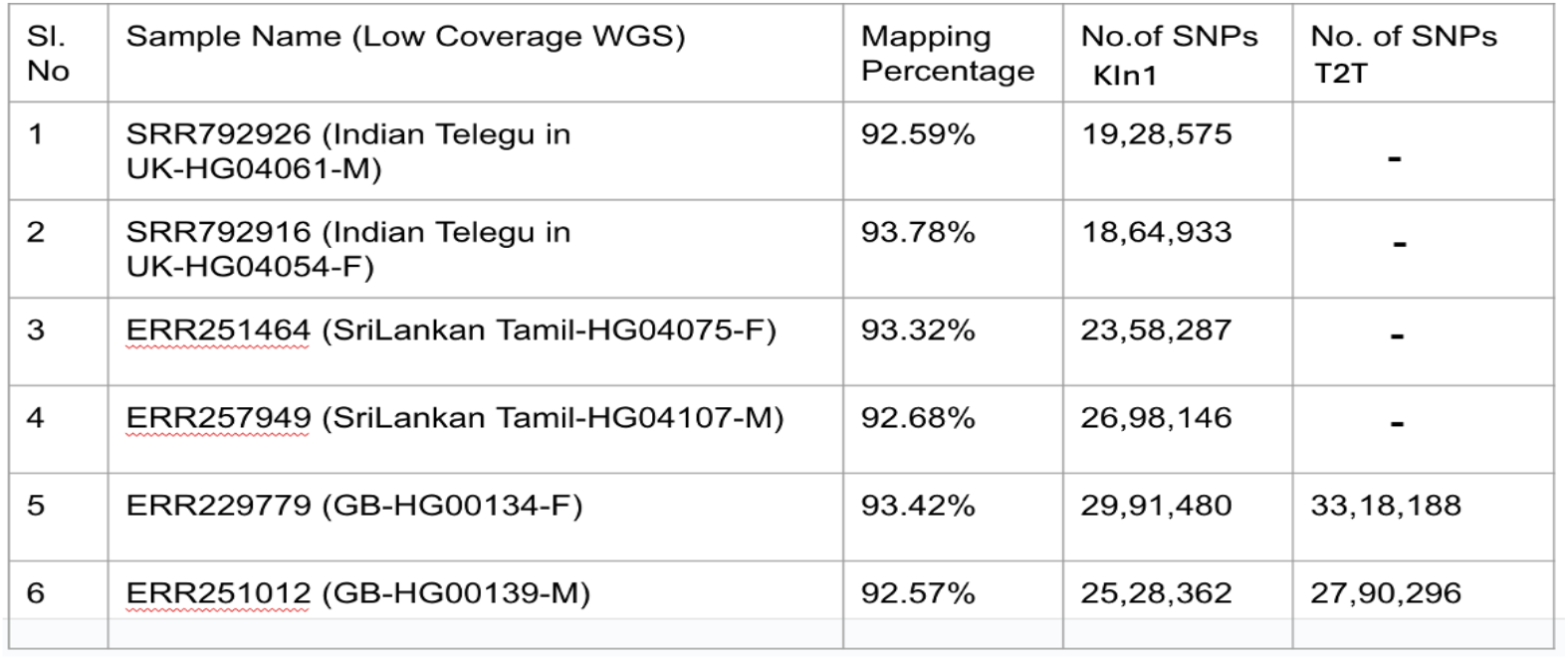
Number of variants called using KIn1 as reference compared to that using T2T.

| Sl. No | Sample Name (Low Coverage WGS) | Mapping Percentage | No.of SNPs KIn1 | No. of SNPs T2T |
| --- | --- | --- | --- | --- |
| 1 | SRR792926 (Indian Telugu in UK-HG04061-M) | 92.59% | 19,28,575 | - |
| 2 | SRR792916 (Indian Telugu in UK-HG04054-F) | 93.78% | 18,64,933 | - |
| 3 | ERR251464 (SriLankan Tamil-HG04075-F) | 93.32% | 23,58,287 | - |
| 4 | ERR257949 (SriLankan Tamil-HG04107-M) | 92.68% | 26,98,146 | - |
| 5 | ERR229779 (GB-HG00134-F) | 93.42% | 29,91,480 | 33,18,188 |
| 6 | ERR251012 (GB-HG00139-M) | 92.57% | 25,28,362 | 27,90,296 |

### Annotation of KIn1 assembly

We used liftoff from T2T annotation to annotate KIn1 assembly. Out of the 61,312 genes annotated in T2T, 59,590 could be lifted off intact authenticating the linearity of the assembly within the gene loci as shown in Figure 4. We also predicted genes from KIn1 using Augustus. The number of protein coding genes are 62,000. As less as 31 genes has gained stop.

**Figure 4:**
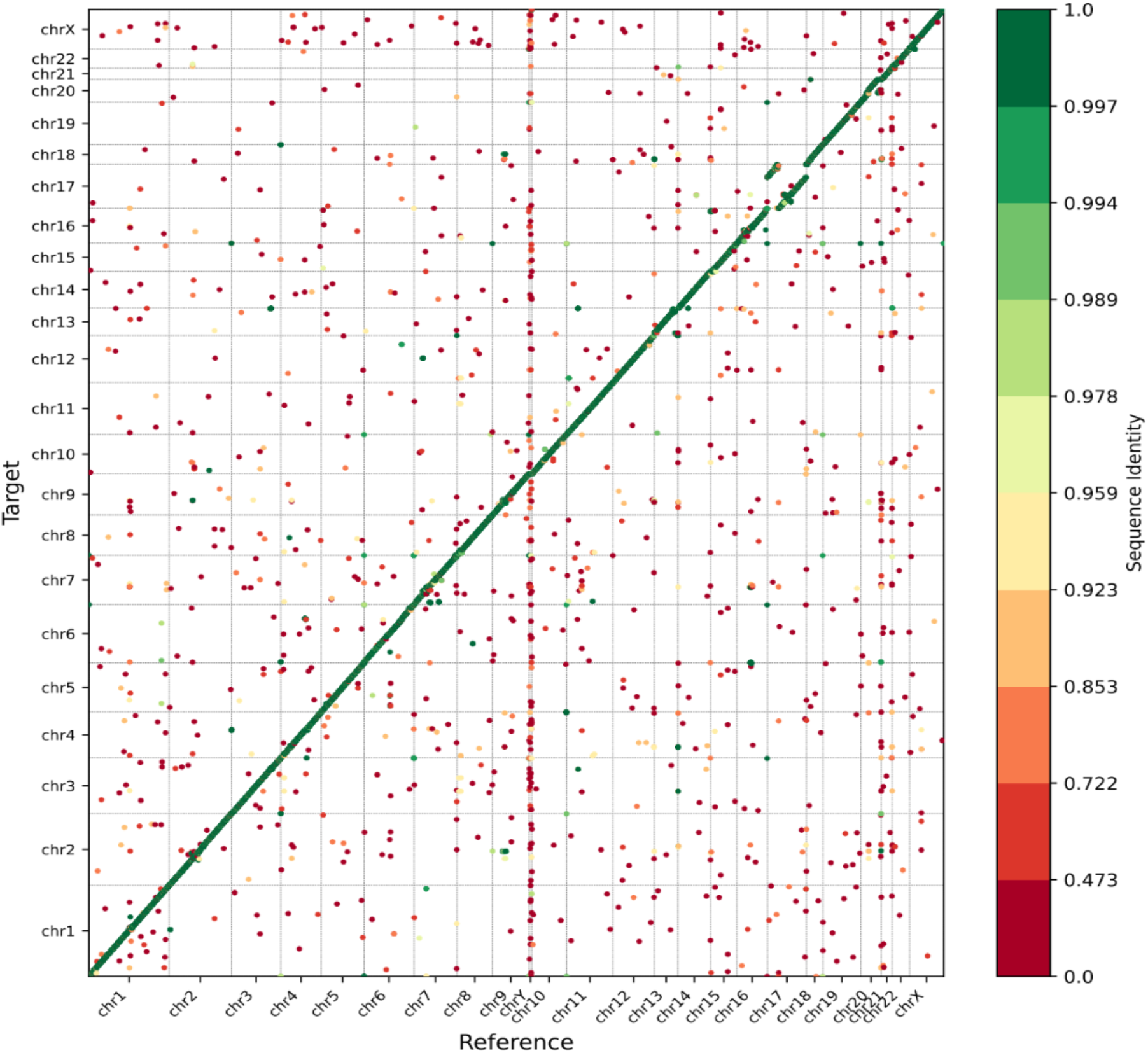
Shows synteny of genes from all chromosomes between T2T and KIn1

We have compared variants of KIn1 genes against T2T using LiftOffTools and against hg38 using Augustus predicted KIn1 genes. Table 4 below provides the statistics for Kin1 compared to variants of Han1 against T2T.

**Table 4:** Variation with T2T within protein coding gene between KIn1 and Han1.

| Numbers | Han1 | KIn1 |
| --- | --- | --- |
| Number of total annotated gene | 181708 | 181708 |
| Identical | 14130 | 16951 |
| Start_lost | 21 | 743 |
| Stop_Gained | 31 | 31 |
| Frameshift | 143 | 1040 |
| Synonymous | 2422 | 1844 |
| Nonsynonymous | 3200 | 2551 |
| 5'_truncated |  | 2 |
| 3'_truncated |  | 228 |
| inframe_deletion | 92 | 582 |
| Inframe_insertion | 110 | 97 |

Except for Start_Lost and frameshift mutations the number of instances of other variations in KIn1 relative to T2T-CHM3-2.0 are very close to that reported for Han1 gapless assembly^5^. An interrogation of the 743 Start-Lost against T2T genome only 183 are primary transcripts. In any event, of the 698 genes out of 743 that are present in KIn1 proteome, 671 genes used alternate Start codon and retains intact structural domains based on Conserved Domain Database (CDD), suggesting functional proteins despite use of alternate start site.

## DISCUSSION

The goal of a de novo assembly of individual genomes is to find large structural changes among individuals such as inversions and translocations. However, the academic temptation to produce high-resolution chromosome-level assembly, without gaps, defeats this goal. For example, during the assembly of PR1^4^ the orientations of some contigs/scaffolds were adjusted to conform to T2T-CHM3-2.0 assembly. Here, we have tried to use 12x coverage of reads from PacBio HiFi derived from blood sample of a non-GIAB and/or non-IGSR individual from Karnataka, India (KIn1) to assemble contigs and used a hybrid approach for scaffolding using HiC data derived from HG04217, an individual with ethnicity closest to KIn1 in IGSR. With a few exceptions, the longest scaffolds covered nearly 90% of the chromosomes providing an L50 of 9 and N50 of 137 Mb, a statistics very close to the lowest and highest attainable values of 8 and 147 Mb for human genomes respectively. Together with unplaced scaffolds and unabsorbed contigs, the assembly totals 2.84 Gbases. The final assembly contains only ∼1 million Ns covering 0.03% of the genome. The majority of the unabsorbed scaffolds and contigs are from centromeric regions, perhaps from lack of reads pairs in HiC connecting the centromere to euchromatic regions.

We used a novel approach for comparative genomics by mapping virtual markers of 2000 bases representing every one million bases on the T2T chromosomes on to other genomes compared here. Our approach is validated by confirming an inversion reported in the p-arm of chromosome 8 in Han1 assembly (Supplementary Figure 1). The linearity of markers on to the longest KIn1 scaffolds suggest near chromosome-level assembly. Interestingly, gaps in the mapping, without shifting from the diagonal, also represent unique variation among genomes compared here.

The KIn1 assembly has inversions not present in individual genomes compared here. For example, in chromosome 7 and chromosome 17 we see large inversions. It is likely that these inversions are resulting from HiC reads from HG04217 during scaffolding as shown in Figure 3. We called variants from WGS reads from samples from IGSR including diverse ethnicity using Kinb1 as reference. These include ITU individuals (HG04061 and HG04054), STU individuals (HG04075 and HG04107), and British from UK (HG00134 and HG00139). Although, individuals in Karnataka and Andhra Pradesh are separated by linguistics, the lower number of variants between KIn1 and ITU (HG04061/HG04054) compared with individuals from STU and UK suggests their close kinship vindicating our use of HiC data for scaffolding KIn1. This also may be suggestive of ITU individuals in IGSR collection may be from a region of Andhra Pradesh bordering Karnataka. Also, assembly of KIn1, for the first time, shows the genetic diversity between KIn1 and STUs, also neighboring states in India.

We believe that this is the first chromosome-level assembly of an individual of Indian origin acting as a reference genome for individuals from this ethnicity and for those from neighboring states. Also, KIn1 assembly will act as a reference facilitating high-resolution population genomics study in India.

## MATERIALS AND METHOD

### Samples

Blood samples from several individuals were collected with consent (NHM/SPM/4-PART FILE/2020-21) after extensive family pedigree information relating them to the ethnicity and the samples were labeled anonymous. DNA from one of these samples were isolated and sent for sequencing.

### Analysis strategy

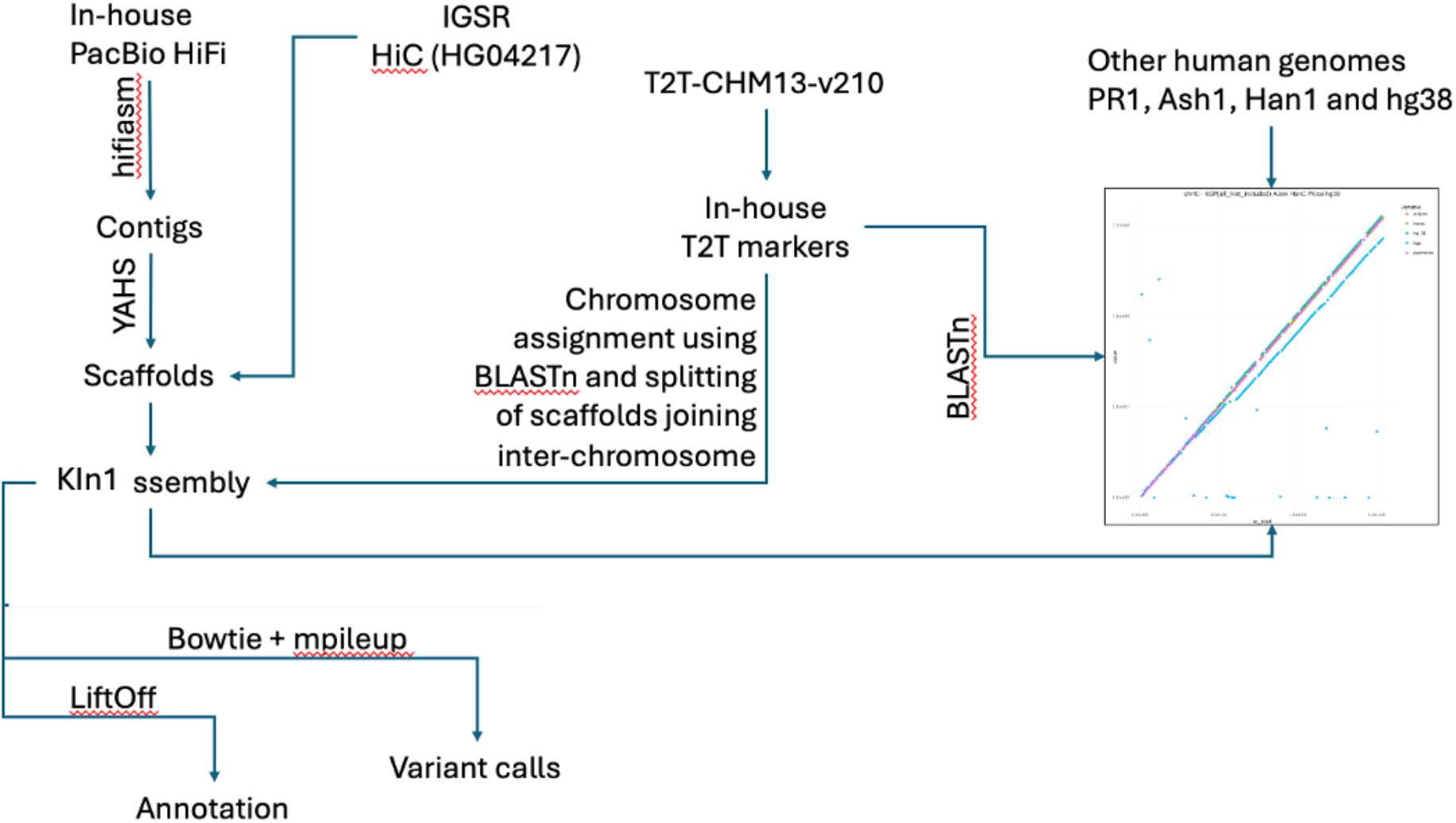

### Assembly

Pacbio tools was used to convert CCS subreads to Hifi reads by filtering for reads with Q>20 and subreads>=3. HiFiAdapterFilt was used to remove pacbio bell adapter contamination.

Resulting reads were assembled into contigs using hifiasm^11^.

Contigs were used to generate scaffolds using YAHS^12^ with 30x HiC data downloaded from IGSR HG04217. Each contigs in the scaffold is separated by 100 Ns. Subreads were used to polish the assembly by mapping on to the scaffolds.

### Validation of Assembly

Markers of length 2kb were generated with a scale of one million bases on T2T assembly using an in-house tool. These markers were mapped using BLASTn on assembled human genomes such as hg38, PR1, Ash1, Han1, and KIn1 to both validate KIn1 assembly based on linearity of the markers and to identify major inversions and translocations in KIn1 compared to others. From the blast output we filtered the hits which were giving the percentage identity close to 100% i.e (∼99.2% to ∼99.87) also we kept the hits which were hitting majority of the scaffolds. The start and end coordinates of the markers against the scaffolds from the blast hits were then used to create dot plots shown in Figure 1 and Supplementary Figure 1.

### Variant Prediction

Illumina short reads from select IGSR individuals were downloaded and mapped to respective assembled genomes using bowtie and mpileup was used to call variants.

## DATA AVAILABILITY

The KIn1 assembly can be found under the accession PRJNA1138089 from NCBI. HiC data used can be downloaded from HG04217 under IGSR. Raw PacBio HiFi reads will be made available under the same project ID shortly.

## ACKNOWLEDGEMENT

PacBio HiFi reads were generated using services from Nucleome Informatics. The authors thank GoK for funding for sequencing cost and data analysis personnel via BioIT grant and computing infrastructure via Department of IT, BT and ST.

## REFERENCES

1. Lander, E. S. et al. Initial sequencing and analysis of the human genome. Nature 409, 860–921 (2001).

2. Schneider, V. A. et al. Evaluation of GRCh38 and de novo haploid genome assemblies demonstrates the enduring quality of the reference assembly. Genome Res. 27, 849–864 (2017).

3. Nurk, S. et al. The complete sequence of a human genome. Science 376, 44–53 (2022).

4. Zimin, A. V. et al. A reference-quality, fully annotated genome from a Puerto Rican individual. Genetics 220, iyab227 (2022).

5. Chao, K.-H., Zimin, A. V., Pertea, M. & Salzberg, S. L. The first gapless, reference-quality, fully annotated genome from a Southern Han Chinese individual. G3 Bethesda Md 13, jkac321 (2023).

6. Shumate, A. et al. Assembly and annotation of an Ashkenazi human reference genome. Genome Biol. 21, 129 (2020).

7. Church, G. M. The personal genome project. Mol. Syst. Biol. 1, 2005.0030 (2005).

8. Zook, J. M. et al. Extensive sequencing of seven human genomes to characterize benchmark reference materials. Sci. Data 3, 160025 (2016).

9. A draft human pangenome reference - PubMed. https://pubmed.ncbi.nlm.nih.gov/37165242/.

10. Garg, S. et al. Chromosome-scale, haplotype-resolved assembly of human genomes. Nat. Biotechnol. 39, 309–312 (2021).

11. Cheng, H., Concepcion, G. T., Feng, X., Zhang, H. & Li, H. Haplotype-resolved de novo assembly using phased assembly graphs with hifiasm. Nat. Methods 18, 170–175 (2021).

12. Zhou, C., McCarthy, S. A. & Durbin, R. YaHS: yet another Hi-C scaffolding tool. Bioinforma. Oxf. Engl. 39, btac808 (2023).

