## Supplementary Material for "The first assembled chromosome-level genome of an individual from Karnataka, India"

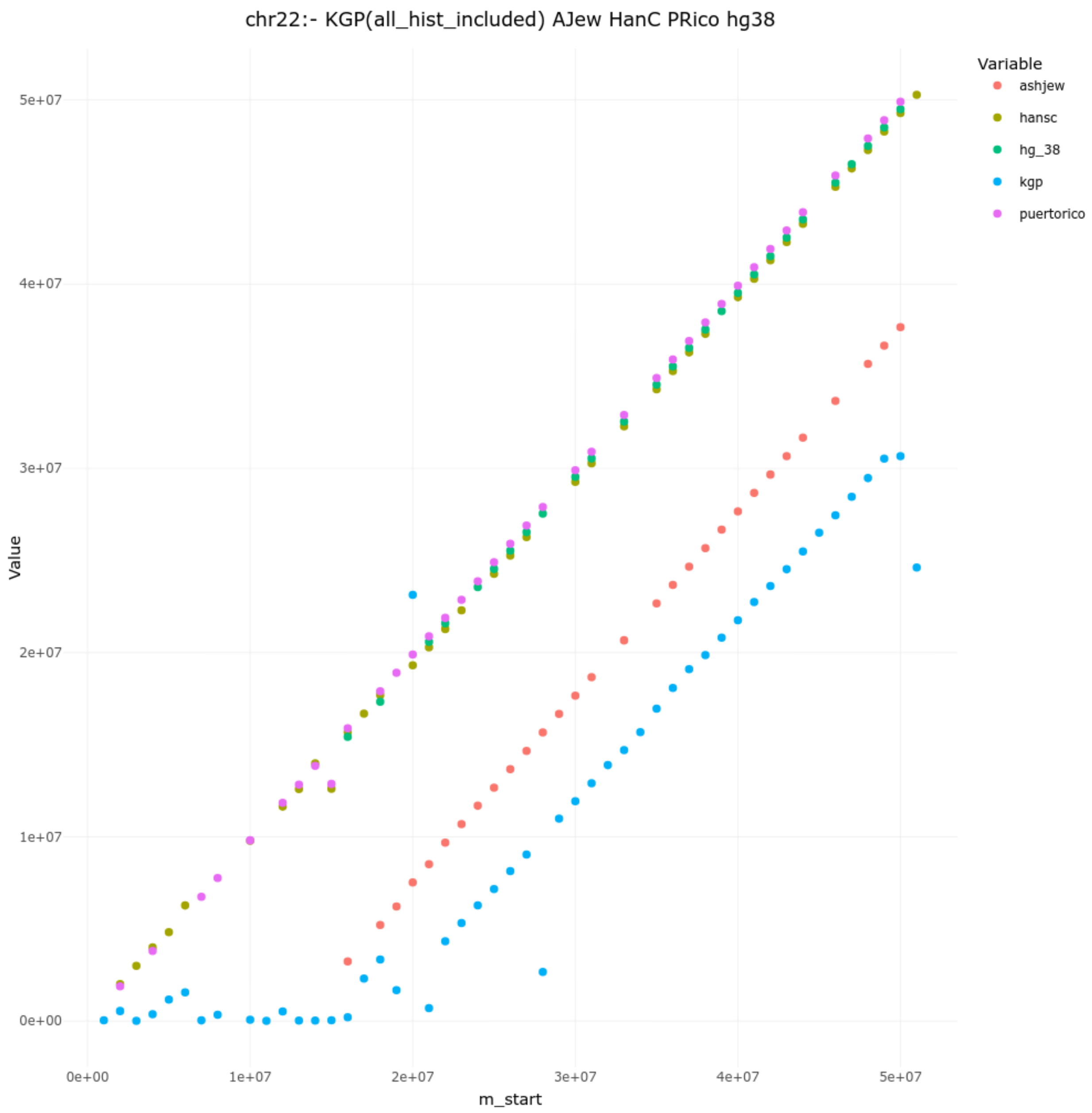
chr22:- KGP(all_hist_included) AJew HanC PRico hg38
Variable
ashjew
hansc
hg_38
kgp
puertorico
Value
5e+07
4e+07
3e+07
2e+07
1e+07
0e+00
0e+00
1e+07
2e+07
3e+07
4e+07
5e+07
m_start

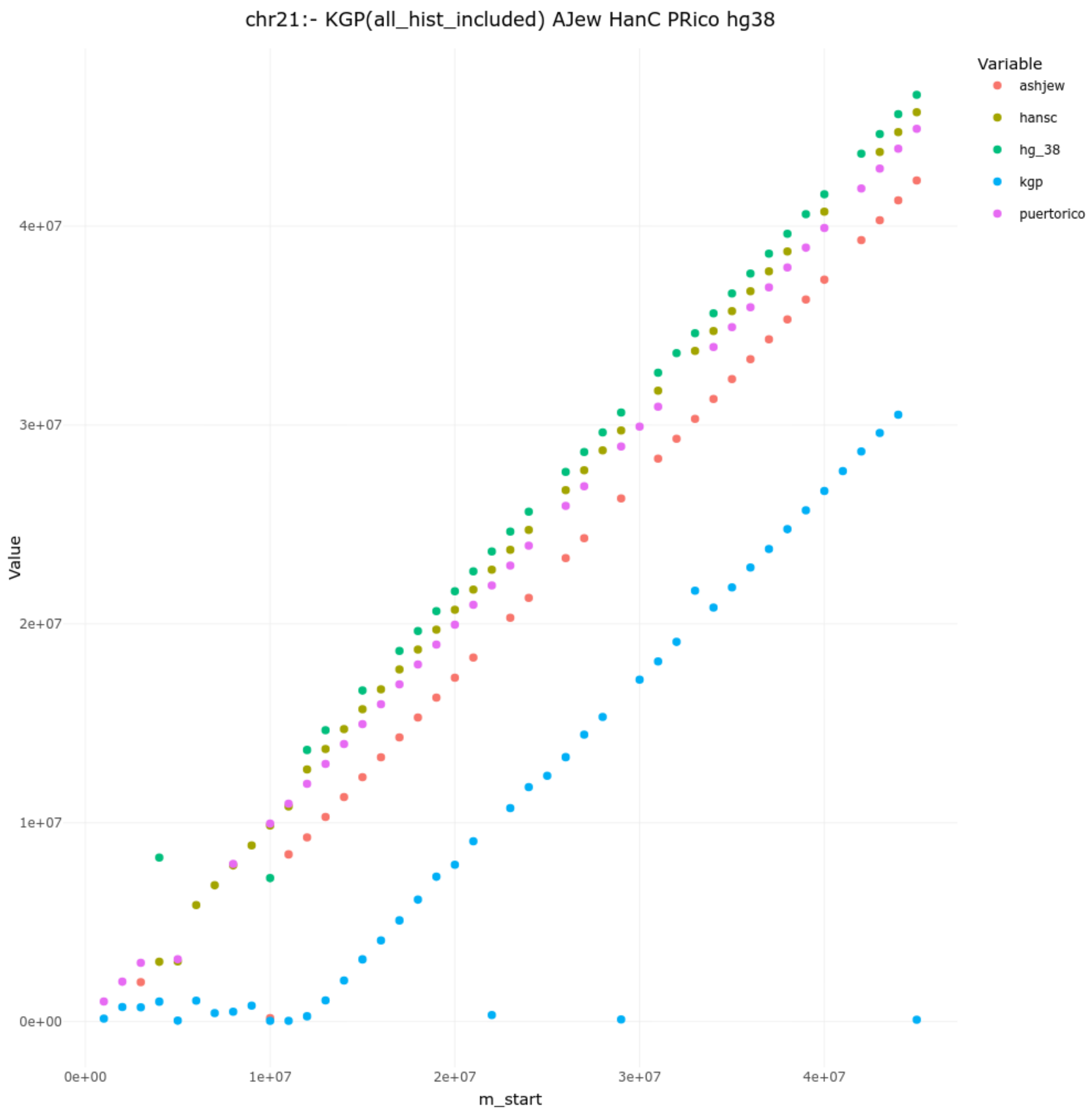

chr21:- KGP(all_hist_included) AJew HanC PRico hg38
Variable
ashjew
hansc
hg_38
kgp
puertorico
Value
4e+07
3e+07
2e+07
1e+07
0e+00
0e+00
1e+07
2e+07
3e+07
4e+07
m_start

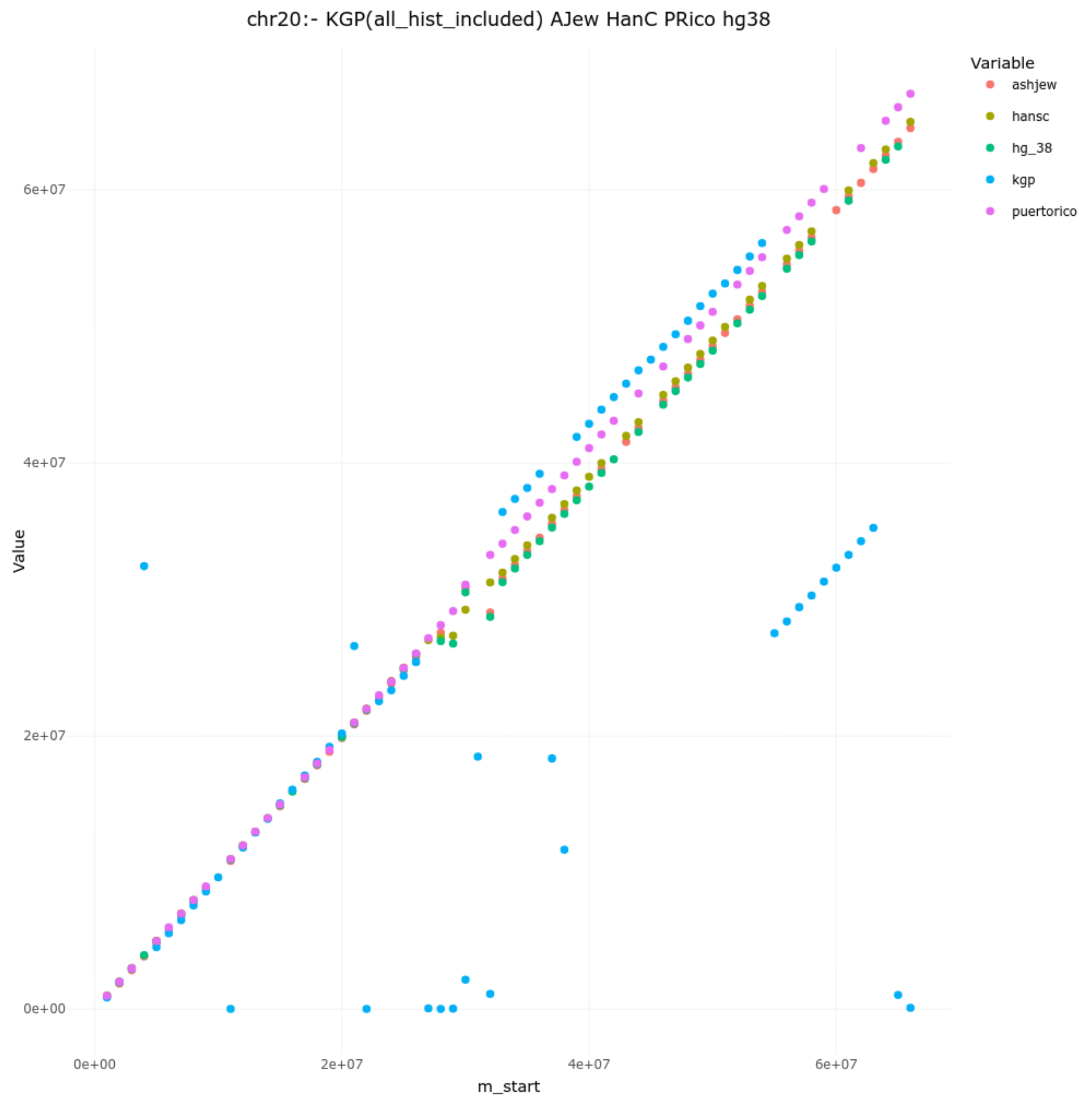
chr20:- KGP(all_hist_included) AJew HanC PRico hg38
Variable
ashjew
hansc
hg_38
kgp
puertorico
Value
6e+07
4e+07
2e+07
0e+00
0e+00
2e+07
4e+07
6e+07
m_start

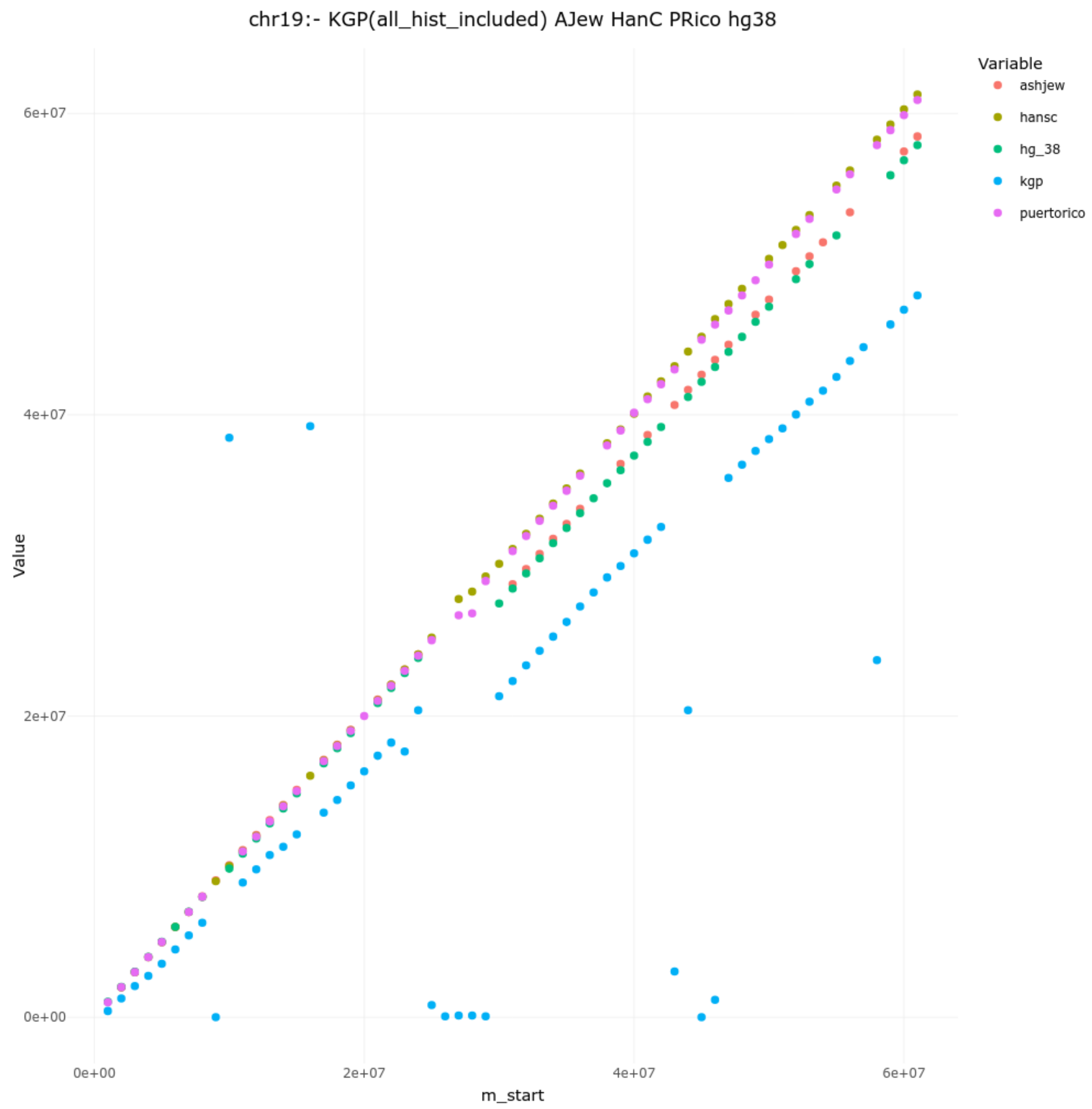

chr19:- KGP(all_hist_included) AJew HanC PRico hg38
Variable
ashjew
hansc
hg_38
kgp
puertorico
Value
6e+07
4e+07
2e+07
0e+00
m_start
0e+00
2e+07
4e+07
6e+07

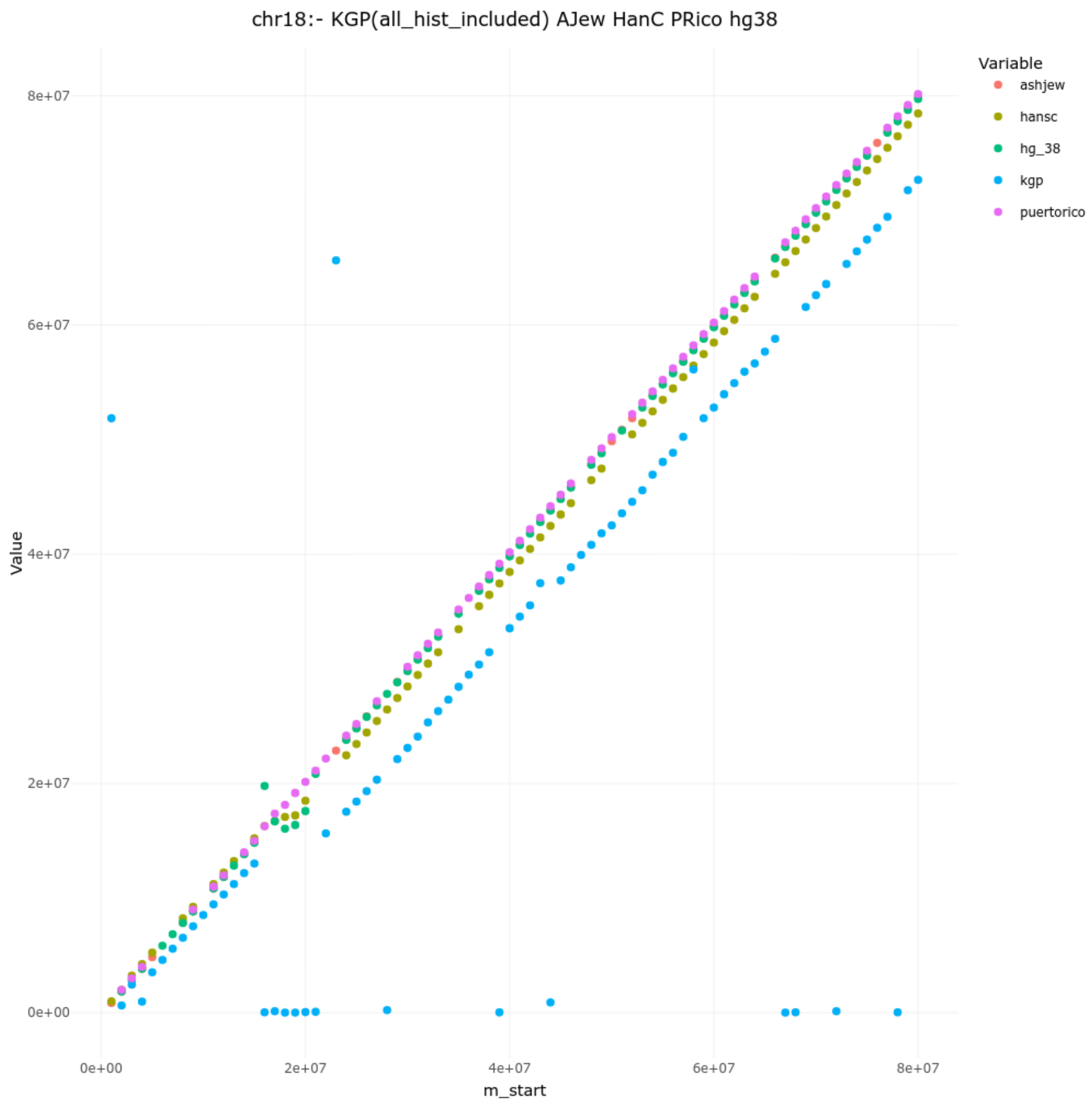

chr18:- KGP(all_hist_included) AJew HanC PRico hg38
Variable
ashjew
hansc
hg_38
kgp
puertorico
8e+07
6e+07
4e+07
2e+07
0e+00
Value
0e+00
2e+07
4e+07
6e+07
8e+07
m_start


CHR 17

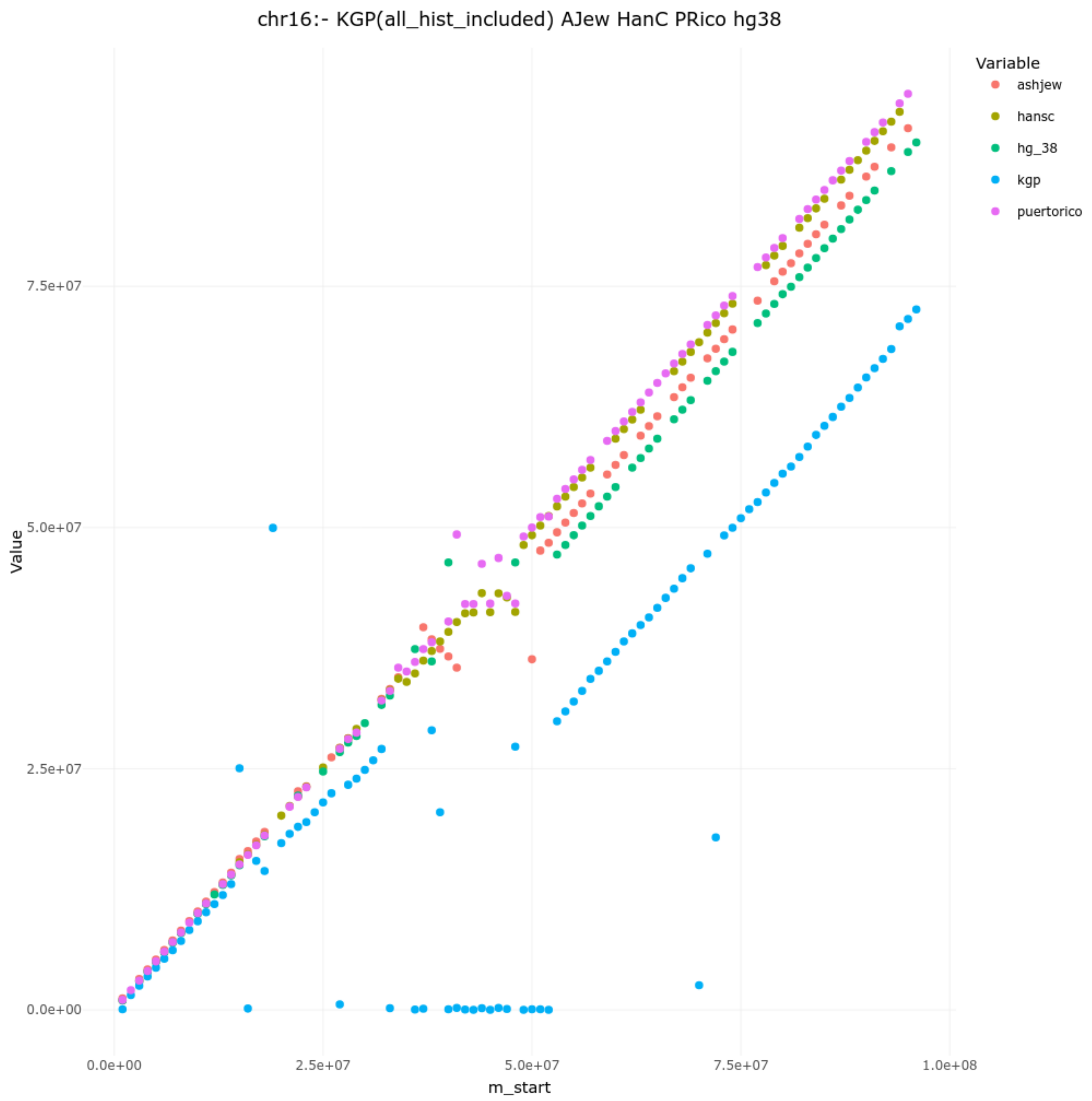
chr16:- KGP(all_hist_included) AJew HanC PRico hg38
Variable
ashjew
hansc
hg_38
kgp
puertorico
Value
7.5e+07
5.0e+07
2.5e+07
0.0e+00
0.0e+00
2.5e+07
5.0e+07
7.5e+07
1.0e+08
m_start

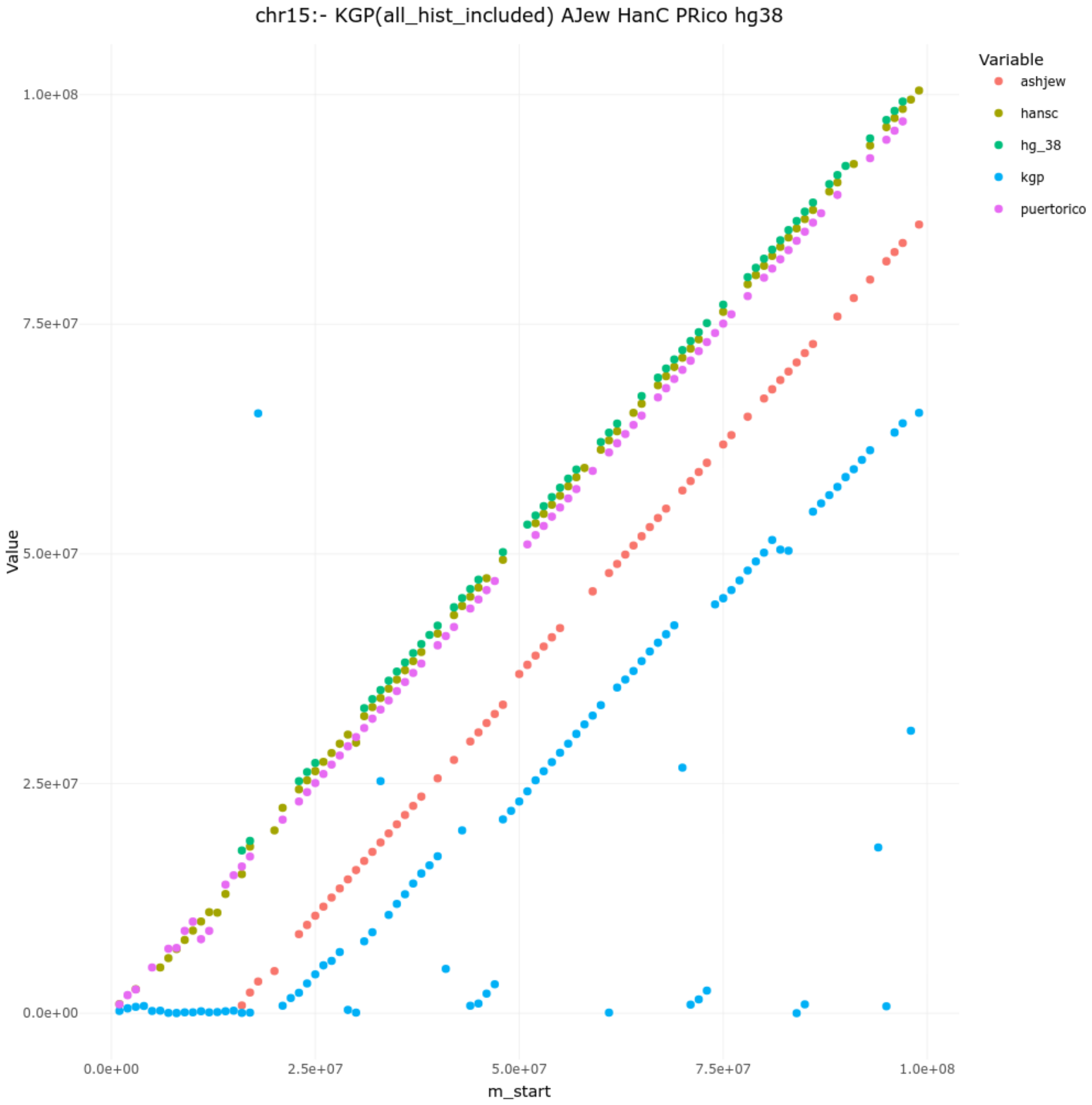

chr15:- KGP(all_hist_included) AJew HanC PRico hg38
Variable
ashjew
hansc
hg_38
kgp
puertorico
Value
1.0e+08
7.5e+07
5.0e+07
2.5e+07
0.0e+00
0.0e+00
2.5e+07
5.0e+07
7.5e+07
1.0e+08
m_start

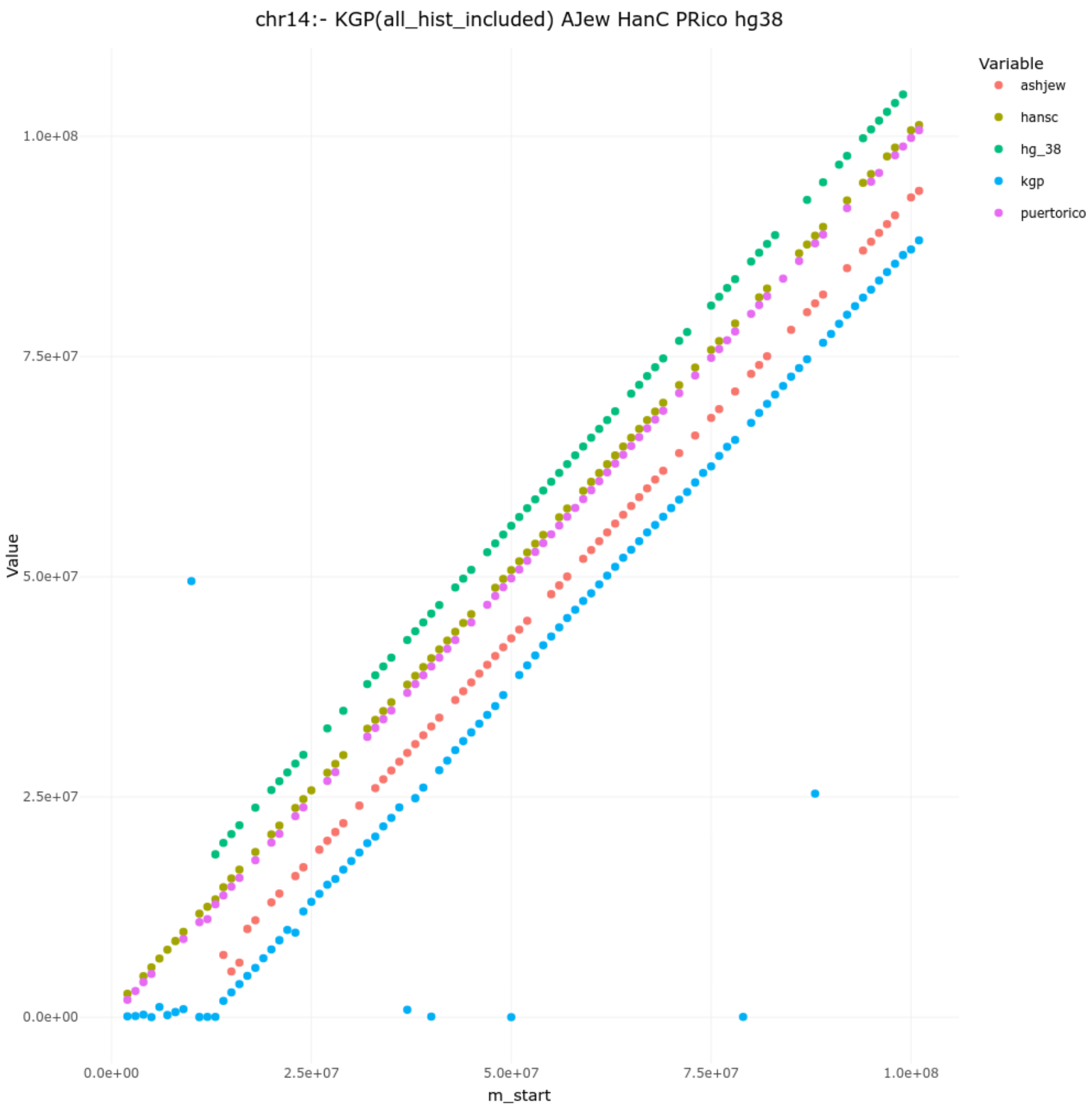
chr14:- KGP(all_hist_included) AJew HanC PRico hg38
Variable
ashjew
hansc
hg_38
kgp
puertorico
Value
1.0e+08
7.5e+07
5.0e+07
2.5e+07
0.0e+00
0.0e+00
2.5e+07
5.0e+07
7.5e+07
1.0e+08
m_start

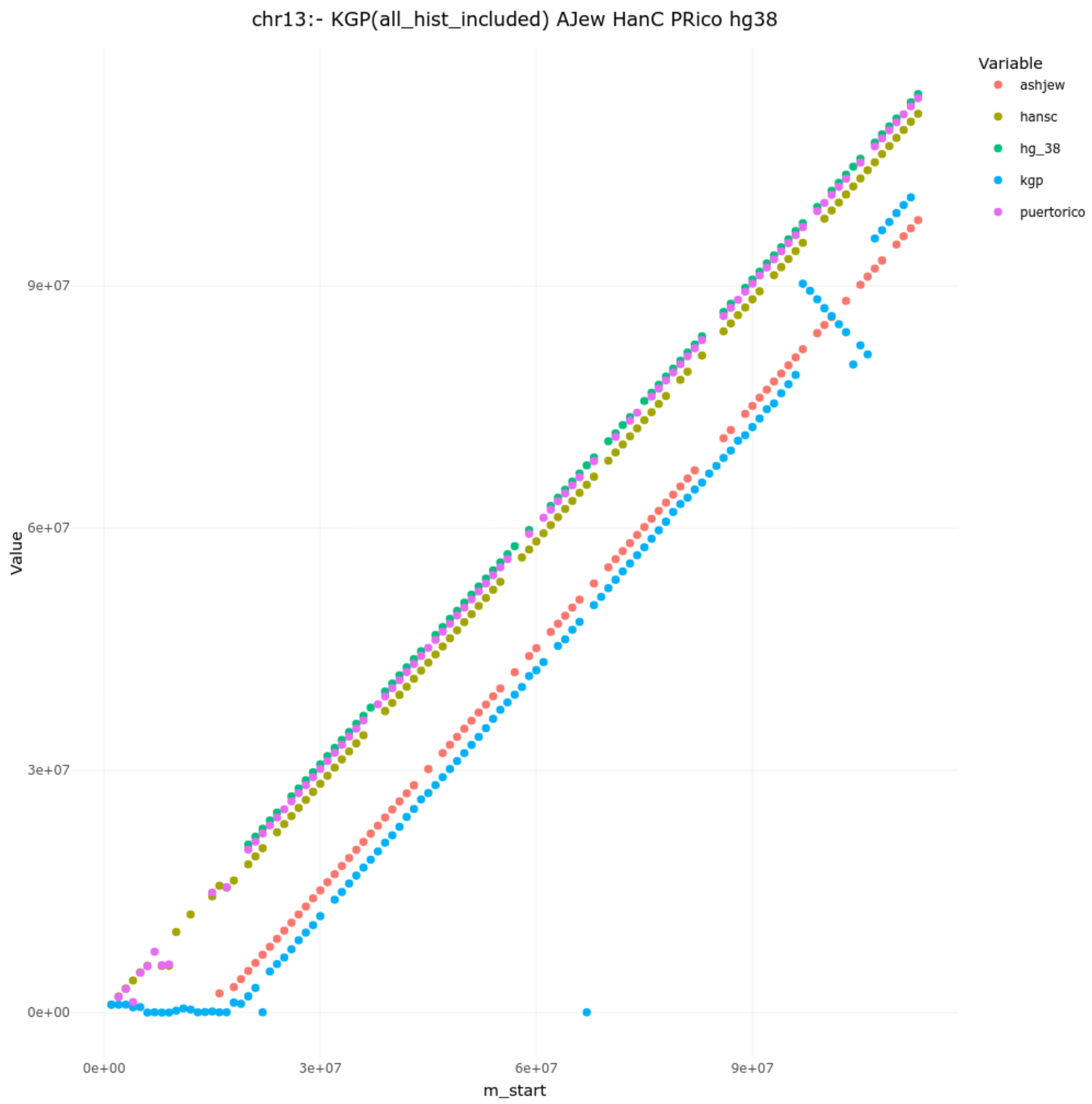

chr13:- KGP(all_hist_included) AJew HanC PRico hg38
Variable
ashjew
hansc
hg_38
kgp
puertorico
9e+07
6e+07
3e+07
0e+00
Value
0e+00
3e+07
6e+07
9e+07
m_start

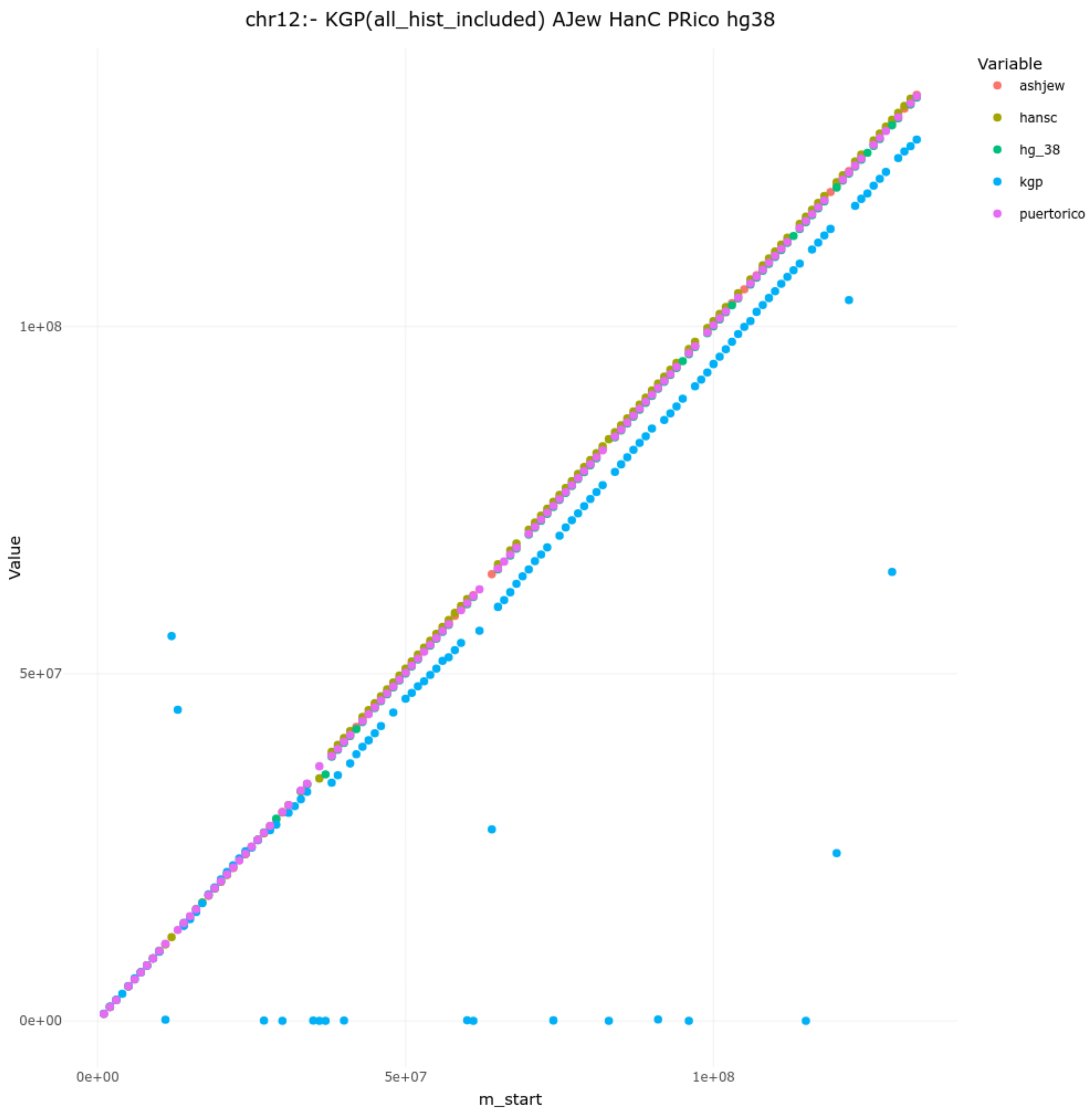

chr12:- KGP(all_hist_included) AJew HanC PRico hg38
Variable
ashjew
hansc
hg_38
kgp
puertorico
Value
1e+08
5e+07
0e+00
m_start
0e+00
5e+07
1e+08

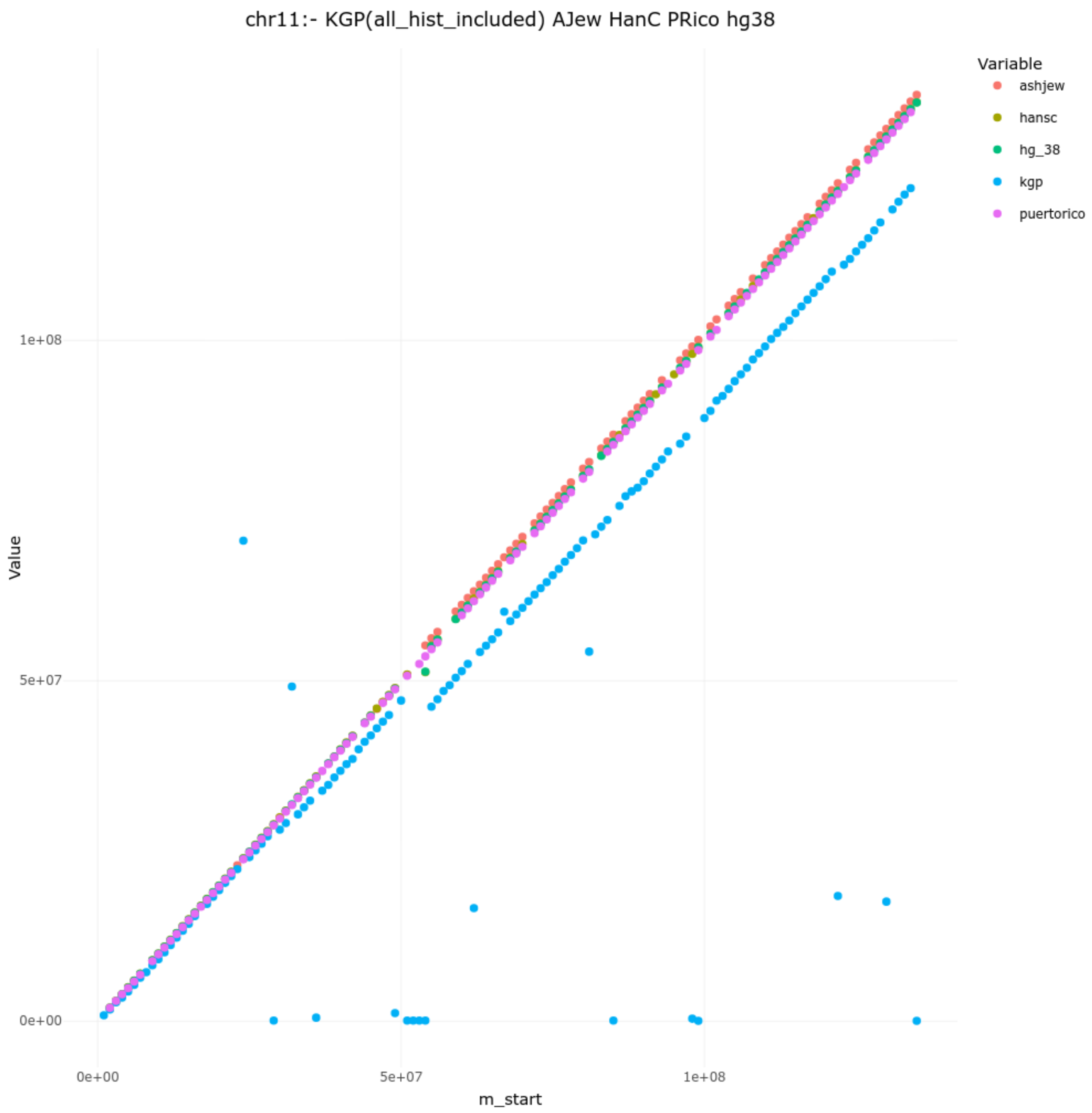

chr11:- KGP(all_hist_included) AJew HanC PRico hg38
Variable
ashjew
hansc
hg_38
kgp
puertorico
Value
1e+08
5e+07
0e+00
0e+00
5e+07
1e+08
m_start

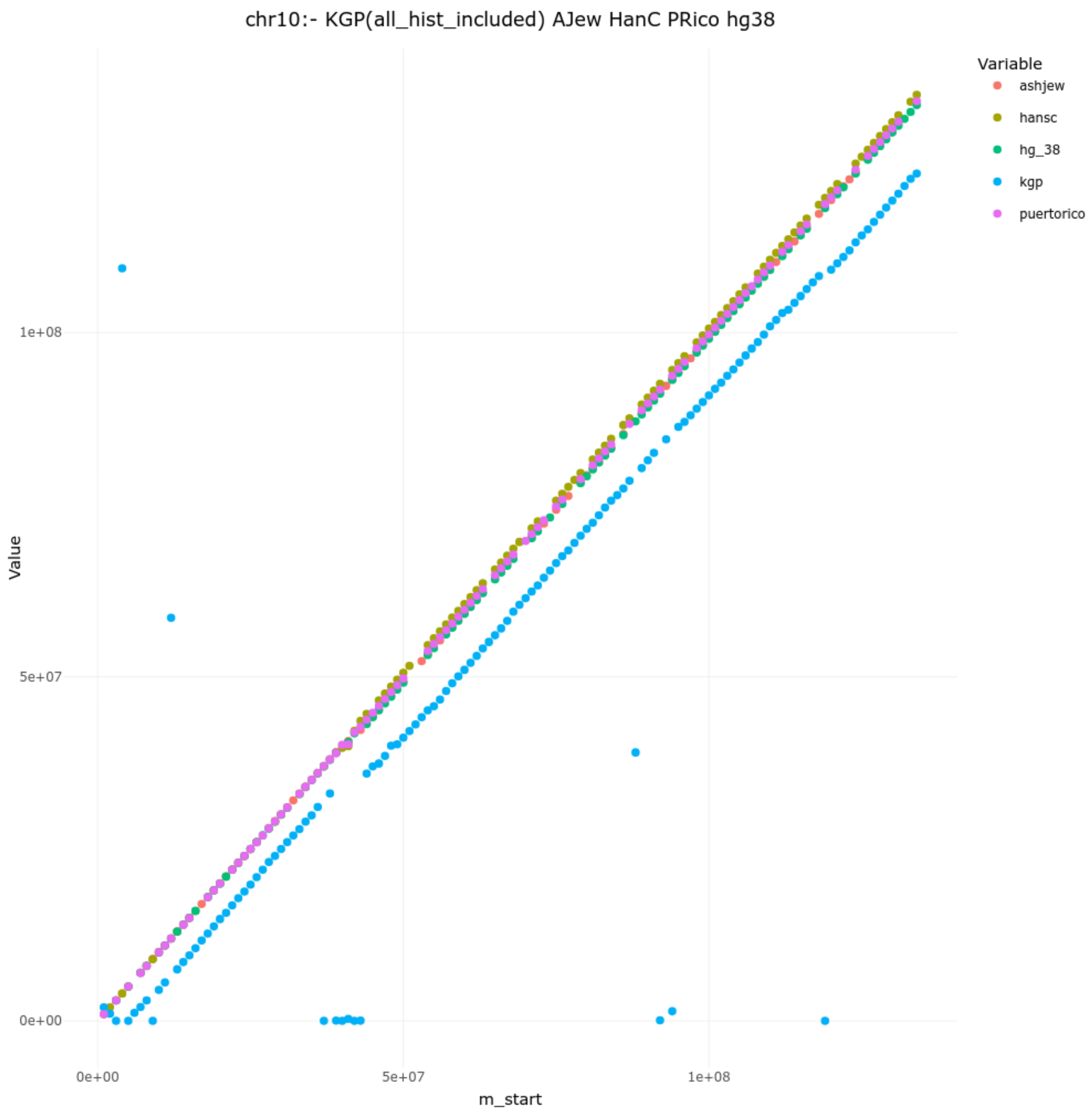

chr10:- KGP(all_hist_included) AJew HanC PRico hg38
Variable
ashjew
hansc
hg_38
kgp
puertorico
Value
1e+08
5e+07
0e+00
0e+00
5e+07
1e+08
m_start

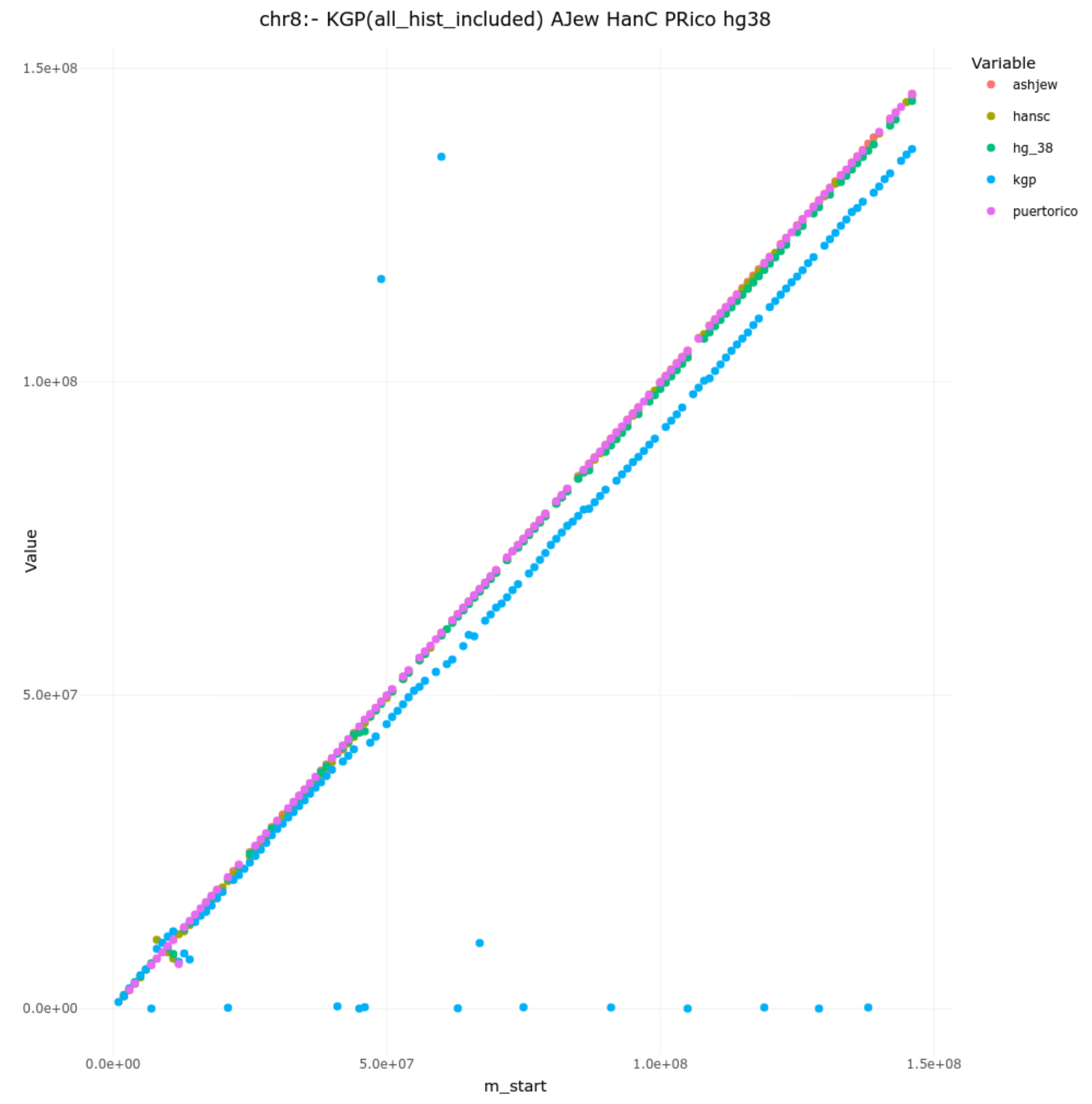
chr8:- KGP(all_hist_included) AJew HanC PRico hg38
Value
1.5e+08
1.0e+08
5.0e+07
0.0e+00
0.0e+00
5.0e+07
1.0e+08
1.5e+08
m_start
Variable
ashjew
hansc
hg_38
kgp
puertorico

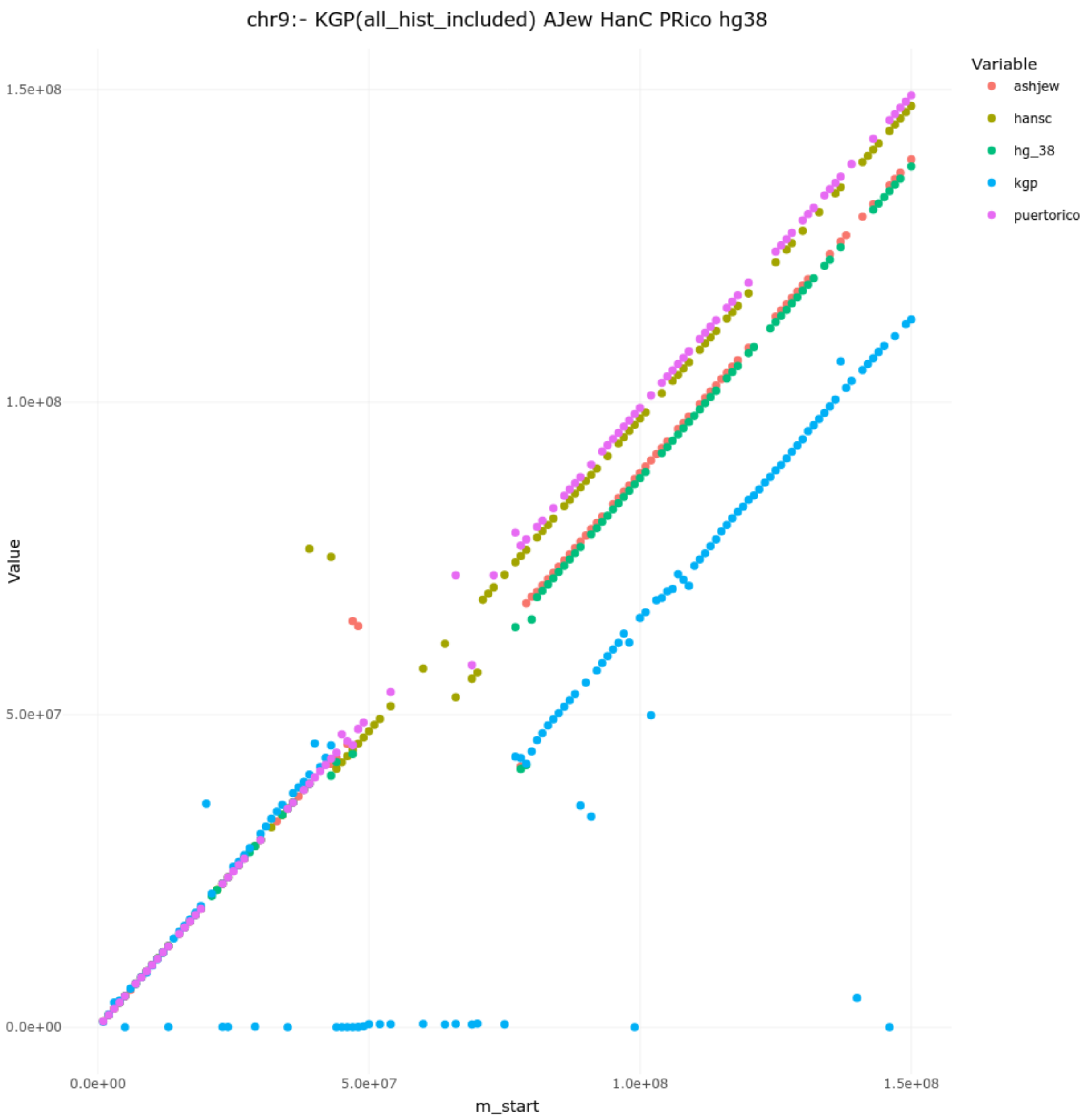
chr9:- KGP(all_hist_included) AJew HanC PRico hg38
Variable
ashjew
hansc
hg_38
kgp
puertorico
Value
1.5e+08
1.0e+08
5.0e+07
0.0e+00
0.0e+00
5.0e+07
1.0e+08
1.5e+08
m_start

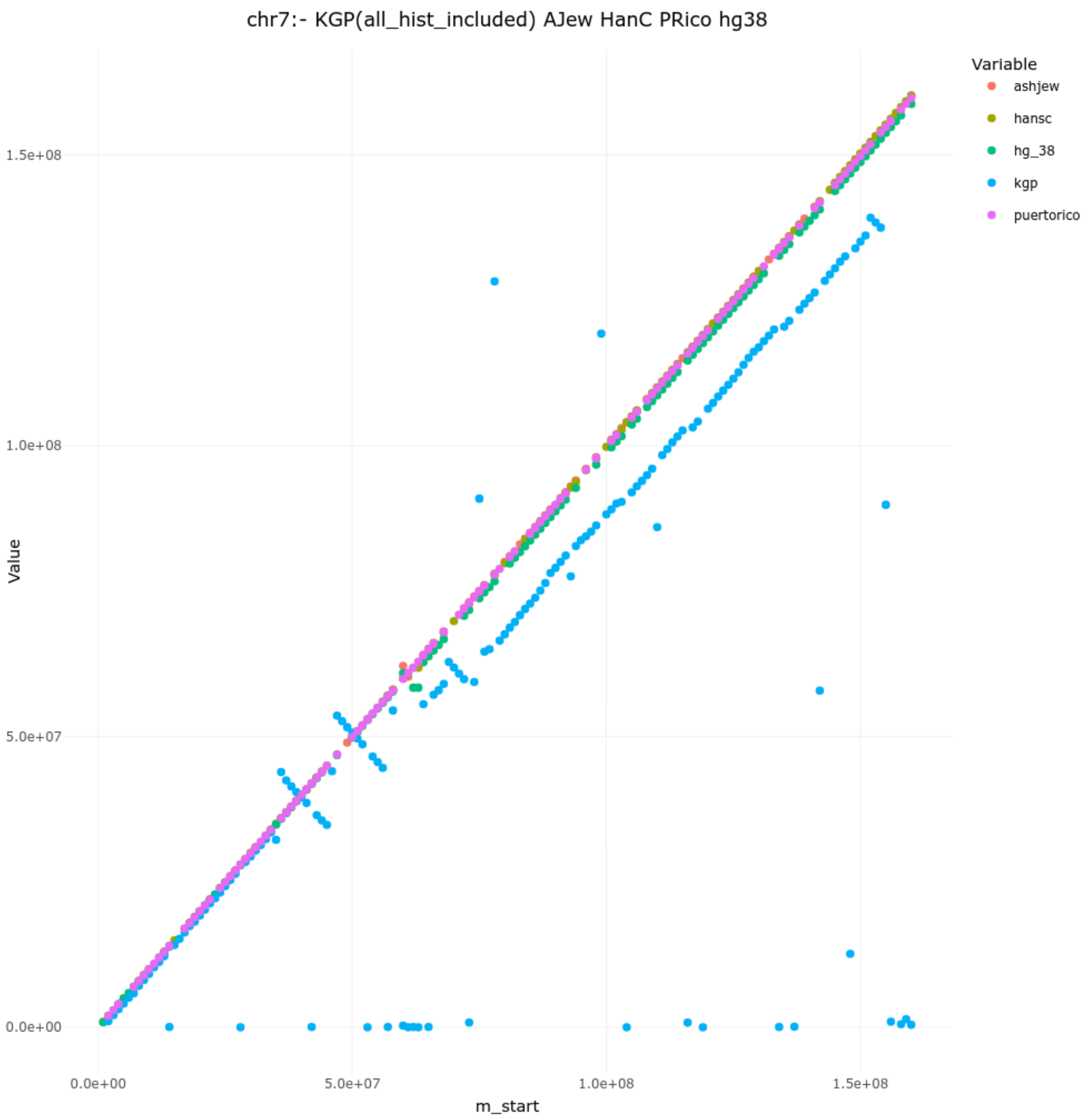
chr7:- KGP(all_hist_included) AJew HanC PRico hg38
Variable
ashjew
hansc
hg_38
kgp
puertorico
Value
1.5e+08
1.0e+08
5.0e+07
0.0e+00
m_start
0.0e+00
5.0e+07
1.0e+08
1.5e+08

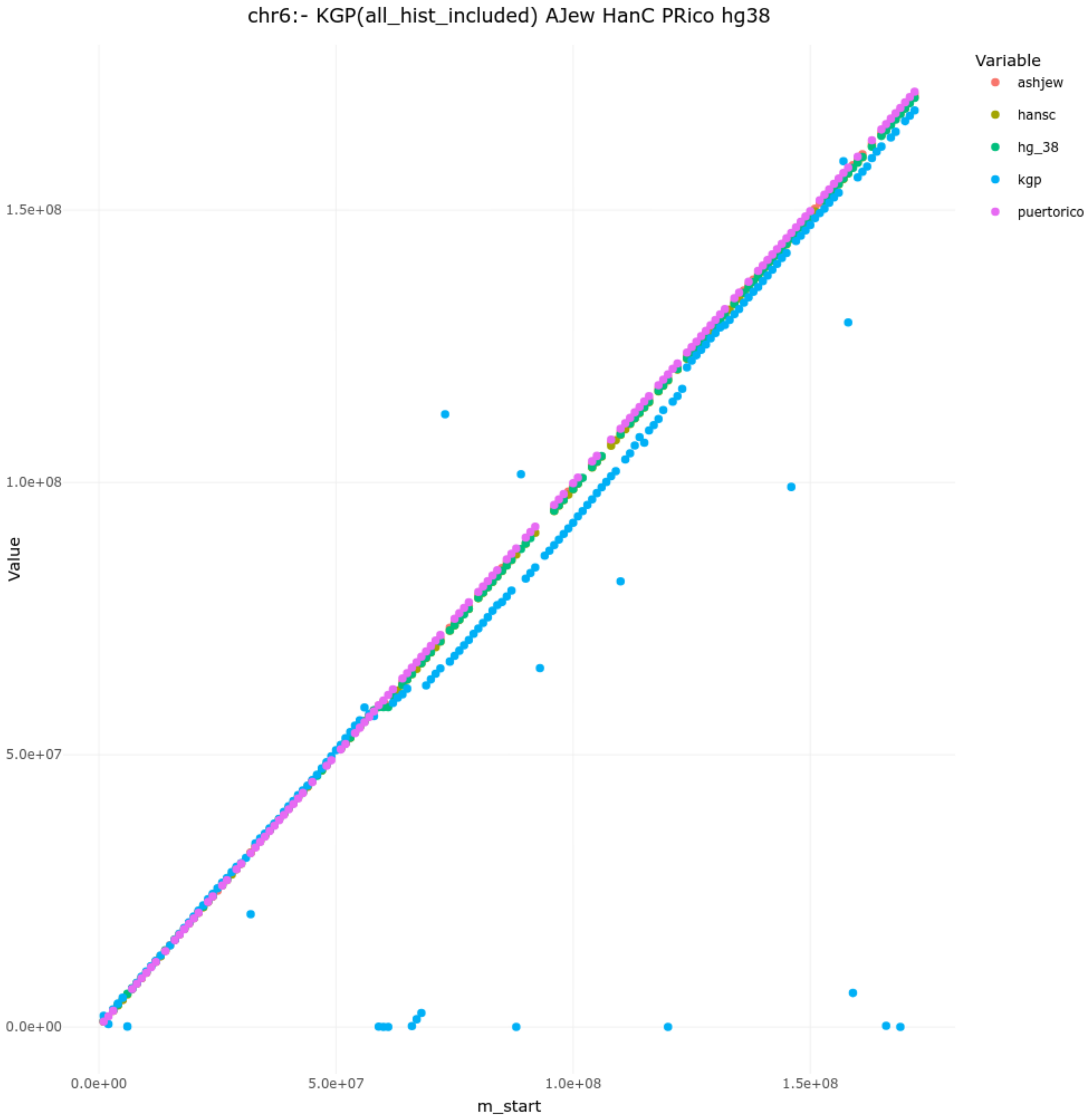
chr6:- KGP(all_hist_included) AJew HanC PRico hg38
Variable
ashjew
hansc
hg_38
kgp
puertorico
1.5e+08
1.0e+08
5.0e+07
0.0e+00
Value
0.0e+00
5.0e+07
1.0e+08
1.5e+08
m_start

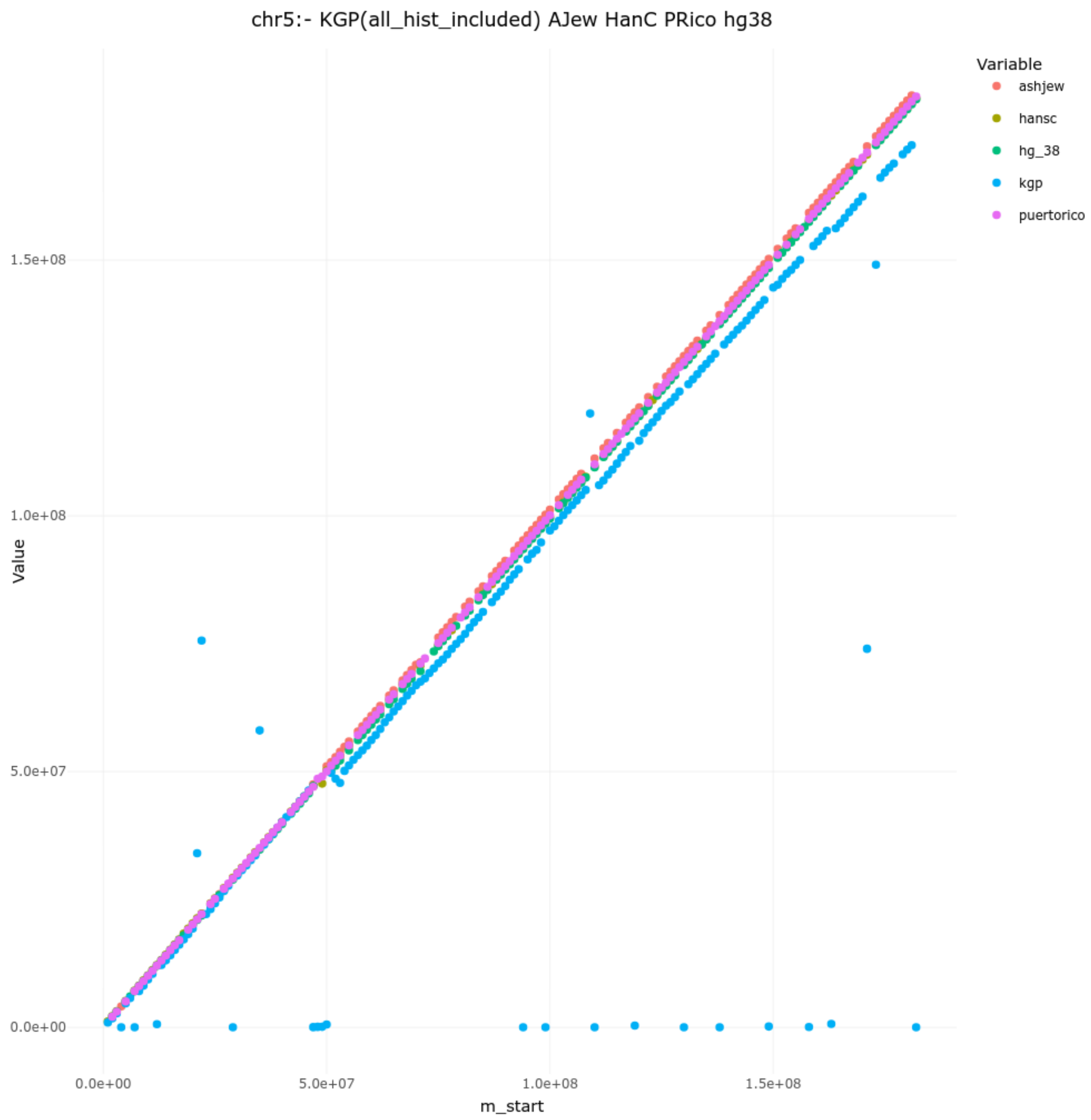

chr5:- KGP(all_hist_included) AJew HanC PRico hg38
Variable
ashjew
hansc
hg_38
kgp
puertorico
Value
1.5e+08
1.0e+08
5.0e+07
0.0e+00
0.0e+00
5.0e+07
1.0e+08
1.5e+08
m_start

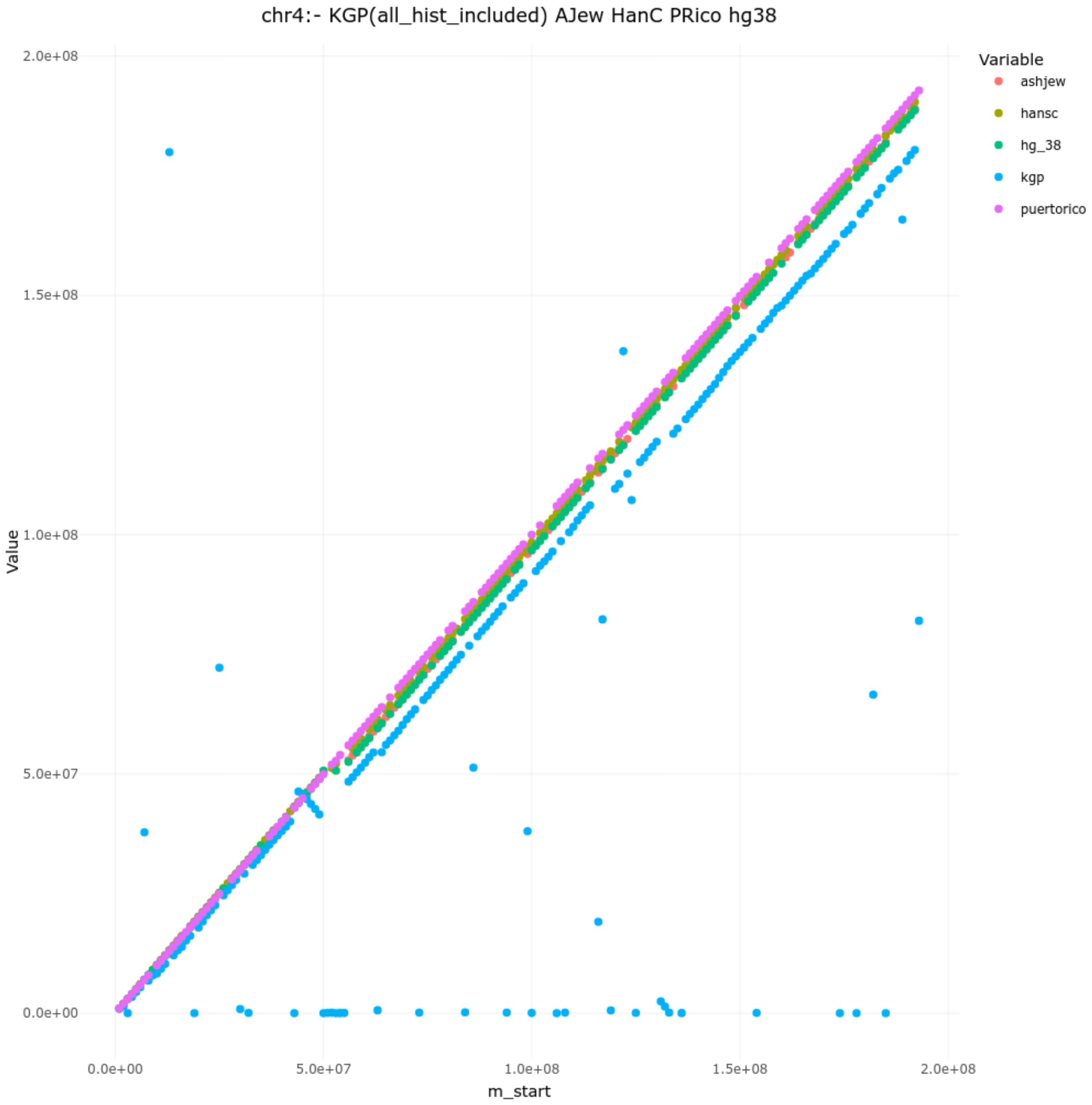
chr4:- KGP(all_hist_included) AJew HanC PRico hg38
Value
m_start
2.0e+08
1.5e+08
1.0e+08
5.0e+07
0.0e+00
0.0e+00
5.0e+07
1.0e+08
1.5e+08
2.0e+08
Variable
ashjew
hansc
hg_38
kgp
puertorico

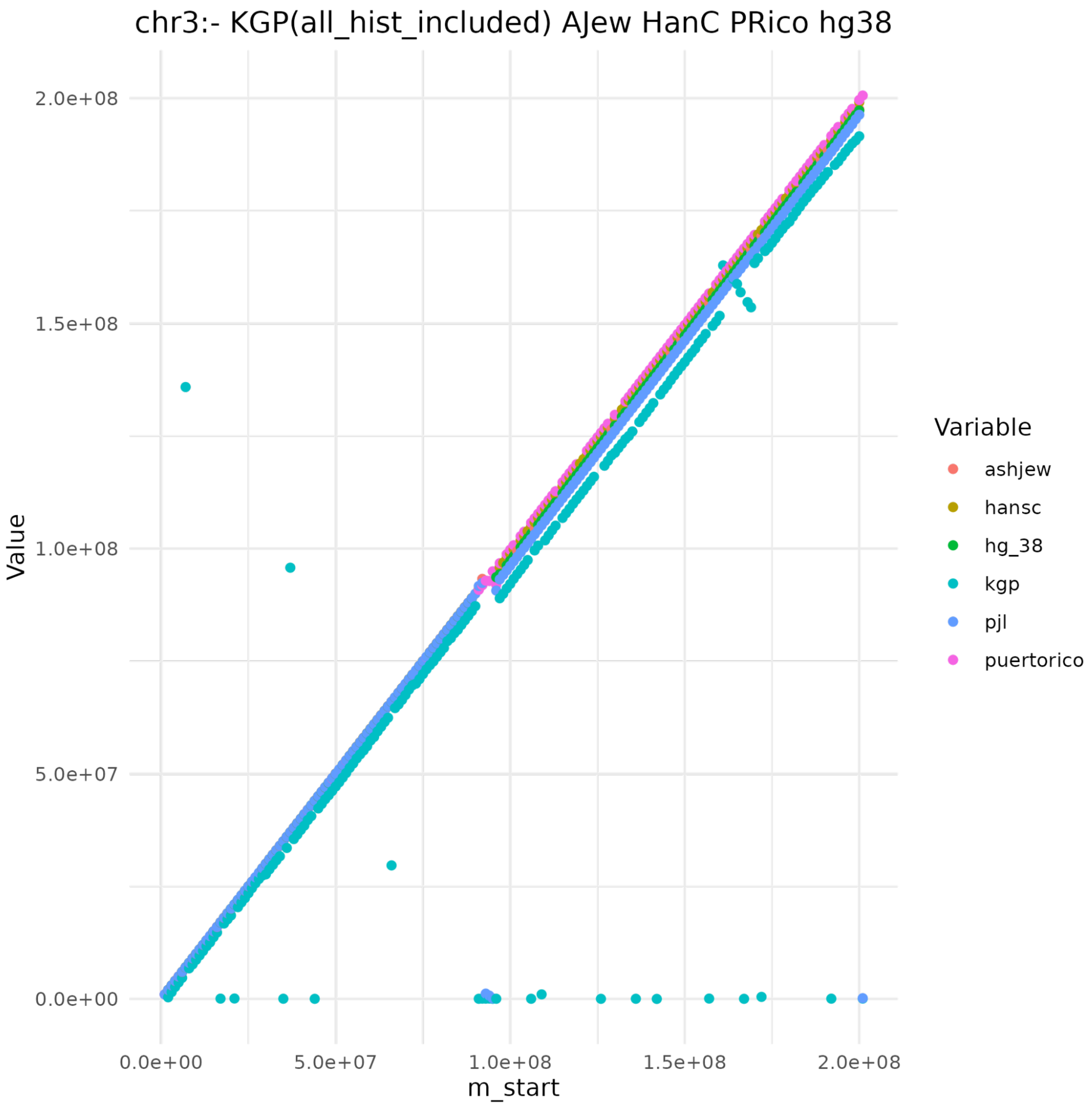

chr3:- KGP(all_hist_included) AJew HanC PRico hg38
Value
2.0e+08
1.5e+08
1.0e+08
5.0e+07
0.0e+00
0.0e+00
5.0e+07
1.0e+08
1.5e+08
2.0e+08
m_start
Variable
ashjew
hansc
hg_38
kgp
pjl
puertorico

-

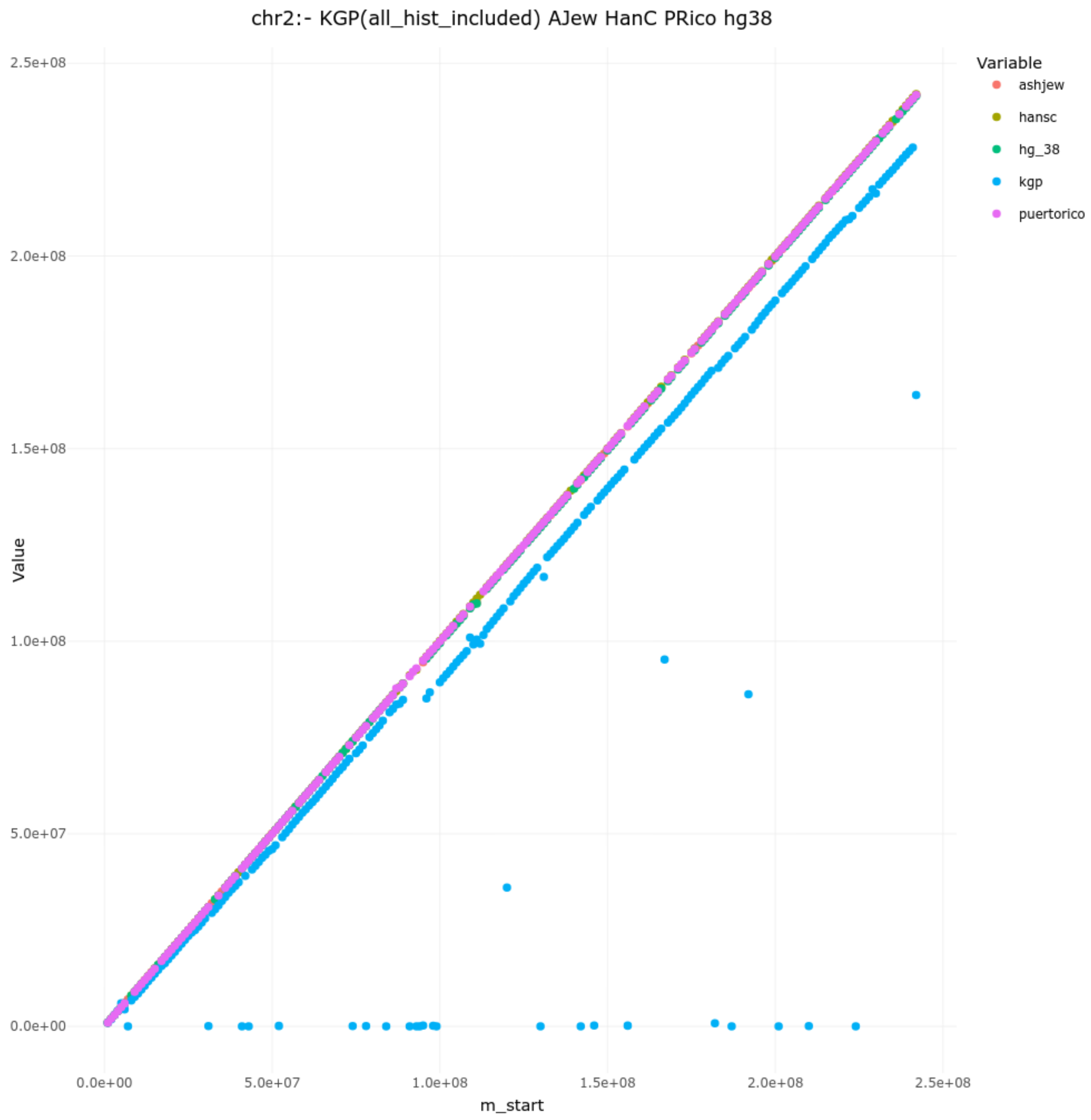

chr2:- KGP(all_hist_included) AJew HanC PRico hg38
Value
m_start
0.0e+00
5.0e+07
1.0e+08
1.5e+08
2.0e+08
2.5e+08
Variable
ashjew
hansc
hg_38
kgp
puertorico

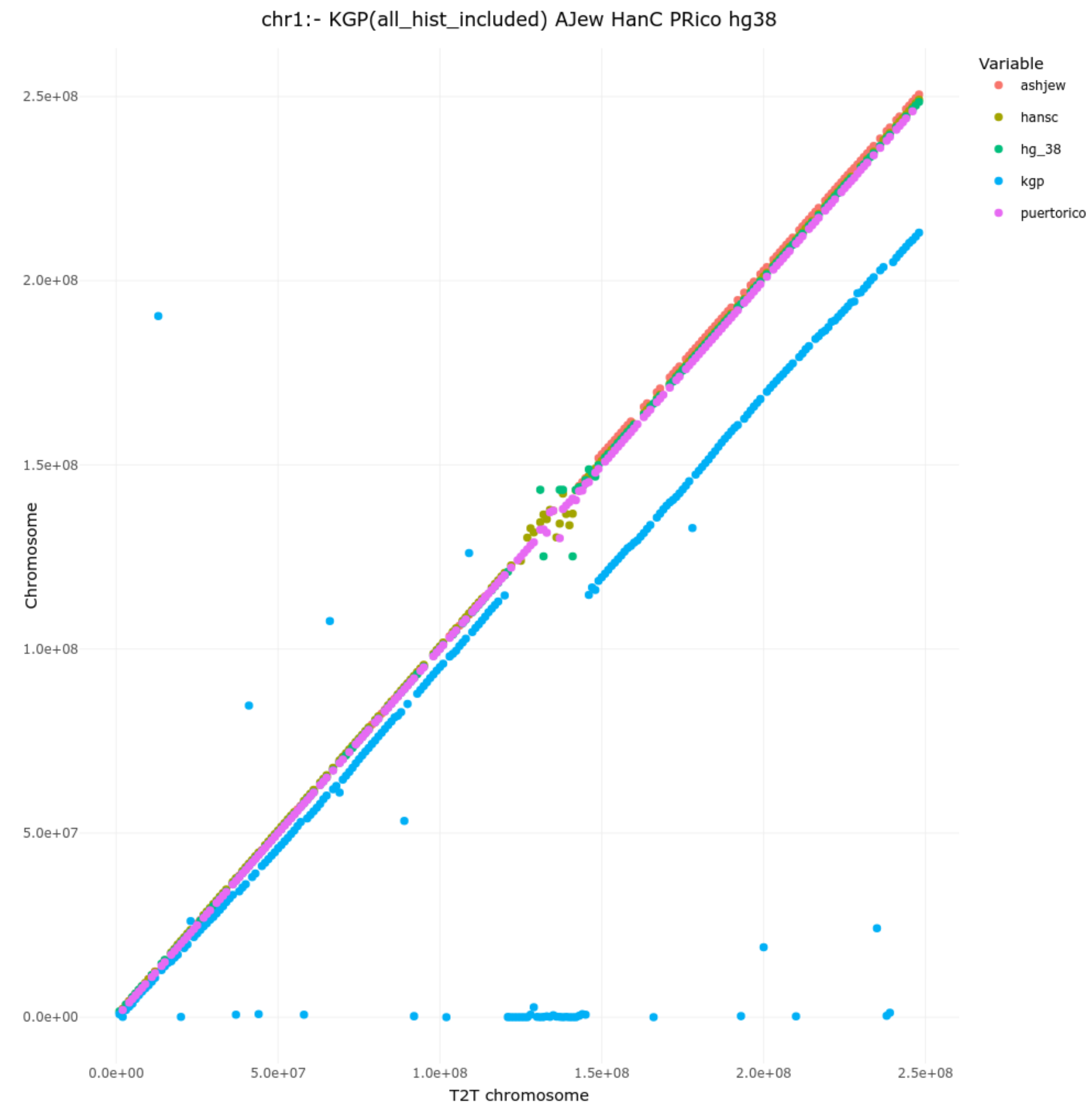
chr1:- KGP(all_hist_included) AJew HanC PRico hg38
Variable
ashjew
hansc
hg_38
kgp
puertorico
Chromosome
T2T chromosome
0.0e+00
5.0e+07
1.0e+08
1.5e+08
2.0e+08
2.5e+08

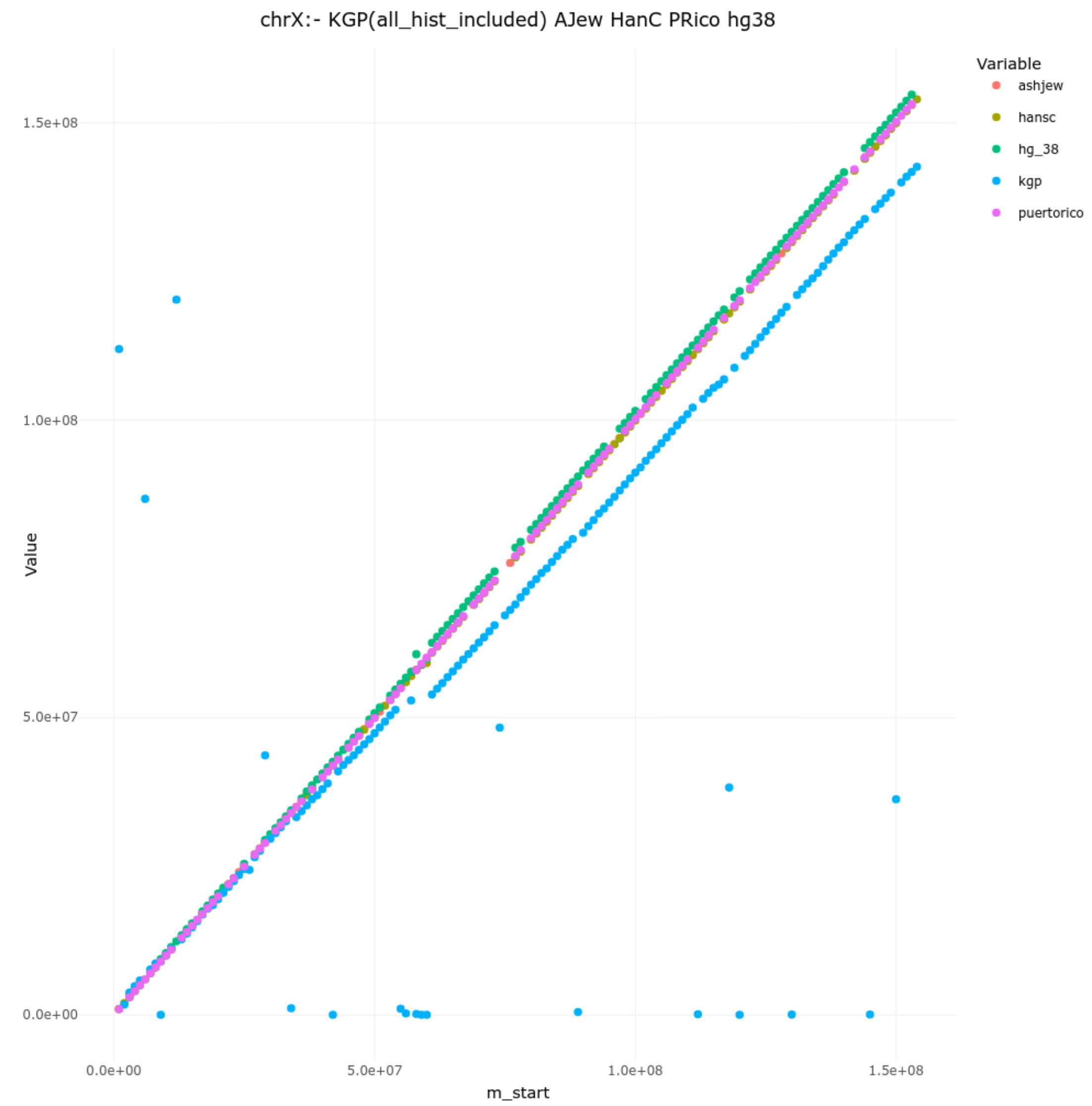


Supplementary Figure 1: Comparative dot plot of chromosomes depicting linearity of genomes of PR1, Ash1, Han1, hg38 and T2T..

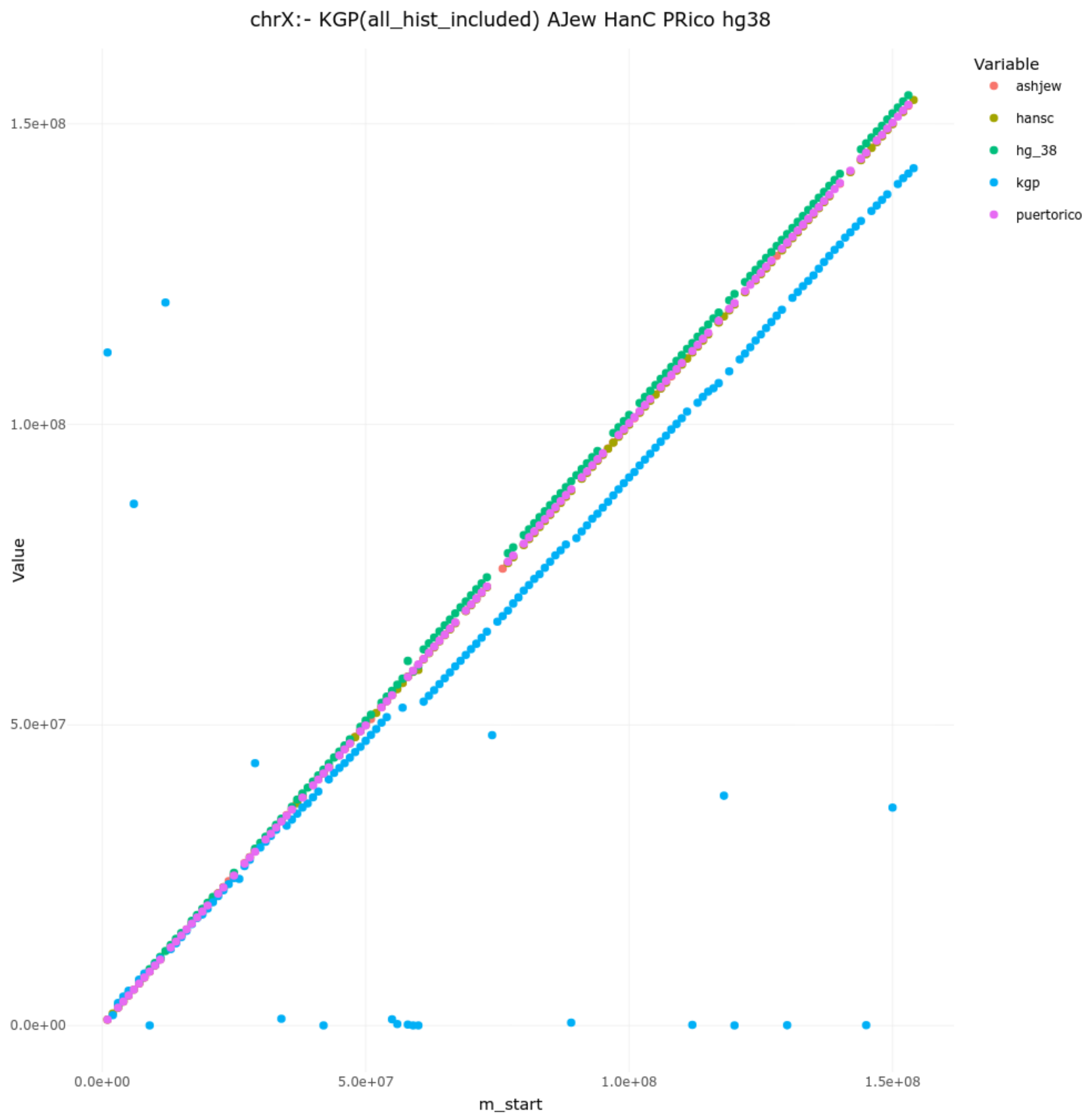
chrX:- KGP(all_hist_included) AJew HanC PRico hg38
Variable
ashjew
hansc
hg_38
kgp
puertorico
Value
1.5e+08
1.0e+08
5.0e+07
0.0e+00
0.0e+00
5.0e+07
1.0e+08
1.5e+08
m_start

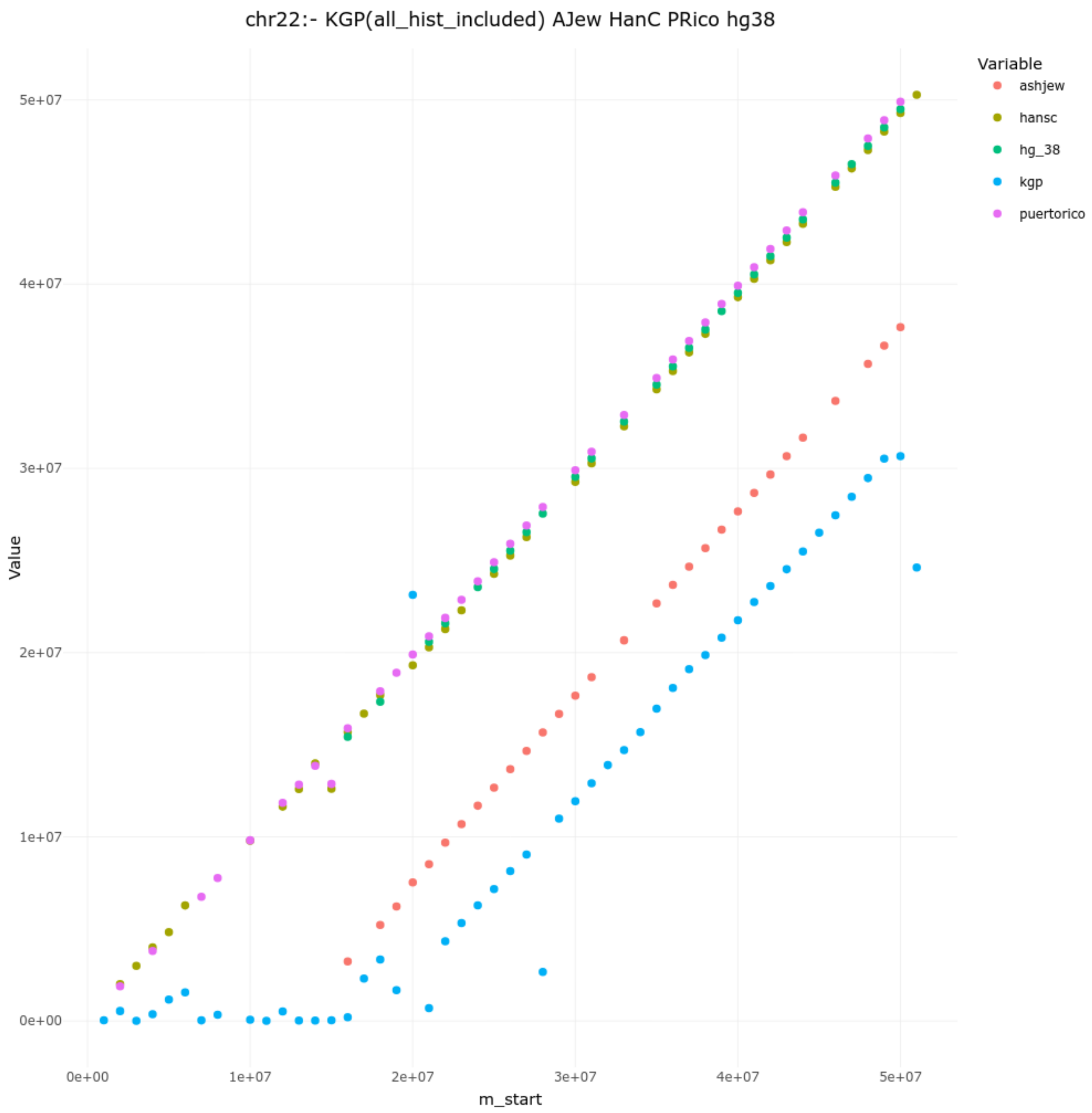

chr22:- KGP(all_hist_included) AJew HanC PRico hg38
Variable
ashjew
hansc
hg_38
kgp
puertorico
Value
5e+07
4e+07
3e+07
2e+07
1e+07
0e+00
0e+00
1e+07
2e+07
3e+07
4e+07
5e+07
m_start

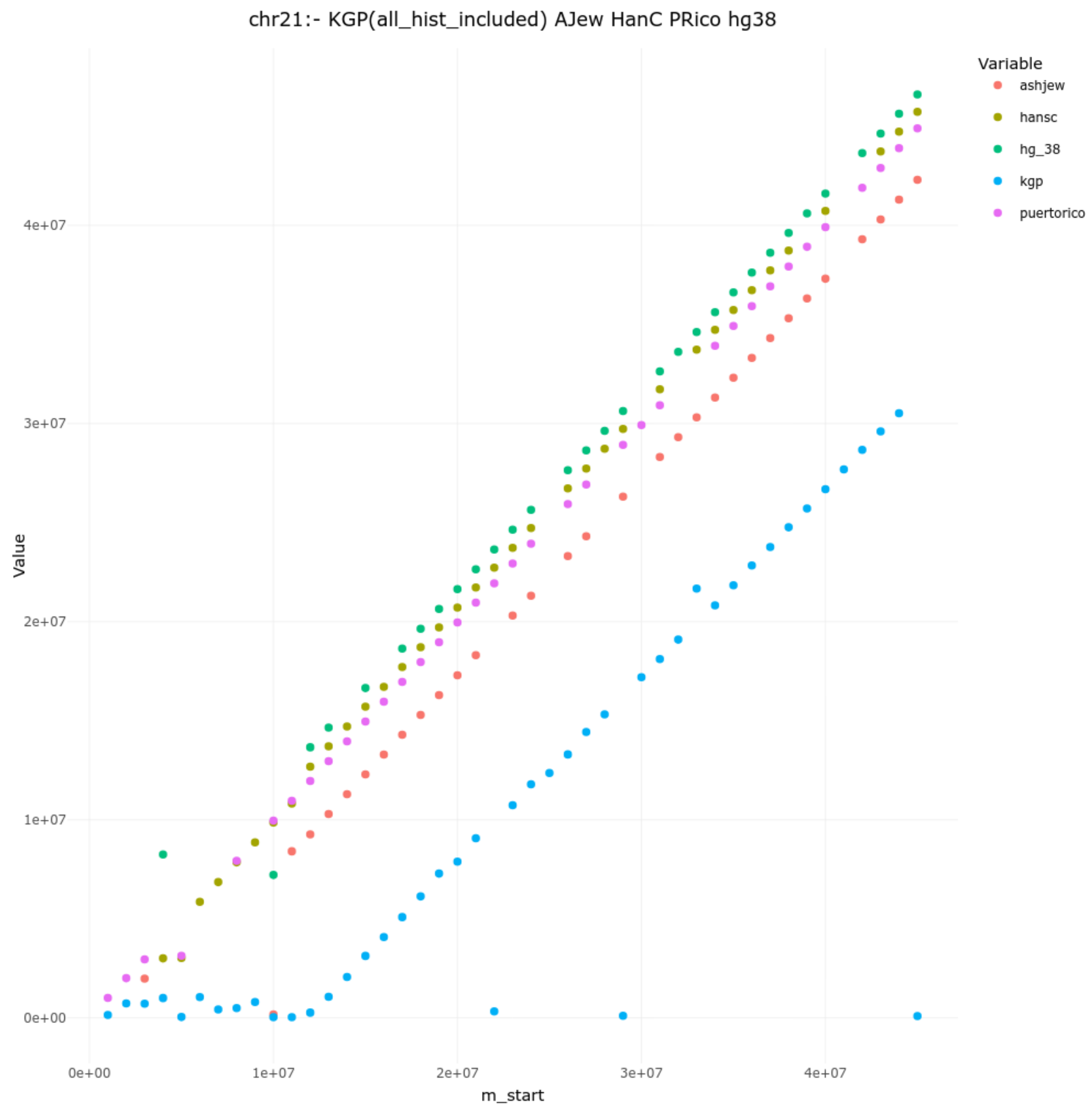

chr21:- KGP(all_hist_included) AJew HanC PRico hg38
Variable
ashjew
hansc
hg_38
kgp
puertorico
Value
4e+07
3e+07
2e+07
1e+07
0e+00
0e+00
1e+07
2e+07
3e+07
4e+07
m_start

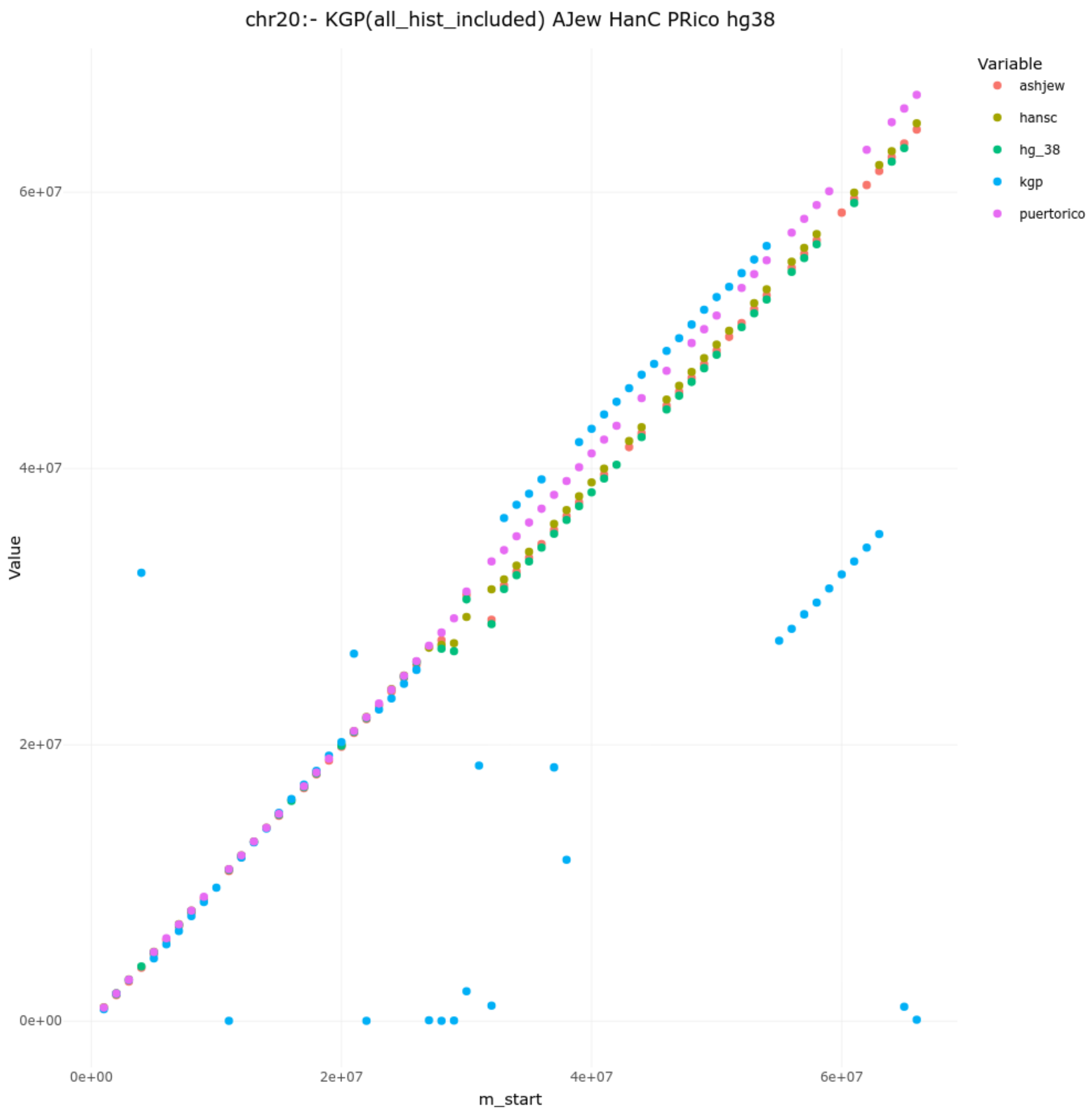
chr20:- KGP(all_hist_included) AJew HanC PRico hg38
Variable
ashjew
hansc
hg_38
kgp
puertorico
Value
6e+07
4e+07
2e+07
0e+00
0e+00
2e+07
4e+07
6e+07
m_start

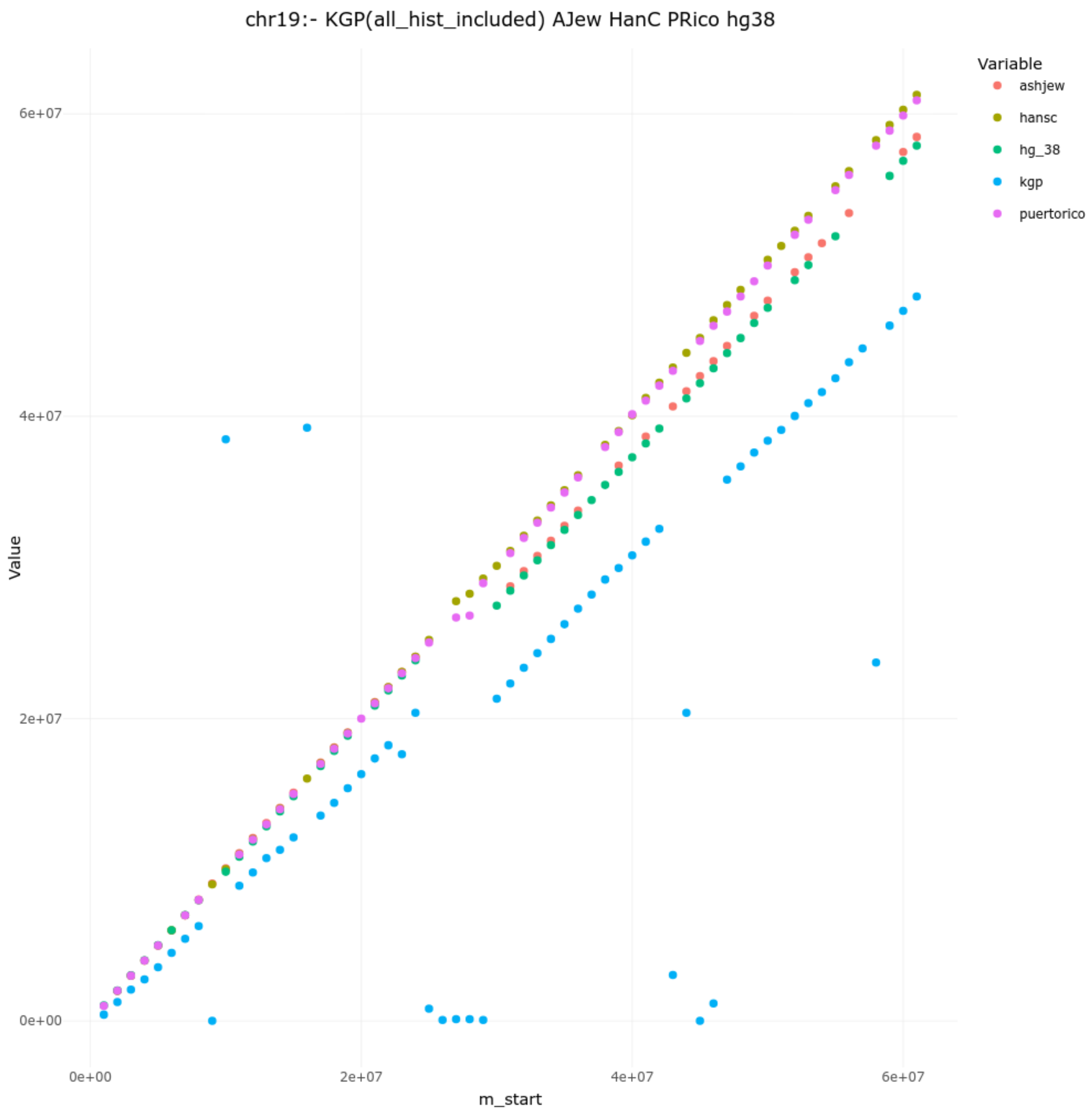
chr19:- KGP(all_hist_included) AJew HanC PRico hg38
Value
m_start
0e+00
2e+07
4e+07
6e+07
0e+00
2e+07
4e+07
6e+07
Variable
ashjew
hansc
hg_38
kgp
puertorico

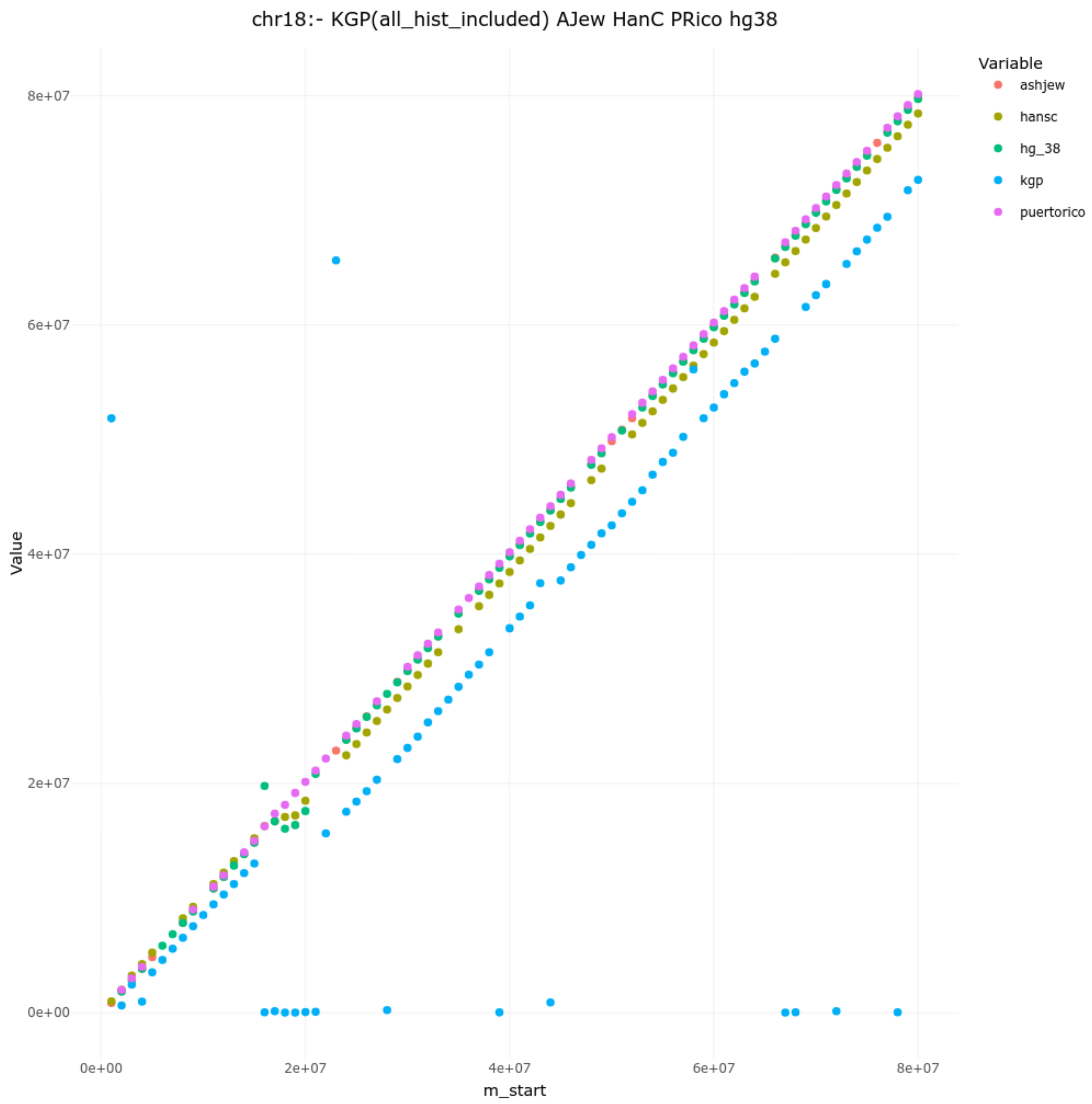
chr18:- KGP(all_hist_included) AJew HanC PRico hg38
Variable
ashjew
hansc
hg_38
kgp
puertorico
Value
8e+07
6e+07
4e+07
2e+07
0e+00
0e+00
2e+07
4e+07
6e+07
8e+07
m_start

CHR17

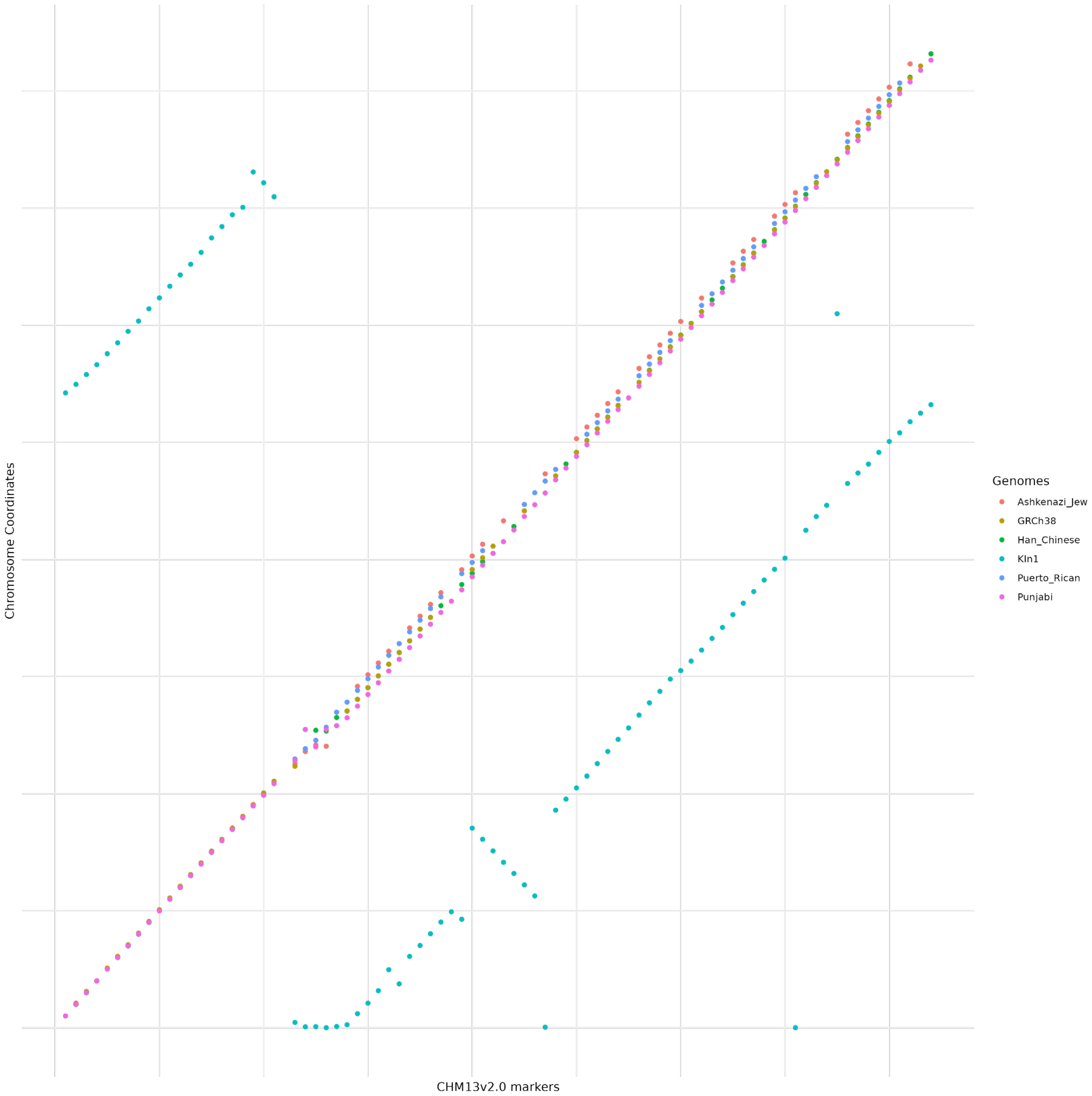
Chromosome Coordinates
CHM13v2.0 markers
Genomes
Ashkenazi_Jew
GRCh38
Han_Chinese
KIn1
Puerto_Rican
Punjabi

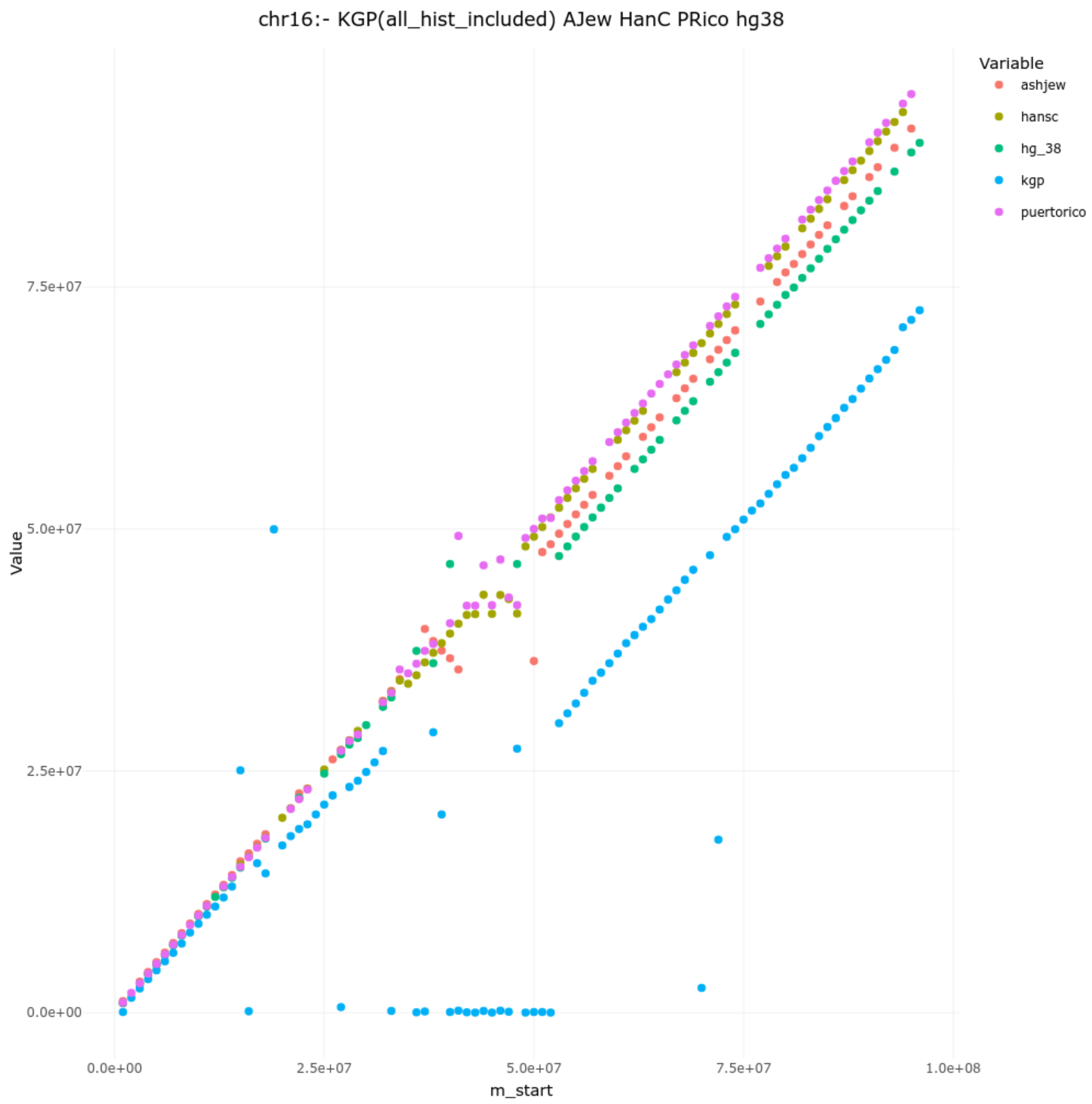

chr16:- KGP(all_hist_included) AJew HanC PRico hg38
Variable
ashjew
hansc
hg_38
kgp
puertorico
Value
7.5e+07
5.0e+07
2.5e+07
0.0e+00
0.0e+00
2.5e+07
5.0e+07
7.5e+07
1.0e+08
m_start

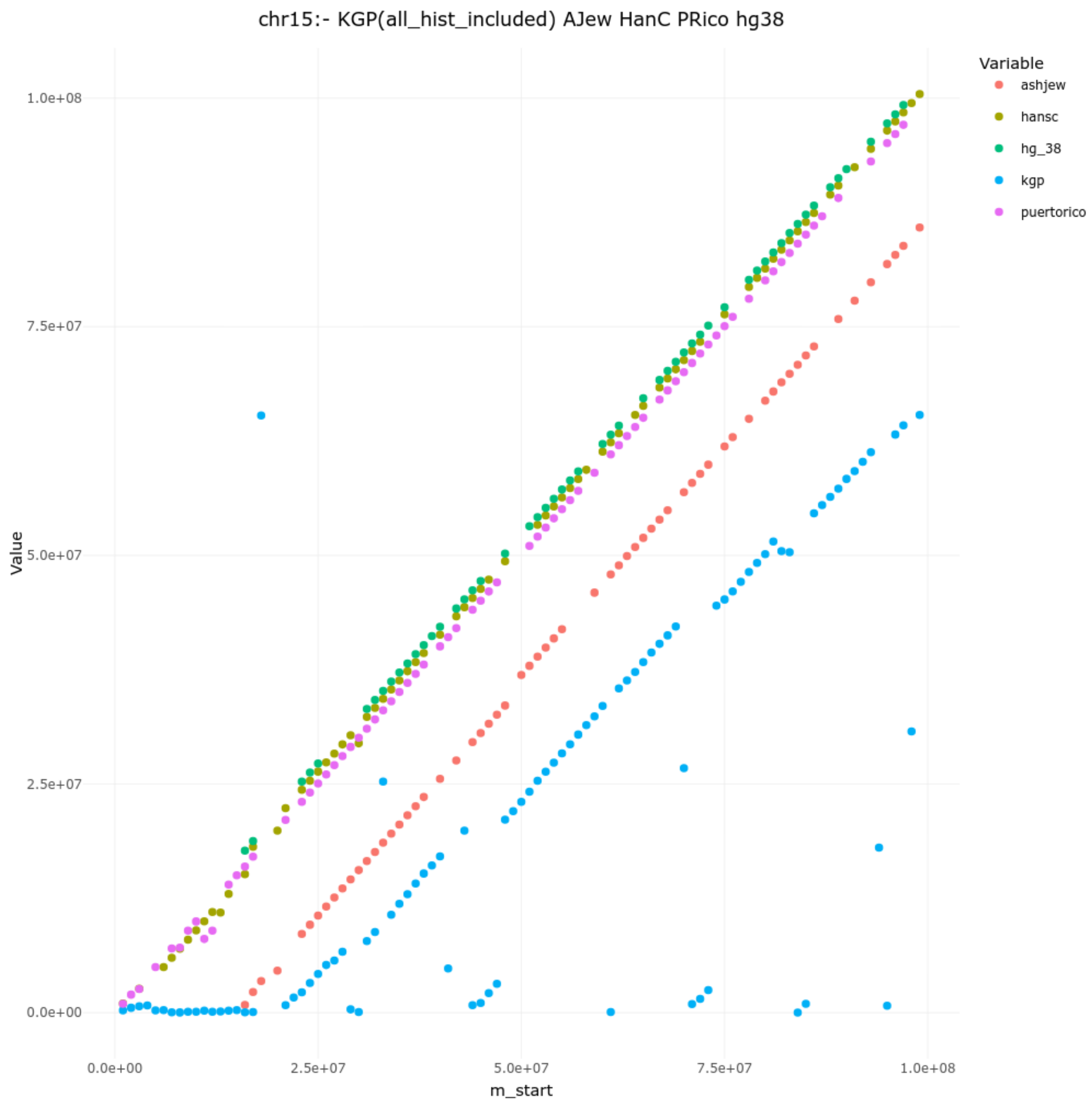
chr15:- KGP(all_hist_included) AJew HanC PRico hg38
Value
1.0e+08
7.5e+07
5.0e+07
2.5e+07
0.0e+00
0.0e+00
2.5e+07
5.0e+07
7.5e+07
1.0e+08
m_start
Variable
ashjew
hansc
hg_38
kgp
puertorico

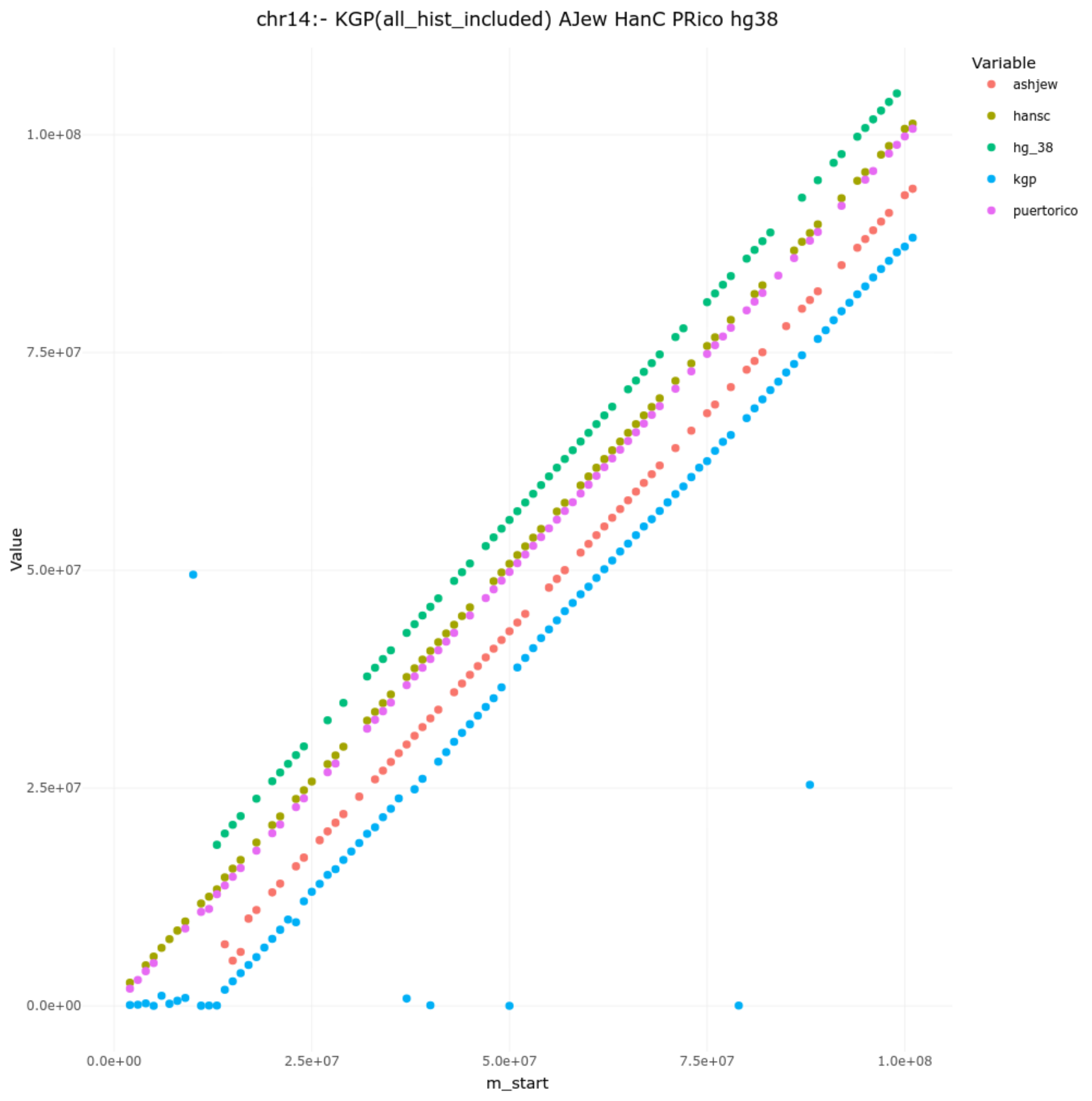

chr14:- KGP(all_hist_included) AJew HanC PRico hg38
Variable
ashjew
hansc
hg_38
kgp
puertorico
Value
1.0e+08
7.5e+07
5.0e+07
2.5e+07
0.0e+00
0.0e+00
2.5e+07
5.0e+07
7.5e+07
1.0e+08
m_start

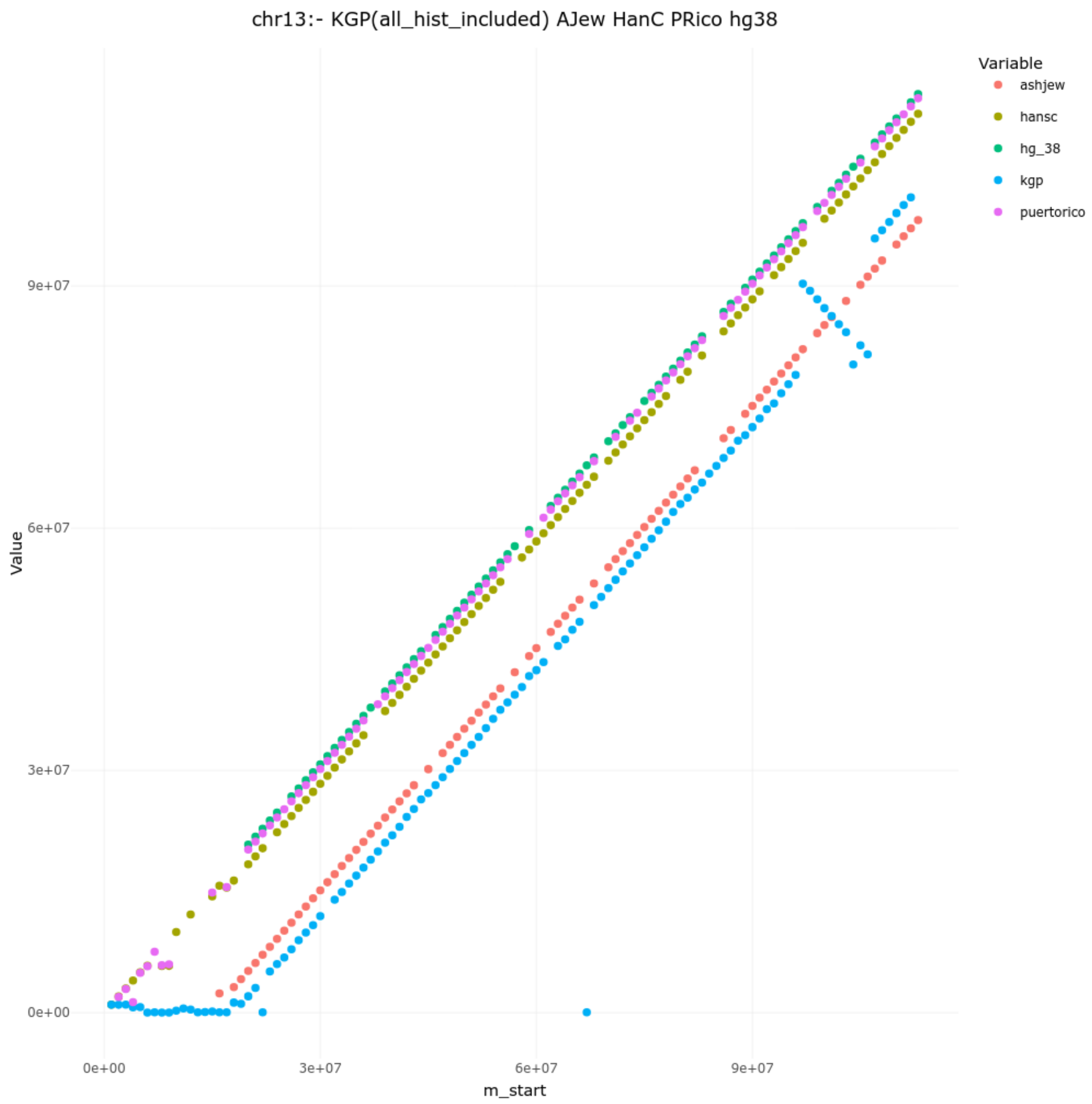

chr13:- KGP(all_hist_included) AJew HanC PRico hg38
Variable
ashjew
hansc
hg_38
kgp
puertorico
Value
9e+07
6e+07
3e+07
0e+00
0e+00
3e+07
6e+07
9e+07
m_start

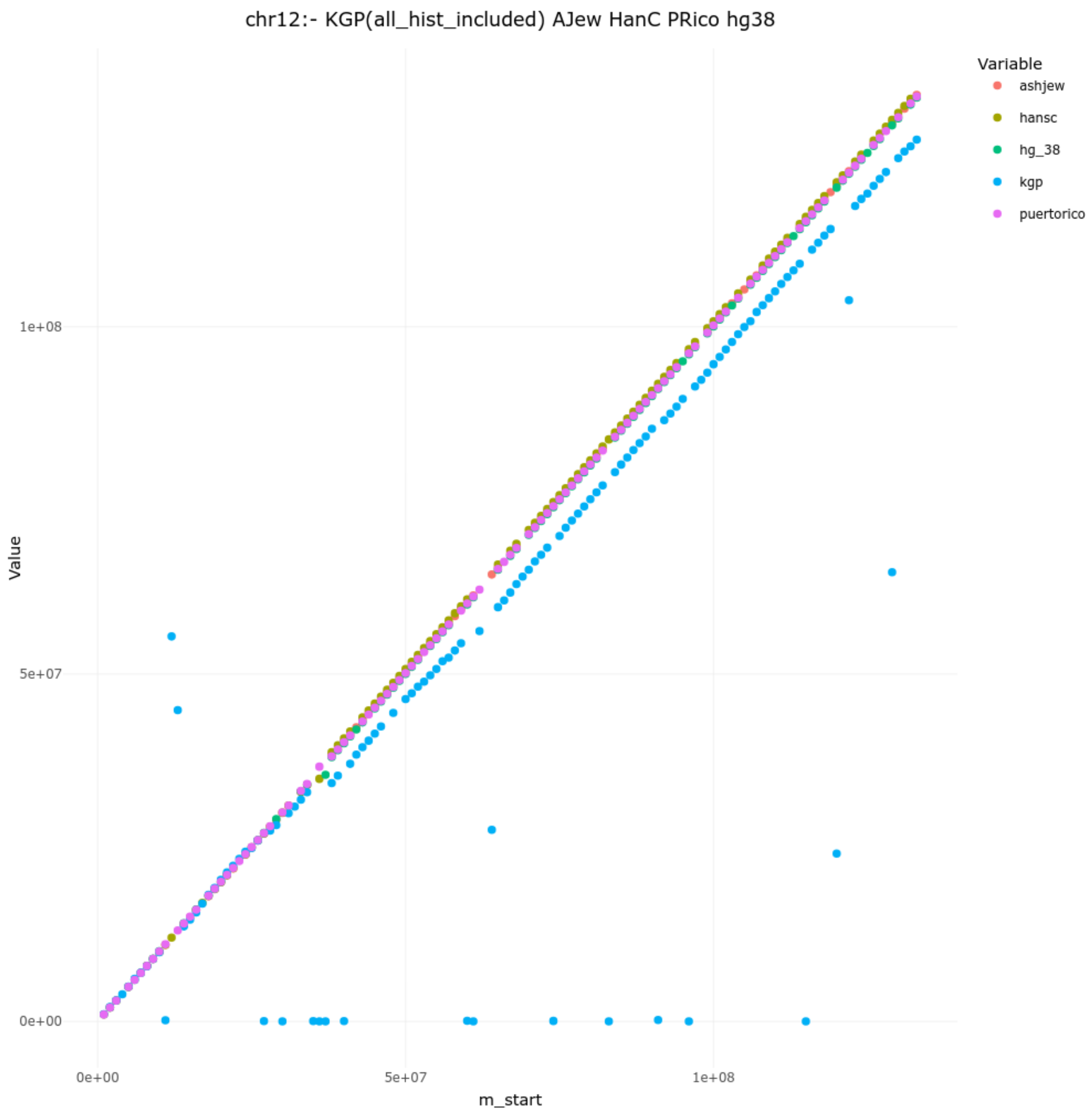
chr12:- KGP(all_hist_included) AJew HanC PRico hg38
Variable
ashjew
hansc
hg_38
kgp
puertorico
Value
1e+08
5e+07
0e+00
m_start
0e+00
5e+07
1e+08

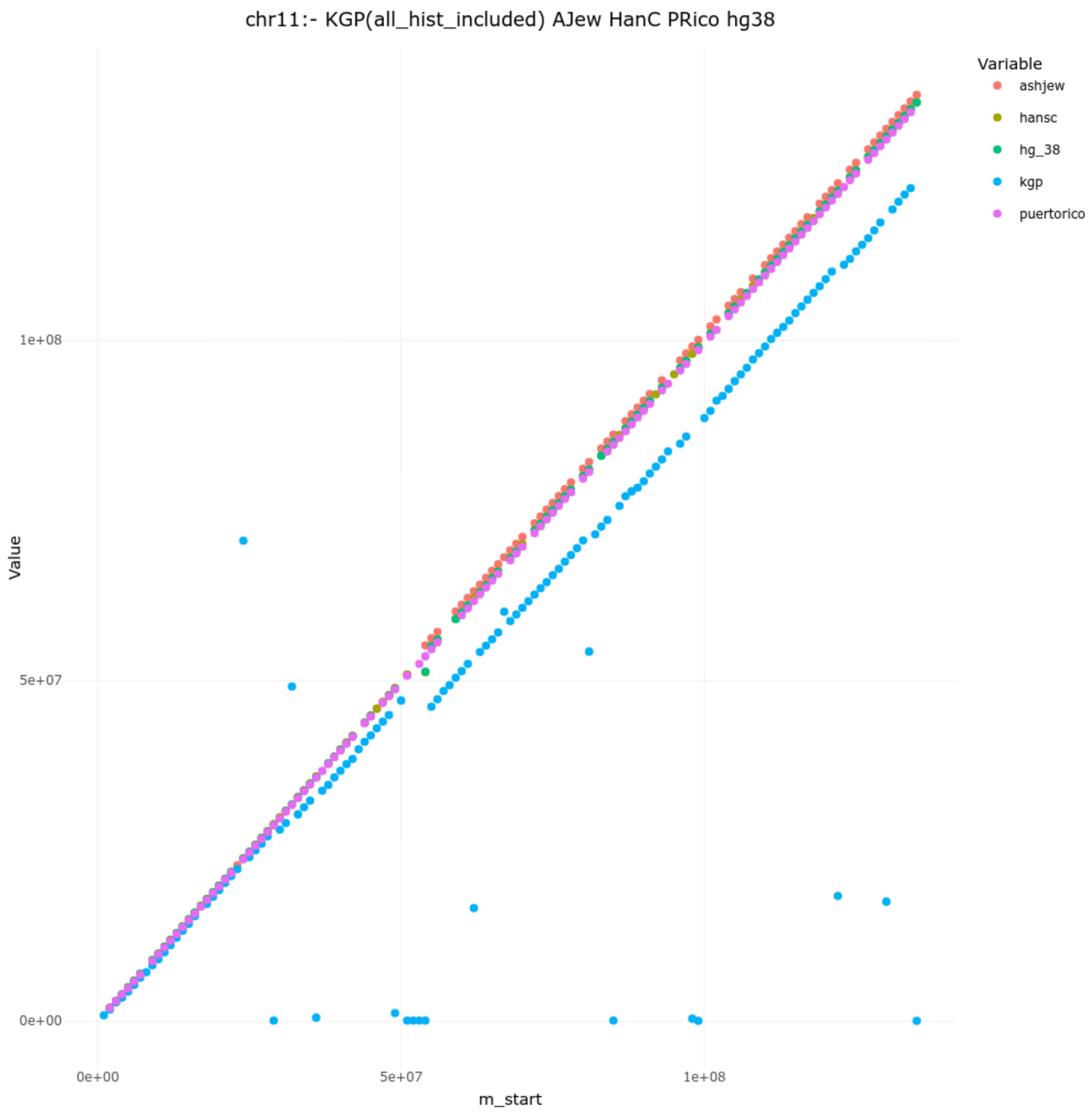

chr11:- KGP(all_hist_included) AJew HanC PRico hg38
Variable
ashjew
hansc
hg_38
kgp
puertorico
Value
1e+08
5e+07
0e+00
m_start
0e+00
5e+07
1e+08

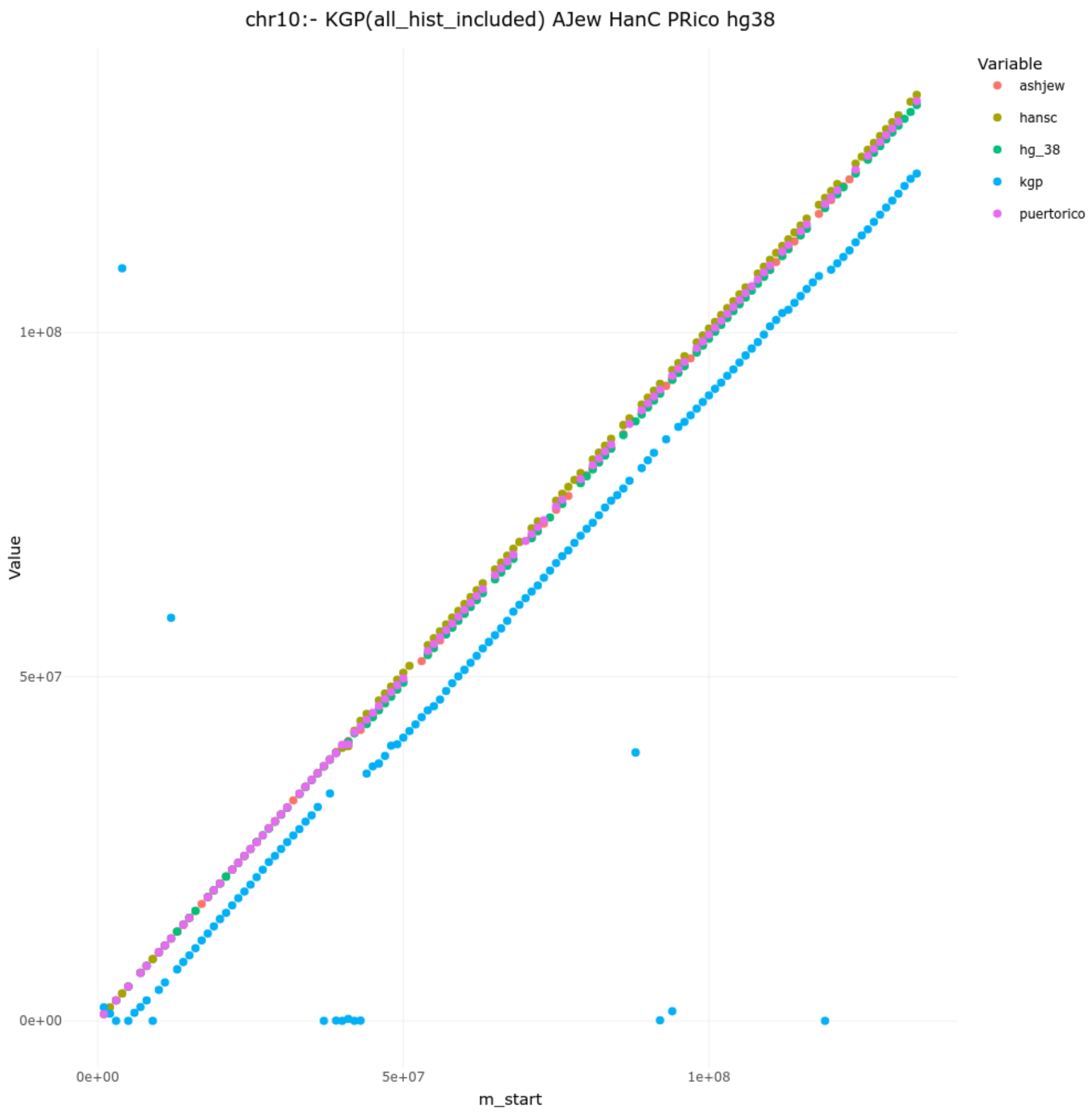
chr10:- KGP(all_hist_included) AJew HanC PRico hg38
Variable
ashjew
hansc
hg_38
kgp
puertorico
Value
1e+08
5e+07
0e+00
0e+00
5e+07
1e+08
m_start

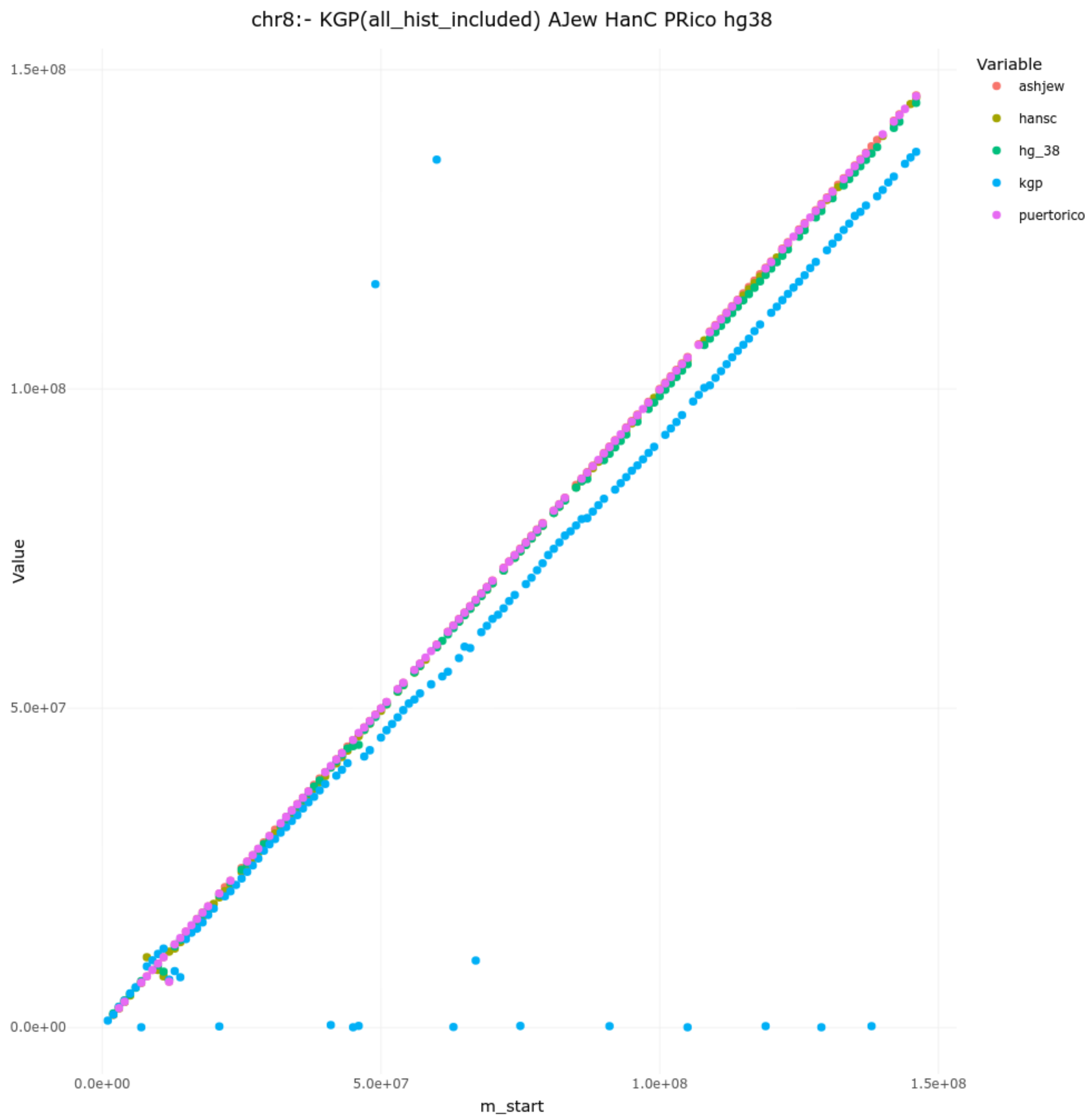

chr8:- KGP(all_hist_included) AJew HanC PRico hg38
Value
m_start
1.5e+08
1.0e+08
5.0e+07
0.0e+00
0.0e+00
5.0e+07
1.0e+08
1.5e+08
Variable
ashjew
hansc
hg_38
kgp
puertorico

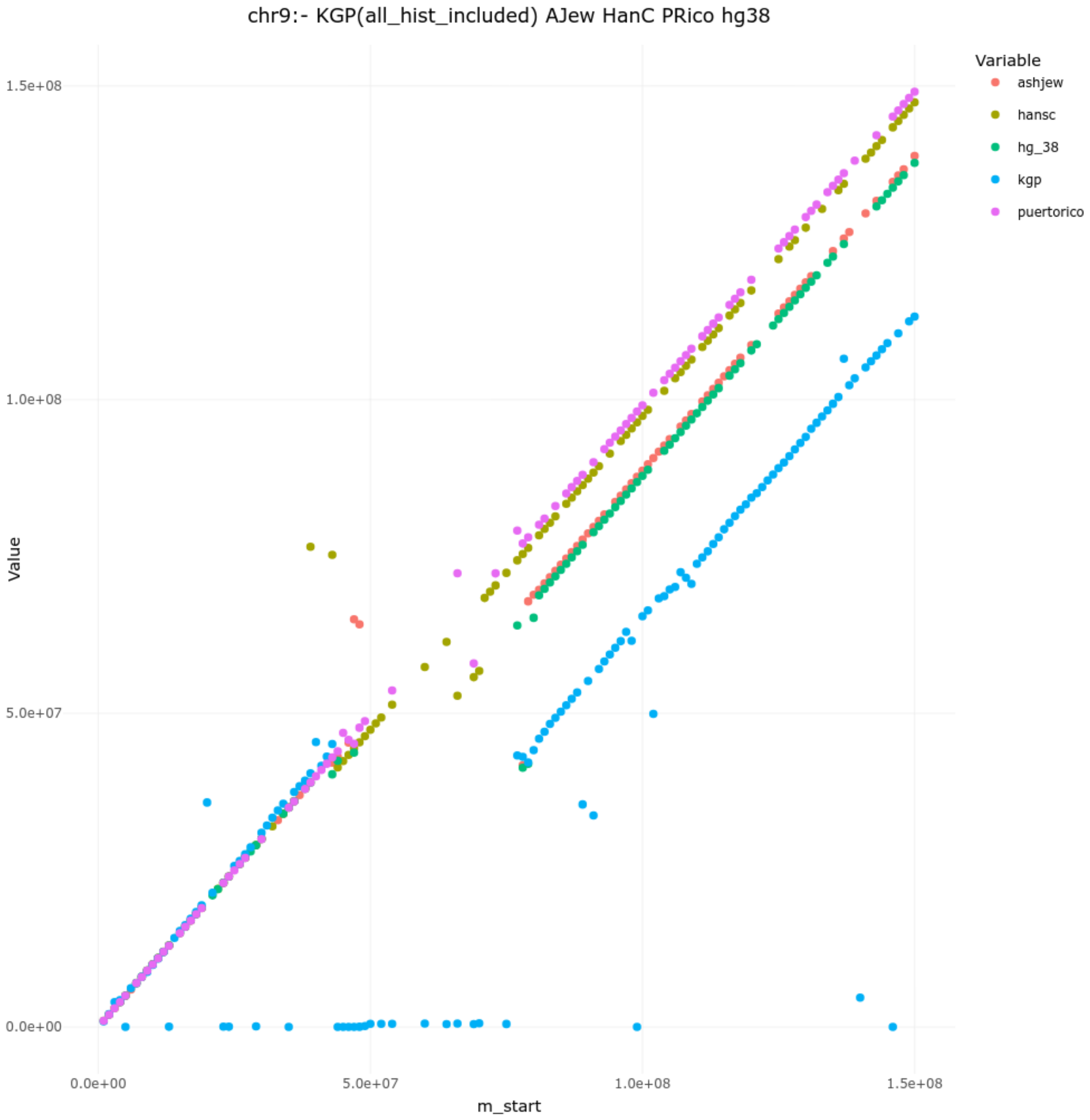

chr9:- KGP(all_hist_included) AJew HanC PRico hg38
Variable
ashjew
hansc
hg_38
kgp
puertorico
Value
1.5e+08
1.0e+08
5.0e+07
0.0e+00
0.0e+00
5.0e+07
1.0e+08
1.5e+08
m_start

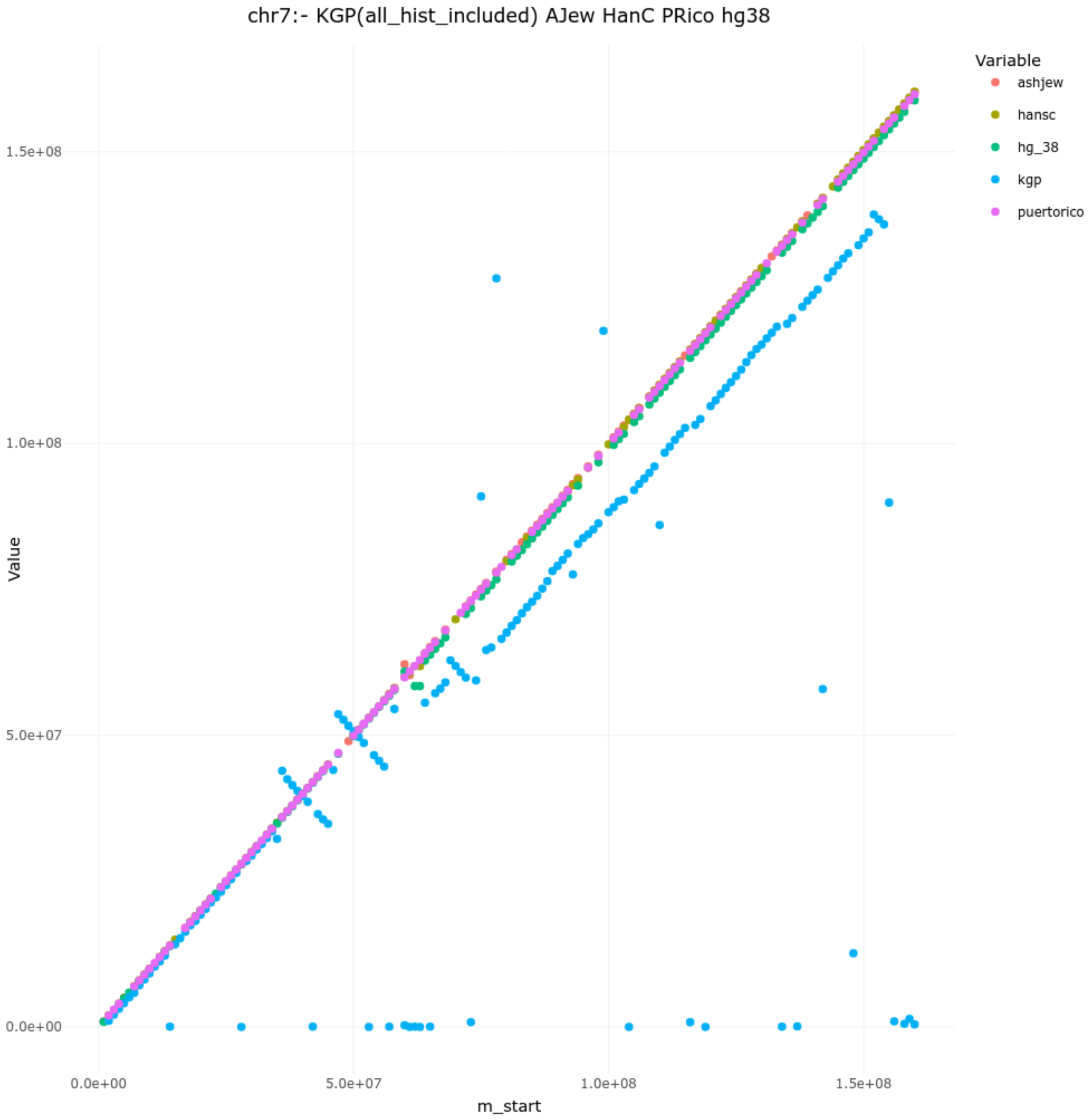
chr7:- KGP(all_hist_included) AJew HanC PRico hg38
Variable
ashjew
hansc
hg_38
kgp
puertorico
Value
1.5e+08
1.0e+08
5.0e+07
0.0e+00
0.0e+00
5.0e+07
1.0e+08
1.5e+08
m_start

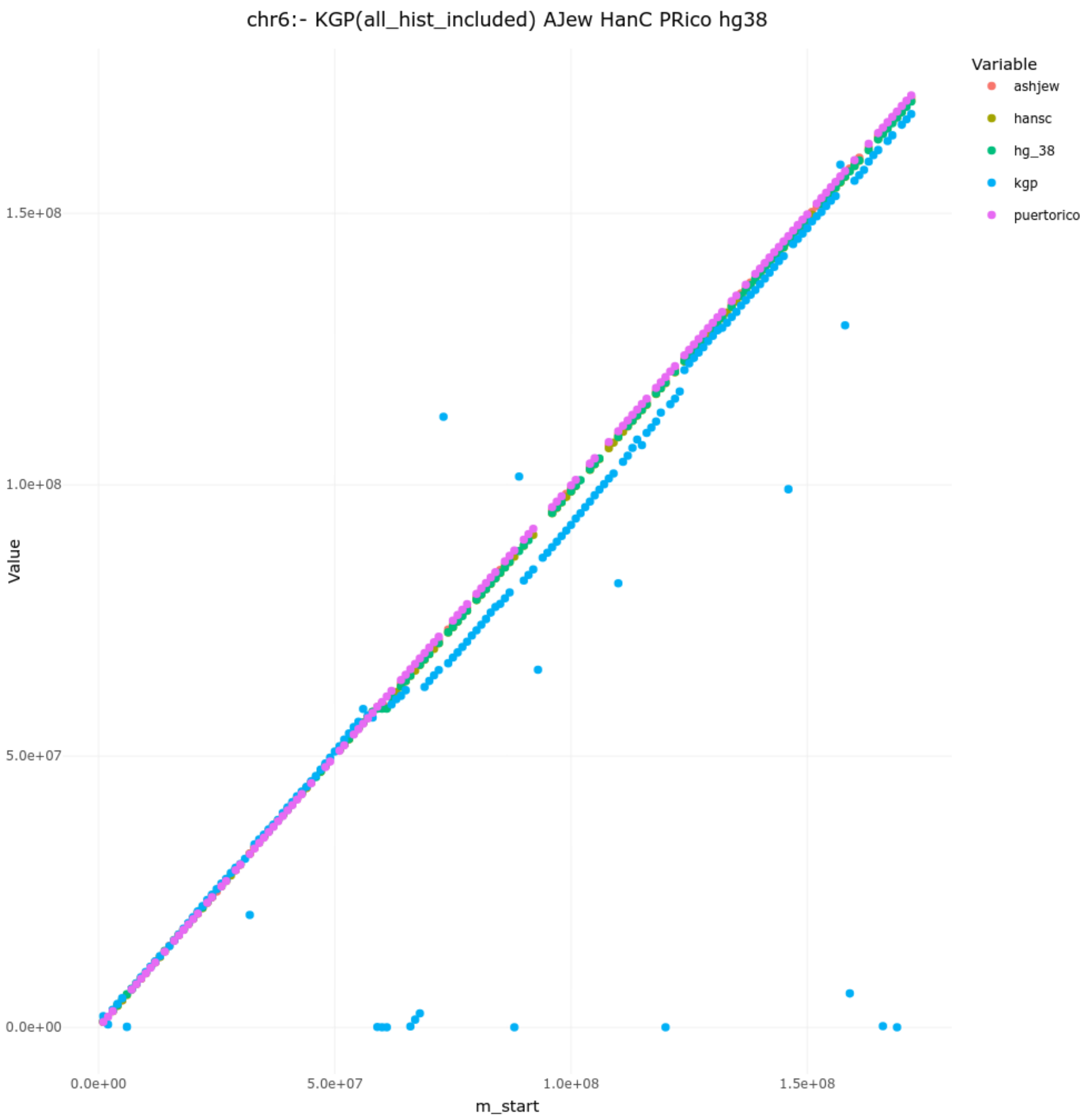

chr6:- KGP(all_hist_included) AJew HanC PRico hg38
Variable
ashjew
hansc
hg_38
kgp
puertorico
Value
1.5e+08
1.0e+08
5.0e+07
0.0e+00
0.0e+00
5.0e+07
1.0e+08
1.5e+08
m_start

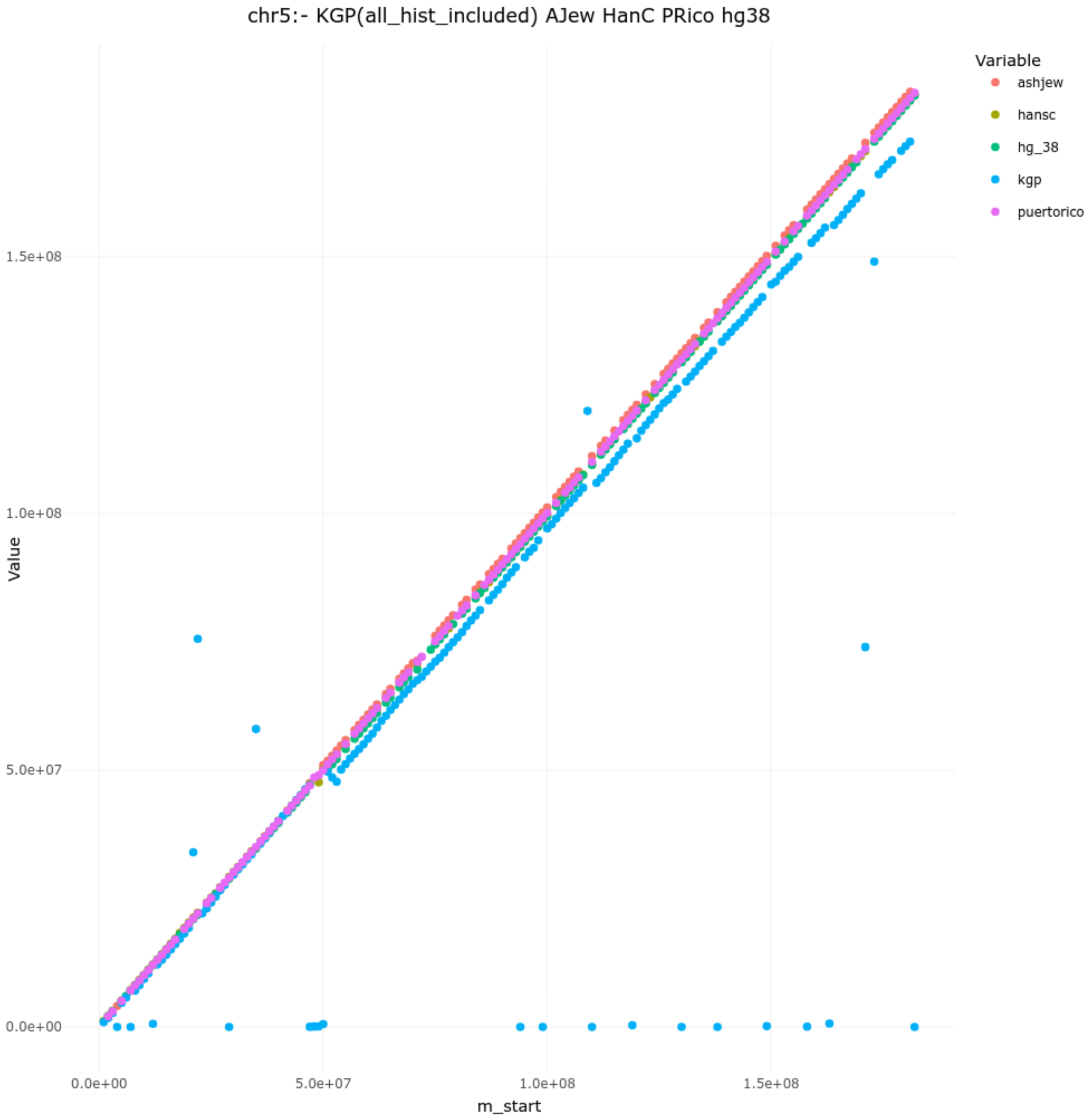

chr5:- KGP(all_hist_included) AJew HanC PRico hg38
Variable
ashjew
hansc
hg_38
kgp
puertorico
1.5e+08
1.0e+08
5.0e+07
0.0e+00
Value
0.0e+00
5.0e+07
1.0e+08
1.5e+08
m_start

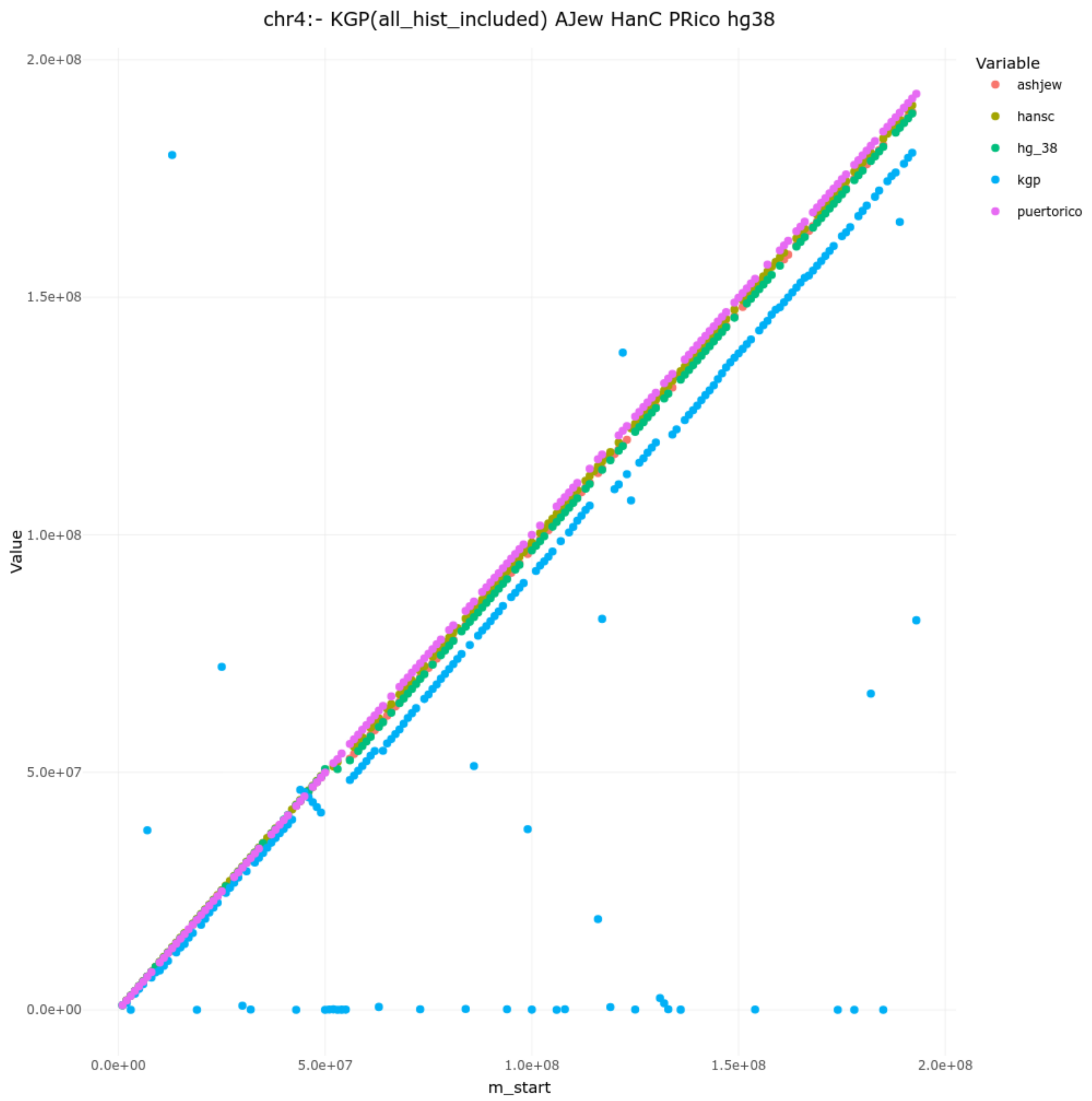
chr4:- KGP(all_hist_included) AJew HanC PRico hg38
Value
2.0e+08
1.5e+08
1.0e+08
5.0e+07
0.0e+00
0.0e+00
5.0e+07
1.0e+08
1.5e+08
2.0e+08
m_start
Variable
ashjew
hansc
hg_38
kgp
puertorico

-

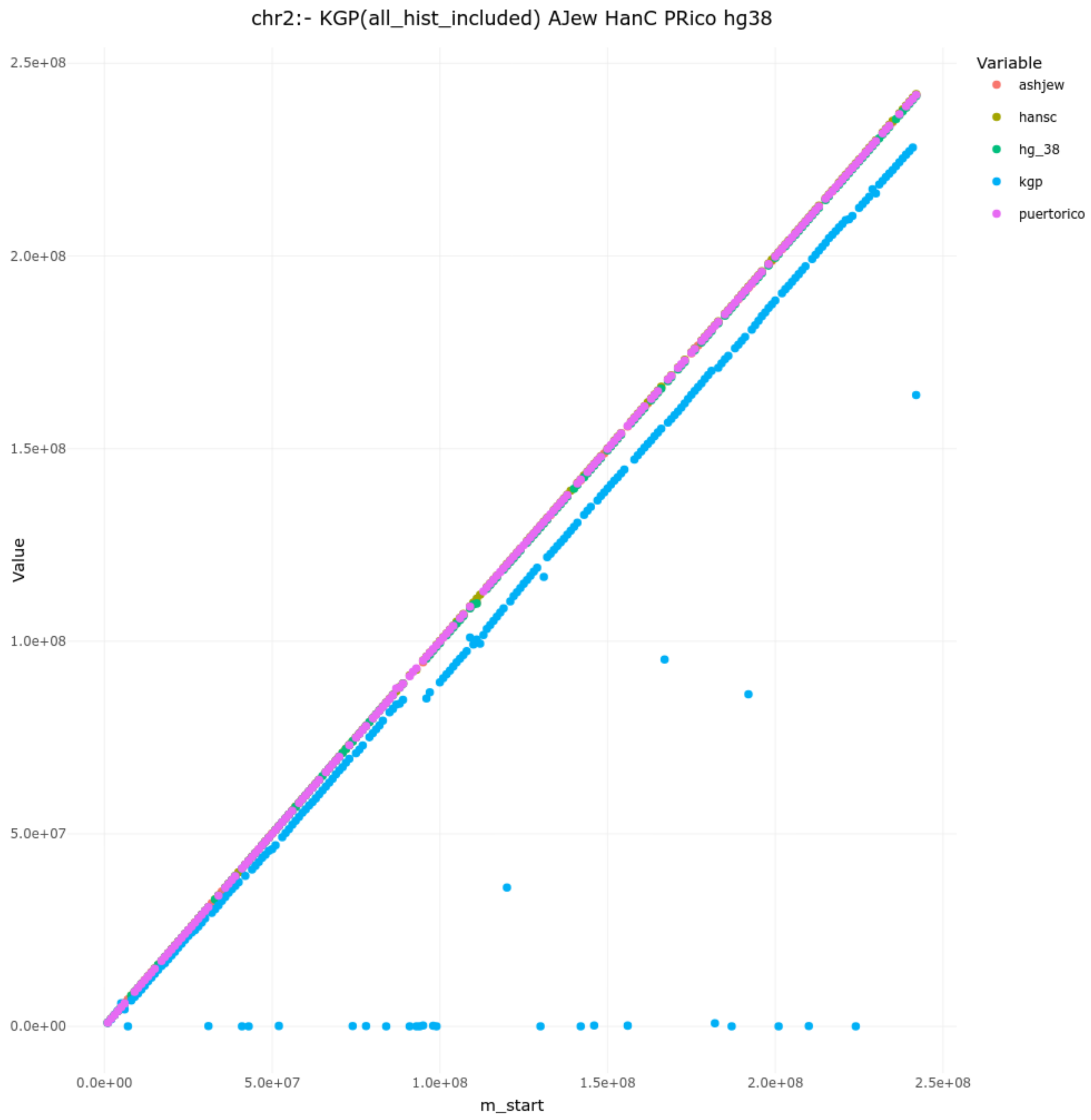
chr2:- KGP(all_hist_included) AJew HanC PRico hg38
Variable
ashjew
hansc
hg_38
kgp
puertorico
Value
2.5e+08
2.0e+08
1.5e+08
1.0e+08
5.0e+07
0.0e+00
0.0e+00
5.0e+07
1.0e+08
1.5e+08
2.0e+08
2.5e+08
m_start

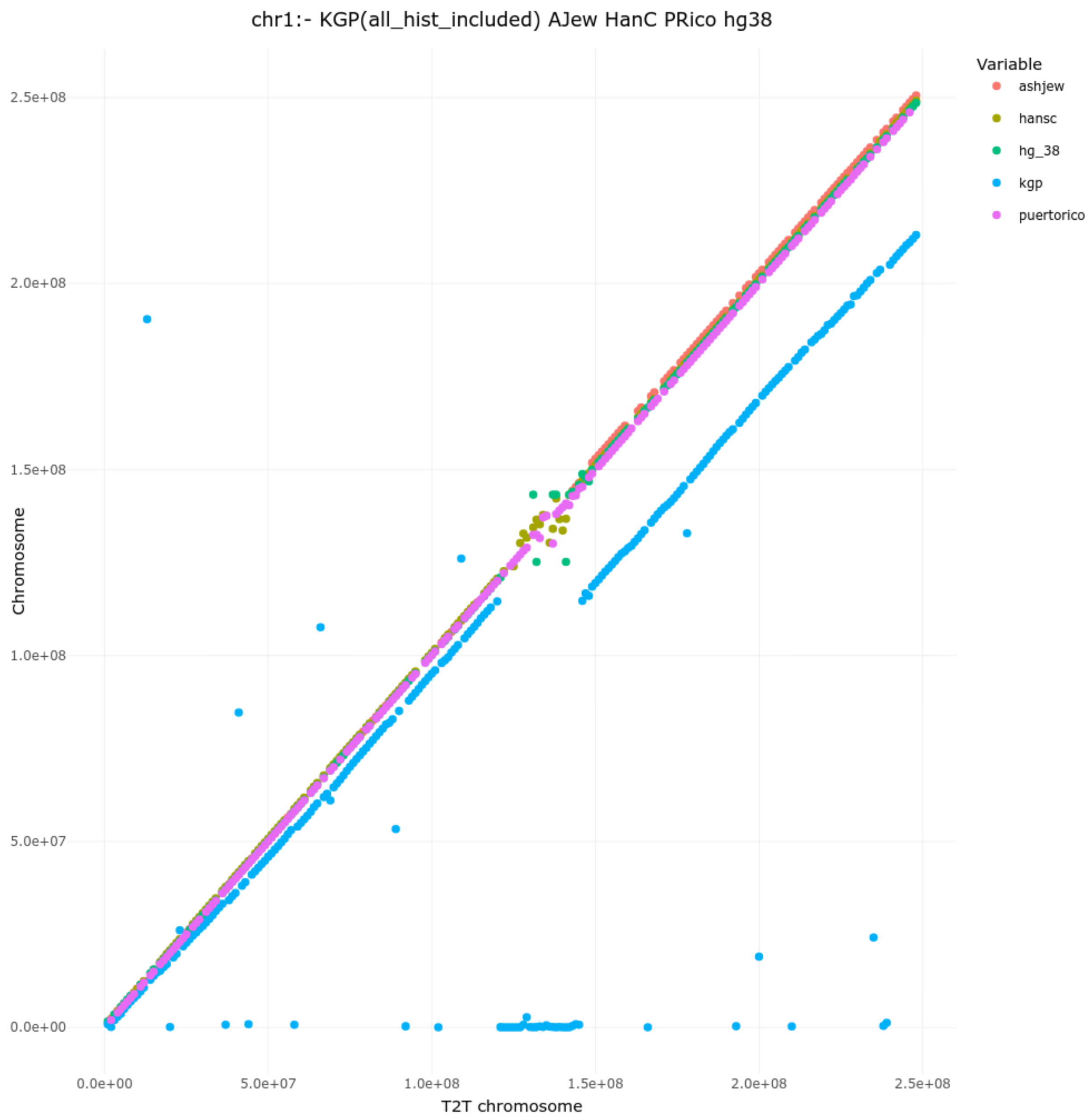
chr1:- KGP(all_hist_included) AJew HanC PRico hg38
Variable
ashjew
hansc
hg_38
kgp
puertorico
Chromosome
T2T chromosome
2.5e+08
2.0e+08
1.5e+08
1.0e+08
5.0e+07
0.0e+00
0.0e+00
5.0e+07
1.0e+08
1.5e+08
2.0e+08
2.5e+08
